# Prediction of new candidate proteins and analysis of sub-modules and protein hubs associated with seed development in rice (*Oryza sativa*) using an ensemble network-based systems biology approach

**DOI:** 10.1101/2024.11.03.621786

**Authors:** M.R.P. De Silva, J.W.J.K. Weeraman, P.C. Fernando

**Author notes:** **Contributing Authors:**.

## Abstract

**Background:** Rice is a critical global food source, but it faces challenges due to nutritional deficiencies and the pressures of a growing population. Understanding the molecular mechanisms and protein functions in rice seed development is essential to improve yield and grain quality. However, there is still a significant knowledge gap regarding the key proteins and their interactions that govern rice seed development. Protein-protein interaction (PPI) analysis is a powerful tool for studying developmental processes like seed development, though its potential in rice research is yet to be fully realized. With the aim of unravelling the protein interaction landscape associated with rice seed development, this systems biology study conducted a PPI network-based analysis. Using a list of known seed development proteins from the Gene Ontology (GO) knowledgebase and literature, novel candidate proteins for seed development were predicted using an ensemble of network-based algorithms, including majority voting (MV), Hishigaki, Functional flow, and Random Walk with Restart (RWR). The predictions were validated using enrichment analysis, and the rice seed development subnetwork was further analyzed for community and hub detection.

**Results:** The study predicted 196 new proteins linked to rice seed development and identified 14 sub-modules within the network, each representing different developmental pathways such as endosperm development and seed growth regulation. Of these, 17 proteins were identified as intra-modular hubs and 6 as inter-modular hubs. Notably, the protein SDH1 emerged as a dual hub, acting as both an intra-modular and inter-modular hub, highlighting its crucial role in coordinating interactions during rice seed development.

**Conclusions:** These findings, including the identified hub proteins and sub-modules, provide a better understanding of the PPI interaction landscape governing seed development in rice. This information is useful for improving rice cultivars for better yield and quality through genetic engineering and breeding. This study implements an ensemble of algorithms for the analysis and showcases how systems biology techniques can be applied in developmental biology.

## 1 Introduction

Rice (*Oryza sativa*), widely consumed as a staple food globally, holds a significant role in plant research being the model monocot organism. The small genome size of rice [1] has enabled in-depth exploration of various biological aspects aided by technological advancement. Rice seeds play an important role in plant growth, development, and propagation [2]. It is also a major nutrient supplier on a global scale. Therefore, grain yield and quality are significantly important for rice agriculture. The growing population has created a substantial demand for high rice yields [3]. The global population is expected to exceed 8.5 billion by 2030 [4], which requires the development of high-yielding and nutritional rice varieties as a sustainable and economical strategy.

The mechanism of rice seed development is fragile and complicated. Starting from double fertilization, it takes about 20 days to produce mature seeds [5], during which three main phases are identified: embryo morphogenesis, endosperm filling, and seed maturation. Embryo and endosperm developments are concurrently harmonized to ensure proper grain development [6]. However, a comprehensive understanding of the molecular processes underlying seed development is currently lacking [5,7]. In this context, further molecular biological investigations into rice seed development can improve the understanding of underlying biochemical pathways and help create novel high-yielding varieties.

Proteins and their interactions are particularly important in regulating developmental processes such as seed development. Typically, these developmental phenotypes involve multiple proteins interacting in intricate pathways identified as functional modules [8,9] PPI networks represent the protein interactions using computational graphs, which can be used for further analysis [10,11]. Different computational algorithms are used to analyze and unravel these networks to produce useful information. For instance, network-based candidate protein prediction algorithms are widely used to predict new protein candidates for selected molecular functions and phenotypes [9,12]. These algorithms are considered more accurate at predicting new proteins associated with development phenotypes than other sequence or structure-based protein prediction methods, such as BLAST and iterative threading assembly refinement. [12–14]. This is because developmental phenotypes involve various proteins with different sequences and structures; therefore, sequence and structure-based prediction methods may fail to predict important proteins with lower sequence and structural similarity to original proteins. Because the network-based algorithms follow a systems biology approach of using existing protein or gene interactions to predict new candidates, they capture information from proteins with different sequences and structures during the prediction. Among network-based algorithms, community detection algorithms are crucial for analyzing complex networks, such as PPI networks [15]. These algorithms aid in understanding the network organization by identifying network modules, usually associated with specific functions or phenotypes. For instance, the Louvain community detection algorithm, used in this work, is a hierarchical clustering method that uses modularity optimization to identify communities [16]. The algorithm’s robustness has established it as a dependable option for analyzing PPI networks.

Another important application of PPI network analysis is the identification of hub proteins, i.e., highly connected proteins within PPI networks. These are key components in maintaining corresponding biological pathways [13,17]. The central-lethality rule, a widely accepted concept in network biology, underscores the critical role of hubs in maintaining the network architecture in dictating biological function [18]. It stresses the pivotal role of hubs over non-hubs within the network, considering their substantial contribution to network organization. Frequently, these hubs serve as drug targets and commercially important genetic engineering and breeding targets [9,19]. Two key types of hubs stand out: inter-modular hubs, which are nodes with dense connections facilitating interactions between distinct functional modules, and intra-modular hubs, which are nodes densely interconnected within a single functional module [17].

Function-specific PPI subnetwork analysis is popular in other domains, such as cancer and vertebrate development [13,20,21]. Despite the potential, there are relatively few applications in plant development. For instance, PPI network analysis has been applied to study root development in rice and to identify genetic factors associated with parthenocarpy in bananas [22,23]. However, a PPI network analysis specifically focused on rice seed development is currently lacking. Hence, this study employed a network-based computational approach to study the protein interactome associated with rice seed development, which may uncover useful information about novel candidate proteins, sub-modules, and hubs. To our knowledge, this study represents the first comprehensive analysis of the PPI network landscape specifically associated with seed development. This involved predicting novel candidate proteins using an ensemble of network-based algorithms. The ensemble approach, a pivotal advancement in data mining and machine learning, combines multiple models into a unified entity, enhancing prediction accuracy and generalization [24]. To our knowledge, such an ensemble approach has not been used in previous network-based candidate protein predictions for biological phenotypes in plants. The next steps involved extracting the sub-network of rice seed development, identifying sub-modules within the extracted sub-network, and uncovering the hubs associated with these sub-modules. The results from this study represent newfound knowledge that paves the way for future genetic breeding and engineering experiments to enhance rice grain yield and quality. It is a valuable resource for researchers and breeders seeking to make targeted improvements in rice cultivars.

## 2 Methods

### 2.1 Data retrieval and preprocessing

A comprehensive PPI network of rice was obtained from the STRING database (version 12; September 2022; https://stringdb.org) [25]. The graph was generated using the NetworkX package (version 3.0) in Python (version 3.8) with a combined score cutoff of 0.7 [26]. To ensure data accuracy, duplicate interactions were eliminated and protein STRING IDs were converted into their preferred names before constructing the graph.

Seed proteins (proteins annotated to rice seed development) necessary to conduct the analysis were obtained through literature mining [5,27–32] and Gene Ontology (GO) search using the QuickGo tool (version 2022-09-16; September 2022; https://www.ebi.ac.uk/QuickGO/) [33]. To validate predictions, differentially expressed proteins (DEPs) associated with seed development were collected through literature mining, including studies utilizing transcriptomics experiments that analyze the complete set of RNA transcripts produced by the genome [34,35].

### 2.2 Seed development subnetwork extraction using an ensemble of network-based algorithms

An ensemble of network-based algorithms was employed to predict new seed development protein candidates [36]. To our knowledge, this study marks the pioneering use of a diverse ensemble of network-based algorithms in rice PPI network research. Four algorithms, including Majority Voting [37], Hishigaki [38], Functional Flow [39], and Network Propagation [40] were integrated using the Rule of Sum [24] to construct the ensemble. The rationale behind this approach was to lower the impact of individual algorithm biases and errors, leading to more accurate and robust predictions [41,42].

#### 2.2.1 Majority Voting (MV) algorithm

The MV algorithm, which was the first in the ensemble, calculates the prediction score by counting the number of seed proteins that are direct neighbors to non-seed proteins [43]. To perform this calculation Eq. (1) was used, where for a protein with *n* neighbors, *xi* represents whether neighbor *i* is a seed protein (x=1), or not (x=0). However, a notable limitation of this approach is its exclusive focus on immediate neighbors. Also, it does not make the best use of the overall network structure, and it is biased towards highly annotated functions as it relies on the frequency of annotations to predict the function of proteins.

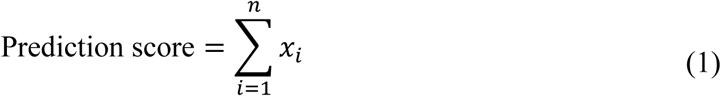

#### 2.2.2 Hishigaki algorithm

The Hishigaki algorithm [38], the second in the ensemble, utilizes Eq. (2) to compute the prediction score. This algorithm extends the MV concept in predicting protein functions by analyzing proteins within a specified radius. It utilizes the chi-squared test to mitigate the bias for overrepresented functional annotations, unlike MV.

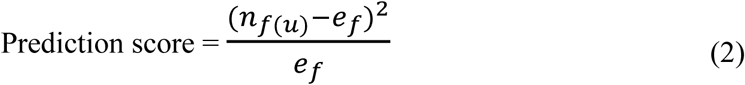

In Eq. (2), *n*_*f*(*u*)_ represents the count of proteins with a specific function (f) within the n-neighborhood of the protein “u”. Additionally, *e*_*f*_ stands for the expected frequency for the particular function, which is calculated using Eq. (3) below.

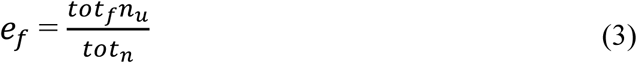

In Eq. (3), *tot*_*f*_ represents the total count of proteins with a particular function in the entire network, *tot*_*n*_ indicates the total number of proteins in the entire network, and *n*_*u*_ stands for the total number of neighbors for the protein “*u*”.

#### 2.2.3 Functional flow algorithm

The functional flow algorithm, the third algorithm in the ensemble, broadens the concept of guilt by association with protein groups, regardless of whether they physically interact with each other [44] This algorithm computes the prediction score for each non-seed protein by following a set of rules iteratively, corresponding to half the network diameter.

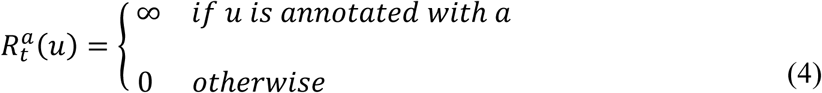

Eq. (4) defines *R*^*a*^_t_(*u*) as the fluid volume of node u for function “a” at time t. Only the nodes corresponding to seed proteins have infinite fluid reserves of function “a” at time 0. Considering the amount of flow received in and exited out of each node, the reservoir of each node is recomputed for each successive time step according to equation 5.

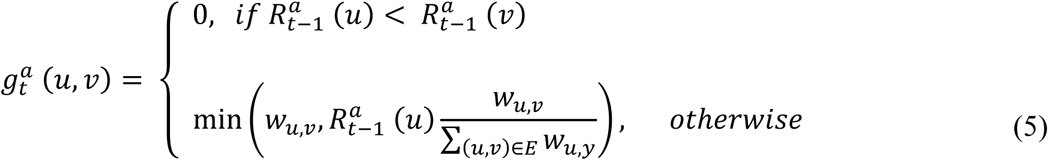

In Eq. (5), *g*^*a*^_t_ (*u*, *v*) stands for the flow received by protein v from protein u for the function “a” at time t, *W*_*u*,*v*_ is the weight of the edge between u and v proteins, *R*^*a*^ (*u*) is the reservoir of u at time t-1 for the function “a”, *R*^*a*^ (*v*) is the reservoir of v at time t-1 for the function “a”, and ∑_(*u*,*v*)∈*E*_ *W*_*u*,*y*_ is the sum of weights of edges connecting the protein u. There is no flow between nodes at time 0. Always the flow is downhill adhering to the capacity constraints in equation 5. The maximum flow allowed between two proteins is corresponding to the weight of the connecting edge.

At the end of each iteration, the reservoir at each node is finalized using Eq. (6).

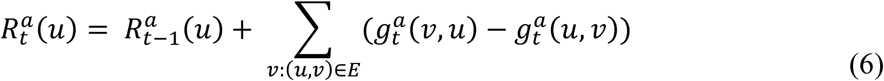

The term ∑_*v*:(*u*,*v*)∈*E*_(*g*^*a*^(*v*, *u*) − *g*^*a*^(*u*, *v*)) of equation 6 represents the net flow change due to each immediate neighbor of the protein. At the end of defined iterations, each protein is annotated with the functional score calculated as the total amount of flow received by the protein. Seven iterations were required to compute the functional score, which is half the network diameter.

#### 2.2.4 Random Walk with Restart

The fourth algorithm of the ensemble was Random Walk with Restart (RWR). This method estimates the likelihood of a “walk” along random edges connecting various nodes, ultimately reaching a specific target node from a predefined set of starting nodes [40]. The random walk length is determined by the number of iterations. The restart parameter, also known as the learning parameter, regulates the probability of the random walk “jumping” back to its starting position mid-walk. The resulting node weights represent the probability of the random walk landing on them when initiated from any of the seed nodes. RWR in matrix form is displayed in Eq. (7) below.

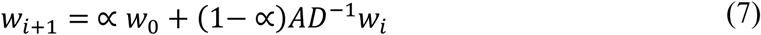

In this context, w0 signifies the initial weight vector, wi represents the weights after the i^th^ iteration, wi+1 denotes the weights after the (i+1)^th^ iteration, and α represents the learning parameter for the algorithm. Additionally, A and D refer to the adjacency and degree matrices of the PPI network graph, respectively.

Here, the initial weight of corresponding nodes in the PPI network was set using the seed proteins list, ensuring a weight of 100 to prioritize the random walker’s inclination to visit these seed nodes. Network propagation underwent 5 iterations with a learning parameter (α) of 0.1, focusing on local interactions [45]. The iteration count was capped at 5 to include relevant neighborhoods containing other seed development-related proteins and prevent exclusion due to longer paths [46].

### 2.3 Total prediction score using the rule of sum

To build the ensemble model following the calculation of prediction scores for non-seed proteins using the four selected algorithms, the scores were normalized using the min-max normalization method and were compared with the list of DEPs obtained through literature mining to calculate the validation score by DEPs [35,47]. A non-seed protein was given a score of 1 if it was a DEP; otherwise, it was assigned a score of 0. The total prediction score for each non-seed protein was obtained by adding the normalized prediction scores from the four algorithms and the DEP validation score using the rule of sum as shown in Eq. (8) below [24].

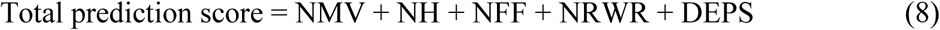

Eq. (8) represents the calculation of the total prediction score for each non-seed protein, where NMV denotes the normalized majority vote score, NH denotes the normalized Hishigaki score, NFF denotes the normalized functional flow score, and NRWR denotes the normalized RWR score.

### 2.4 Optimal cutoff for top candidate selection

Initially, the resulting total prediction scores for non-seed proteins were sorted in descending order to identify the top candidates. Then, to determine the optimal combined score cutoff for selecting the top-ranked candidates, the precision top-N curve [48] method was adapted and modified. This modified curve plotted the precision at the N^th^ position (the percentage of DEPs within the top N predictions which is calculated using Eq. (9) against the number of top predictions (N)).

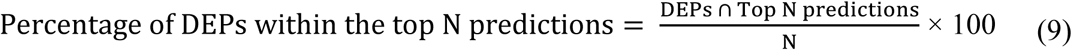

In Eq. (9), the percentage of DEPs within the top N predictions was calculated by plugging in the appropriate values for N and calculating the overlap between the list of DEPs and the top N predictions. The optimal cutoff was determined by selecting the value of N that corresponds to the highest precision.

The seed development subnetwork was then extracted, which included the seed proteins and candidates filtered using the optimal cutoff and their interactions. Only the proteins that are also DEPS within the selected top N candidates were included to reduce false positive predictions.

### 2.5 Validation of the predictions

For computational validation of predictions, the predicted protein candidates were subjected to enrichment analysis using the functional annotation tool in the Database for Annotation, Visualization and Integrated Discovery (DAVID) web application (Version: DAVID 2021; December 2022; https://david.ncifcrf.gov/tools.jsp) [49]. The Biological Process component of Gene Ontology (GO-BP) was used and terms with a p-value below 0.05 were selected [50].

### 2.6 Identifying sub-modules related to seed development

The Louvain community detection algorithm was used to perform sub-module analysis on the seed development subnetwork [16]. This algorithm has been widely applied in biological network studies, demonstrating promising outcomes [51]. The community subpackage in NetworkX (version 3.0) was used to implement the Louvain algorithm. Following the sub-module identification process, isolated sub-modules that consisted of proteins less than 6 were excluded from further analysis [52]. Subsequently, functional enrichment analysis for each detected sub-module was performed using the DAVID functional annotation tool (Version: DAVID 2021) [49]. This analysis aimed to annotate each sub-module with the most relevant GO-BP term associated with seed development, with a p-value less than 0.05.

### 2.7 Detection and analysis of hub proteins

To detect intra-modular hubs, which are densely connected proteins within a sub-module, a Z-score for each node was calculated [53] using Eq. (10).

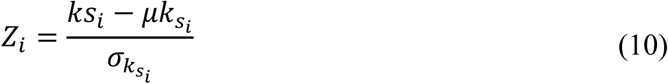

The Z-score for node i is represented by *Z*_*i*_ in Eq. (10), while *ks*_*i*_ denotes the within-modular degree (number of interactions) of node i within the module *s*_*i*_. Moreover, *μk*_*si*_ and σ_*ks*_ represent the mean and standard deviation of within-module degree in the module *s*_*i*_, respectively. Nodes having Z-scores greater than or equal to 1.5 were identified as intra-modular hubs [54].

To identify inter-modular hubs, which are proteins with dense connections linking several different sub-modules, a Partition Coefficient (PC) for each node was calculated using the following formula [55].

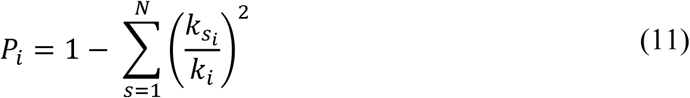

In Eq. (11), *k*_*i*_ represents the network degree of node i, N is the total number of modules, and *k*_*si*_ is the within-modular degree of node i within the module s. The PC value varies between 0 and 1, reflecting a protein’s involvement in either intra-modular or inter-modular interactions. A PC value closer to 0 signifies a higher proportion of intra-modular connections, while a PC closer to 1 indicates a higher proportion of inter-modular connections over intra-modular connections. Inter-modular hubs were selected based on a PC value greater than 0.5 [56].

Finally, structural and functional analyses were performed for each identified hub to further unravel useful information. Through the use of web-based tools for sequence analysis, the family, superfamily, and domains present in the predicted hubs were identified. Additionally, de novo structures predicted for the candidates were searched in the AlphaFold database [57,58]. The web tools used, their corresponding URLs and versions, and the type of analysis performed are listed in Table 1.

**Table 1.** Web tools utilized for candidate hub analysis.

| Web Tool | Web URL | Web Version | Type of analysis |
| --- | --- | --- | --- |
| InterProScan | https://www.ebi.ac.uk/interpro/ | 93.0 | Protein family |
| **Pfam** | https://www.ebi.ac.uk/interpro/entry/pfam/ | 35.0 | Protein family and domains |
| **Panther** | http://www.pantherdb.org/ | 17.0 | Protein family and class |
| **ScanProsite** | https://prosite.expasy.org/scanprosite/ | 2023_01 | Protein domains |
| **Supfam** | https://supfam.org/ | 1.75 | Protein family |
| **SMART** | https://smart.embl.de/ | 9.0 | Protein domain |

### 2.8 The bioinformatics pipeline

The bioinformatics pipeline used for this analysis is illustrated in Fig. 1. This pipeline was constructed in Python 3.8, using NetworkX (version 3.0) and community Python packages. For network visualizations, the Cytoscape software (version 3.9.0) was used. All the data and Python codes can be accessed at: https://github.com/rashpr88/RiceSeedDevelopment.

**Fig. 1.**
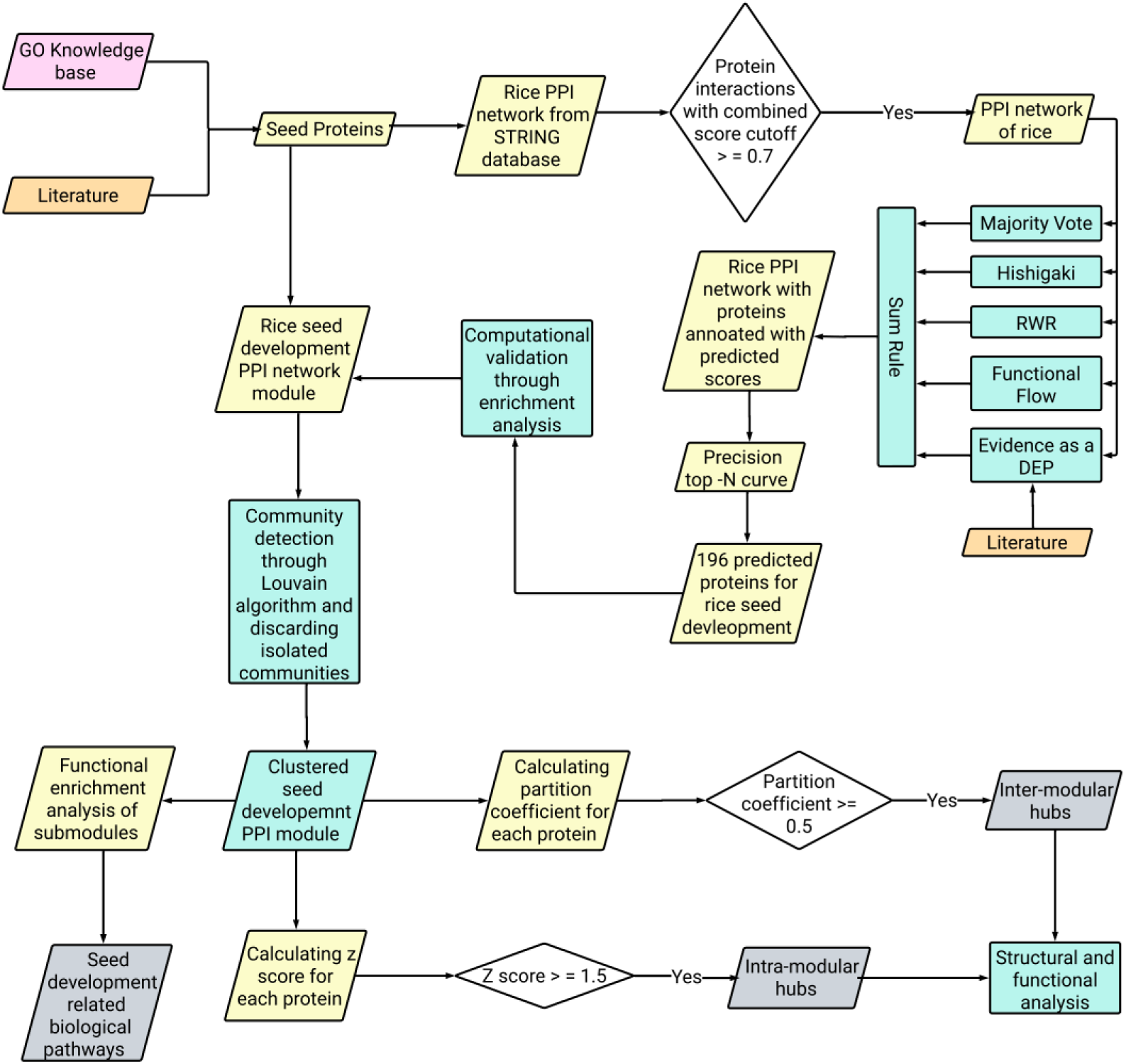
The bioinformatics pipeline used for prediction and validation of protein candidates for seed development in rice. The hub protein analysis procedure and sub-pathway analysis procedure are also represented.

## 3 Results

### 3.1 Data preprocessing

The global PPI network of rice is an undirected graph consisting of 25, 266 nodes and 7, 767, 680 interactions. From this interactome, 102 seed proteins associated with seed development were identified (Additional file 1: Table S1). Additionally, 640 DEPs linked to seed development were retrieved from literature mining (Additional file 1: Table S2). Filtering the PPI network with a combined score cutoff of 0.7 led to the inclusion of 95 out of the 102 seed proteins (Additional file 1: Table S1) and 285 out of the 640 DEPs. These filtered proteins were used for the predictions. The PPI network used to obtain these predictions is depicted below in Fig. 2.

**Fig. 2.**
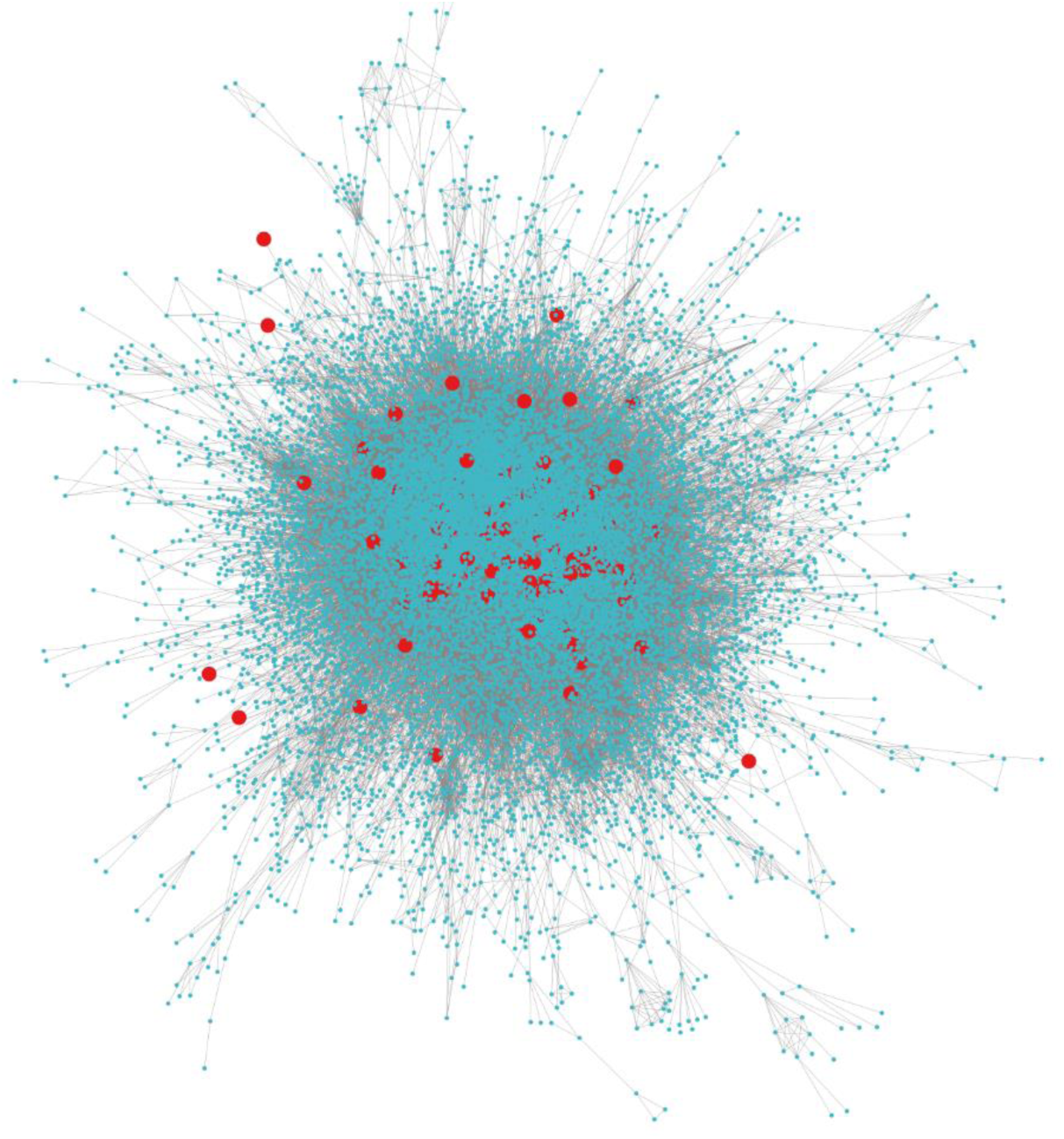
The PPI network of rice based on the 0.7 combined score cutoff. In the visualization, large seed nodes are colored in red, while small non-seeds are colored in blue.

### 3.1 Seed development sub-network extraction

After using the ensemble of network-based algorithms for predicting novel candidates for seed development, a cutoff had to be applied to select the best candidates. Fig. 3 illustrates the modified precision top-N curve generated to determine the cutoff for selecting the top predictions for further analysis.

**Fig. 3.**
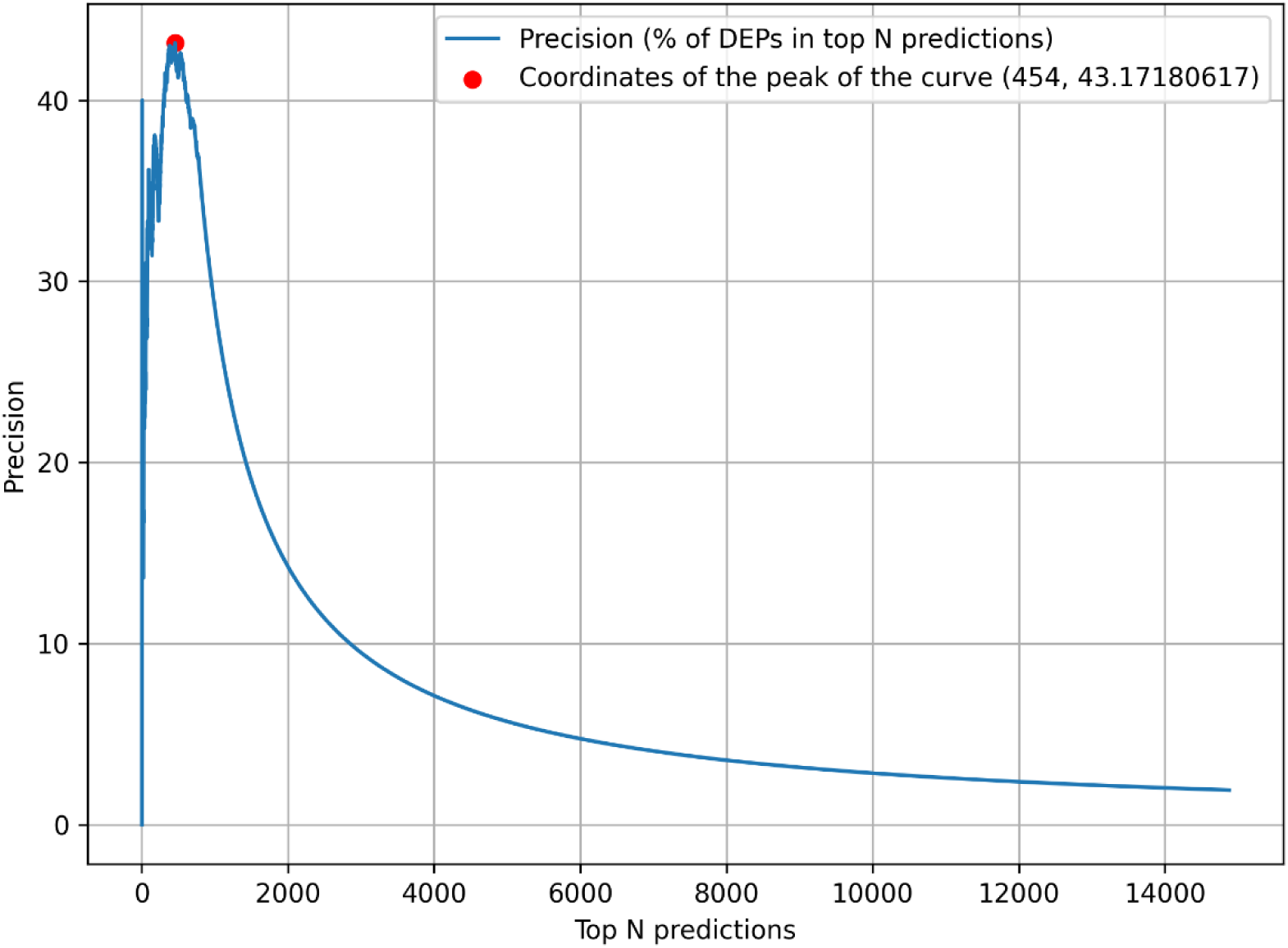
The modified precision top-N curve for predictions obtained using the rice PPI network.

Based on the curve analysis, the highest precision of 43.17% was achieved at 454 top-predicted proteins, which was chosen as the optimal cutoff. This resulted in 196 DEPs being selected for further analysis.

### 3.2 Computational validation of the predicted protein candidates

Table 2 presents the top 10 GO-BP terms that were enriched for the predicted protein candidates linked to seed development. The GO-BP term “organic hydroxy compound biosynthetic process” (GO:1901617) is the top ranked term. Seed development is a complex process that involves multiple biosynthetic pathways [59]. Many hydroxy compounds are known to play important roles in seed development. Therefore, the fact that the “organic hydroxy compound biosynthetic process” exhibits the highest level of enrichment underscores the importance of the predicted protein candidates in seed development.

**Table 2.**
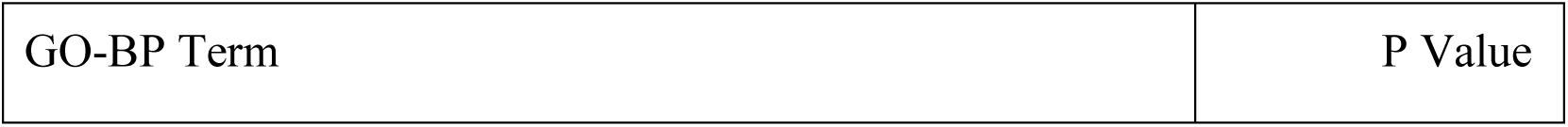
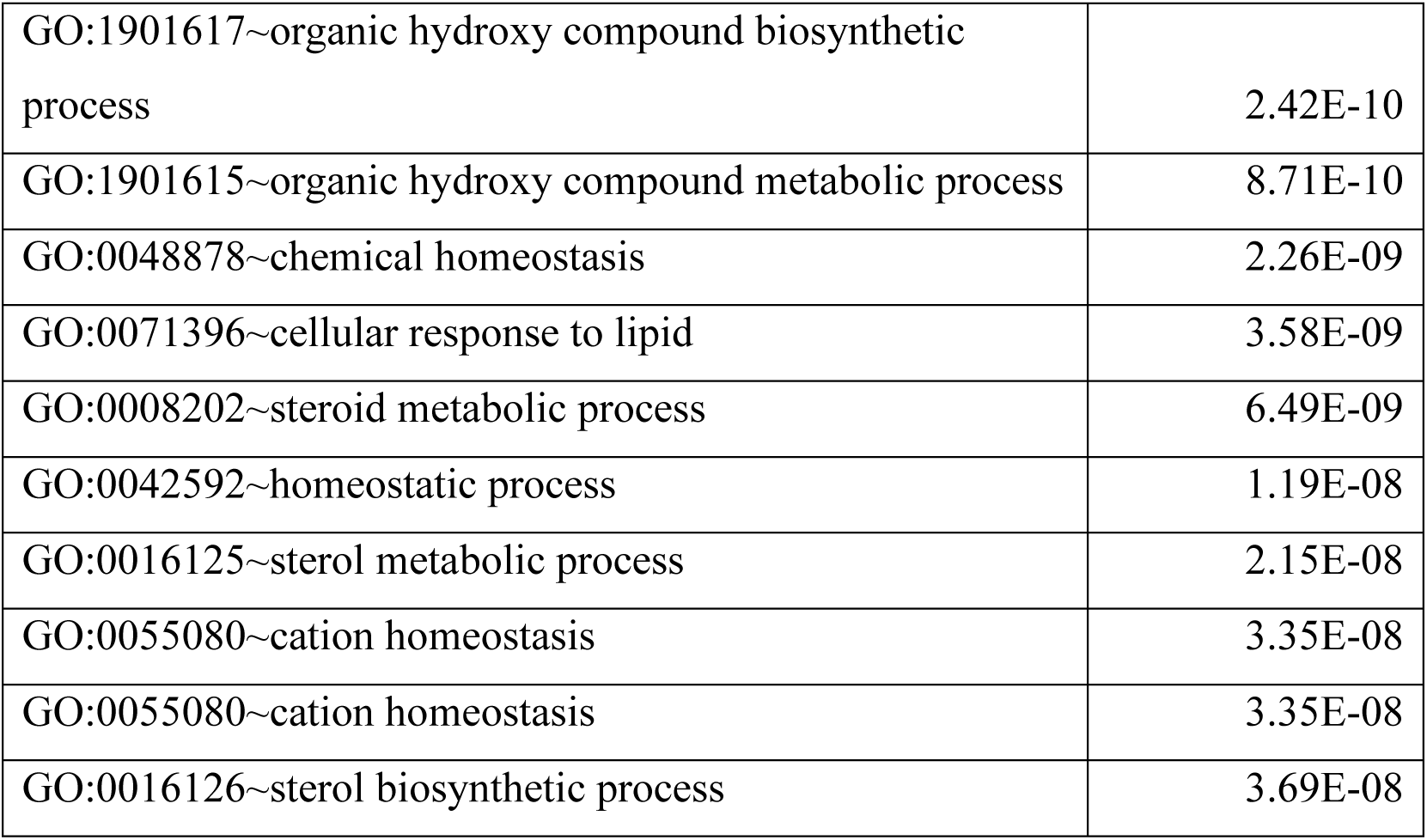
Enriched GO-BP terms for predicted proteins related to seed development in rice.

The final seed development subnetwork contained a total of 291 proteins, including 95 seed proteins directly annotated to seed development from literature, and 196 proteins predicted by the ensemble of network-based algorithms and also validated as DEPs.

### 3.3 Sub-modules related to seed development

After the module partitioning process of the extracted network conducted by the Louvain algorithm, modules with more than five nodes were kept [52]. This resulted in fourteen sub-modules, which are depicted in Fig. 4.

**Fig. 4.**
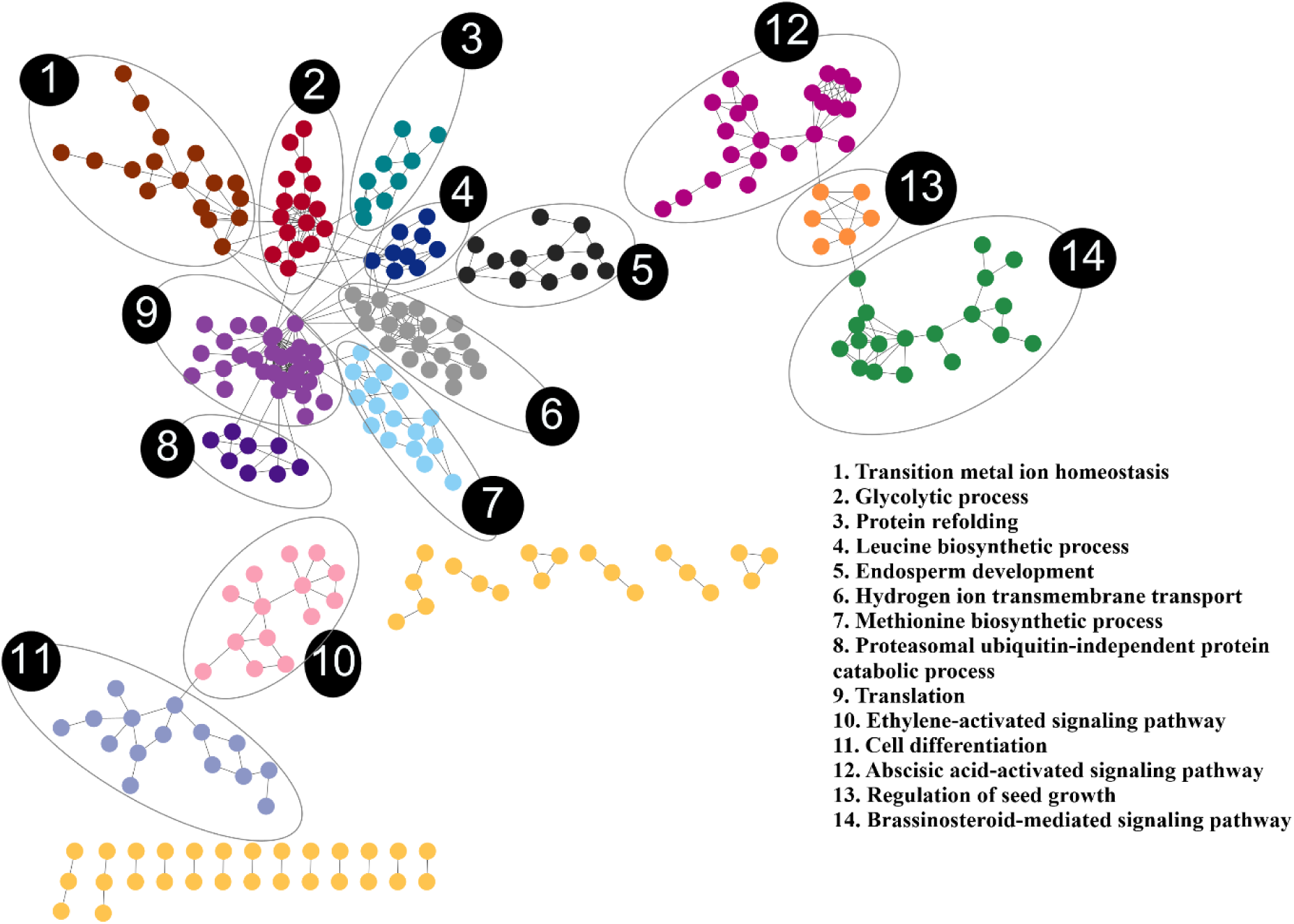
A visual representation of the seed development PPI sub-network of rice, with different colors assigned to the detected sub-modules, based on the Louvain community detection algorithm. The sub-modules are numbered and the most relevant GO-BP term related to seed development according to enrichment analysis results are listed in the legend.

Additional file 1: Table S3 presents the results of the functional enrichment analysis for sub-modules, including only the GO terms with a p-value below 0.05. Each sub-module is associated with the most relevant GO-BP term related to seed development, which is listed in Table 3 below.

**Table 3.** Most relevant GO-BP terms assigned to each sub-module of the seed development sub-network of rice.

| Sub-module | Term Identifier | Enriched GO-BP Term |
| --- | --- | --- |
| 1 | GO:0055076 | Transition metal ion homeostasis |
| 2 | GO:0006096 | Glycolytic process |
| 3 | GO:0042026 | Protein refolding |
| 4 | GO:0009098 | Leucine biosynthetic process |
| 5 | GO:0009960 | Endosperm development |
| 6 | GO:1902600 | Hydrogen ion transmembrane transport |
| 7 | GO:0009086 | Methionine biosynthetic process |
| 8 | GO:0010499 | Proteasomal ubiquitin-independent protein catabolic process |
| 9 | GO:0006412 | Translation |
| 10 | GO:0055076 | Ethylene-activated signaling pathway |
| 11 | GO:0030154 | Cell differentiation |
| 12 | GO:0009738 | Abscisic acid-activated signaling pathway |
| 13 | GO:0080113 | Regulation of seed growth |
| 14 | GO:0016126 | Brassinosteroid-mediated signaling pathway |

Additional file 1: Table S4 provides statistical information regarding the total number of proteins, seed proteins, predicted proteins, and the most relevant enriched GO-BP term for each of the 14 analyzed sub-modules.

When considering the enrichment results, certain sub-modules exhibit significant enrichment in seed development-specific pathways, such as endosperm development (sub-module 5), and regulation of seed growth (sub-module 13), while others are enriched for more general pathways, such as translation, cell differentiation, and glycolytic process, associated with seed development.

### 3.4 Hub analysis

Hub analysis identified 17 intra-modular hubs and 6 inter-modular hubs. Fig. 5 illustrates the hubs detected in the seed development sub-network.

**Fig. 5.**
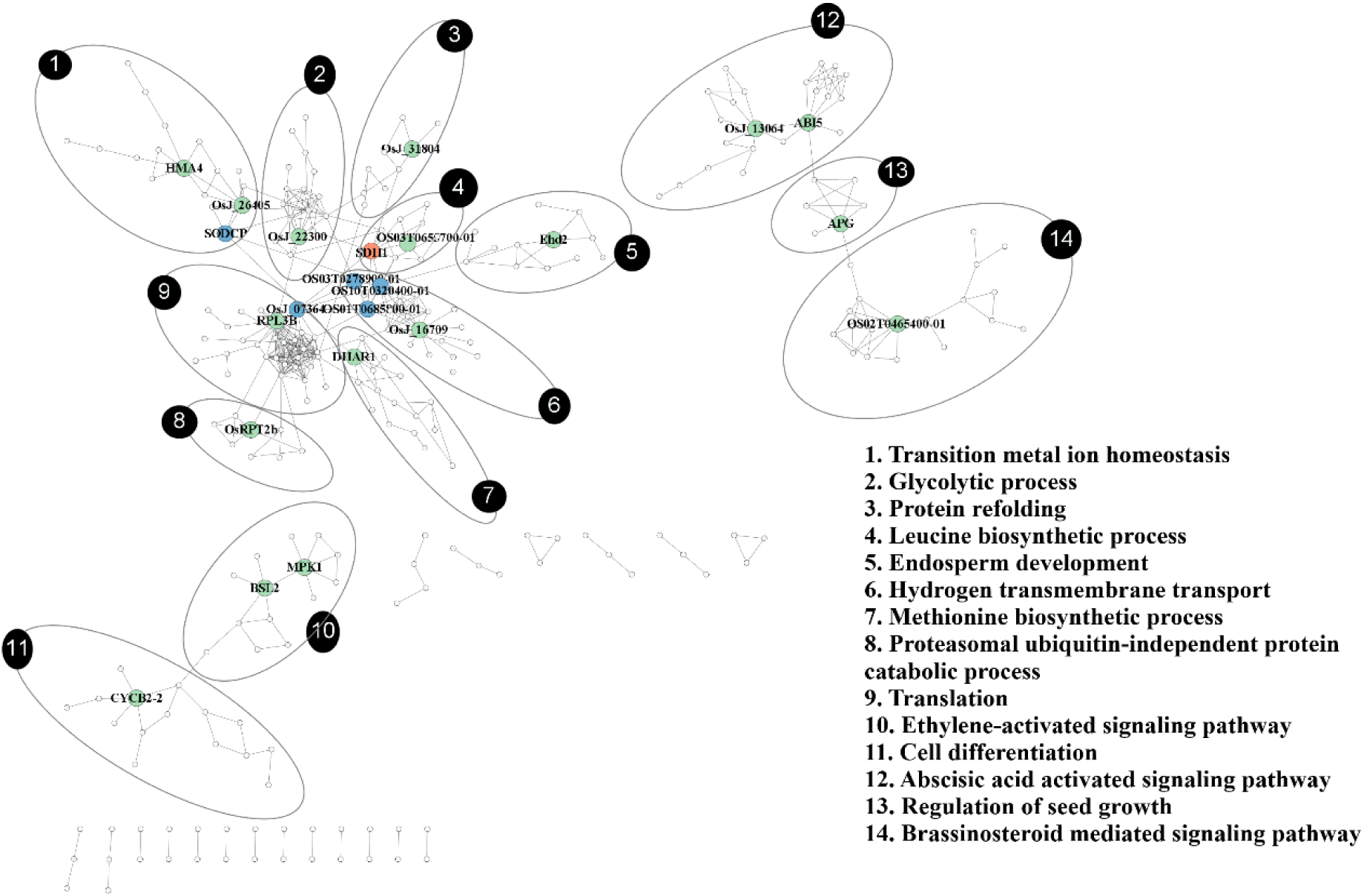
Illustration of detected hubs in the seed development PPI sub-network of rice. Larger green-colored nodes represent intramodular hubs, whereas larger blue nodes represent inter-modular hubs. Furthermore, larger orange nodes indicate proteins with dual roles as inter-modular and intra-modular hubs and small white-colored nodes are non-hubs. The sub-modules are numbered and the most relevant GO-BP term related to seed development according to enrichment analysis results are listed in the legend.

### 3.5 Intra-modular hubs

Table 4 presents the 17 intra-modular hubs detected based on Z-score values. There were 9 seed proteins and 8 predicted proteins among the intra-modular hubs. These predicted proteins are also DEPs associated with seed development according to literature but do not have any other experimental evidence for seed development. They serve as excellent candidates for further wet lab studies related to seed development.

**Table 4.** Identified intra-modular hub proteins along with their respective network degree, module degree, Z scores, the status as either a seed protein or a prediction, and their associated sub-module within the seed development sub-network.

| Node | Network degree | Module degree | Z score | Status | Sub-module |
| --- | --- | --- | --- | --- | --- |
| DHAR1 | 7 | 6 | 2.745626 | Seed | 7 |
| OsJ_16709 | 14 | 14 | 2.658782 | Seed | 6 |
| CYCB2-2 | 5 | 5 | 2.636499 | Seed | 11 |
| HMA4 | 7 | 7 | 2.569458 | Predicted protein | 1 |
| OsJ_26405 | 7 | 7 | 2.569458 | Predicted protein | 1 |
| ABI5 | 8 | 7 | 2.366432 | Predicted protein | 12 |
| OsJ_13064 | 7 | 7 | 2.366432 | Predicted protein | 12 |
| MPK1 | 6 | 6 | 2.311251 | Seed | 10 |
| OS02T0465400-01 | 7 | 7 | 2.227515 | Predicted protein | 14 |
| RPL3B | 22 | 21 | 2.122096 | Predicted protein | 9 |
| OsRPT2b | 6 | 5 | 2.064742 | Predicted protein | 8 |
| APG | 6 | 5 | 1.732051 | Seed | 13 |
| OsJ_31804 | 4 | 4 | 1.725324 | Seed | 3 |
| BSL2 | 5 | 5 | 1.664101 | Predicted protein | 10 |
| OS03T0655700-01 | 5 | 5 | 1.581139 | Seed | 4 |
| SDH1 | 10 | 5 | 1.581139 | Seed | 4 |
| OsJ_22300 | 11 | 10 | 1.572632 | Seed | 2 |

### 3.6 Inter-modular hubs

Table 5 shows the 6 inter-modular hub nodes identified based on PC calculation. This included 4 seed proteins and 2 predicted proteins. The SDH1 protein was identified as both an intra and inter-modular hub.

**Table 5.**
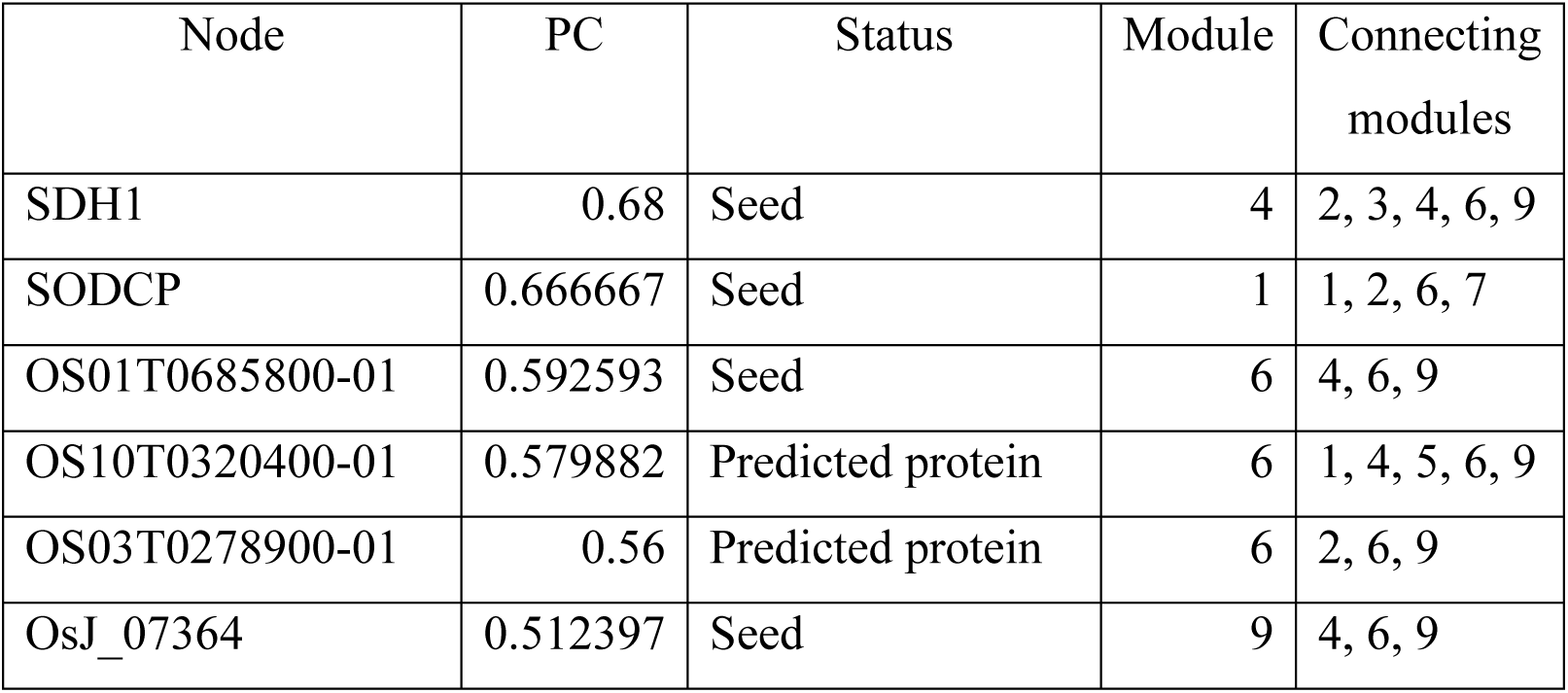
The detected inter-modular hub proteins with their associated PC scores, status as a seed protein or a prediction, the module to which they belong, and the connecting modules in the seed development sub-network.

## 4 Discussion

### 4.1 Analysis of detected seed development sub-modules and intra-modular hubs

The Louvain algorithm uncovered 14 sub-modules associated with seed development, unraveling the PPI network landscape underlying rice seed growth. A comprehensive analysis of the 14 sub-modules and their intra-modular hubs can be found in Additional file 1: Table S5. Certain sub-modules, such as endosperm development (sub-module 6) and regulation of seed growth (sub-module 13), were directly associated with seed development, while others, such as transition metal ion homeostasis (sub-module 1), were indirectly influencing seed development. Of the directly associated ones, sub-module 13 regulates rice seed growth, influencing grain size, weight, and quality. Crucial hub proteins of this sub-module include ILI5, instrumental in controlling grain length, and APG, the module hub responsible for regulating cell division and carbohydrate metabolism [60,61]. Additionally, predicted proteins GAI and OS03T0639300-01 exhibit significant interactions within the module, suggesting their potential roles in rice seed growth, warranting further investigation. The sub-module 5, which governs endosperm development, is the other sub-module directly associated with seed development. Endosperm development is crucial for rice grain development as it provides essential nutrients. This analysis revealed the presence of important proteins, such as OsFIE2 (OsJ_25971) and METB1 within the module, that are crucial for endosperm development in rice grains [62,63].

Of the indirect associations to seed development, sub-module 1 is annotated to essential transition metal ion homeostasis, crucial for enriching rice with iron (Fe) and zinc (Zn). Predicted proteins such as HMA2 within this module play pivotal roles in metal transport and contribute to grain weight [64]. Interestingly, predicted proteins HMA4 and Osj_26405 emerge as module hubs (Additional file 1: Table S4), with HMA4 notably linked to copper accumulation, suggesting potential roles in grain development [65]. Osj_26405, a member of the HMA domain-containing protein family, is essential for metal ion homeostasis and detoxification. While it shows differential expression during grain development, its specific role remains unclear due to limited literature evidence.

The sub-module 2 is annotated to the glycolytic process, which provides energy and carbon skeletons for cellular metabolism [66]. Phosphoglycerate kinase (PGK) serves as the central hub in sub-module 2, and according to literature, increased PGK expression in transgenic rice seeds raised the level of pyruvate, which elevated carotenoids levels in rice seeds [67]. Therefore, the expression of PGK may play a pivotal role in developing high-carotenoid rice seeds in combination with other necessary enzymes to address human dietary needs. Sub-module 3 of the seed development sub-network is associated with protein refolding, crucial during rice grain development, to ensure the correct folding and assembly of newly synthesized proteins in the endoplasmic reticulum, before being transported to their final destination [68]. OsJ_31804 (chaperonin 60) was identified as an intra-modular hub in sub-module 3. It is a heat shock protein that can stabilize and refold proteins during high temperatures, and protects rice grain storage components, including starch, proteins, and RNA, against detrimental effects [69].

The sub-module 4 of the rice seed development sub-network is associated with the leucine biosynthetic process. Rice grain’s essential amino acid levels, such as the leucine level, are important for its nutritional quality, but the mechanisms behind their accumulation are not yet fully understood [70]. OS03T0655700-01/OsIDHa, belonging to the Isocitrate dehydrogenase NAD-dependent protein family, was identified as an intra-modular hub. It participates in the tricarboxylic acid cycle, which is critical in cellular energy metabolism and is involved in the production of ATP [71]. However, its potential involvement in the leucine biosynthetic process, which is energy-dependent, is not yet studied. Hence, further studies may unravel the potential interplay between OsIDHa and the leucine biosynthetic pathway in rice grain development.

Hydrogen ion transmembrane transport (sub-module 6) is crucial for rice grain development as it maintains proper pH levels and ion homeostasis within cells. Various proton pumps and transporters are responsible for this process that may associated with seed development [72]. The protein OsJ_16709, which was revealed as a hub within the sub-module 6, belongs to the Inorganic Pyrophosphatase protein family. This protein plays a critical role in the metabolism of phosphate-containing compounds by hydrolyzing diphosphate in the presence of water. Although its involvement in H+ ion transmembrane transport is uncertain, the hydrolysis reaction of diphosphate releases H+ ions that can participate in the electrochemical gradient driving ion transport across biological membranes.

Sub-module 7 is associated with the methionine biosynthetic process, vital for plant growth. One of its module hubs, DHAR1, is known to increase rice grain yield and biomass when overexpressed. It also improves the Ascorbic Acid and redox homeostasis [73]. The sub-module 8 of the network is associated with the proteasomal ubiquitin-independent protein catabolic process. The regulation of cellular transitions is largely governed by protein turnover, with ubiquitin-independent mechanisms serving as effective means for identifying proteins for degradation [74]. Regulating the abundance of specific proteins and maintaining protein homeostasis is crucial in rice seed development, and protein turnover plays a vital role in this process. It is especially important during the transition between different growth stages. OsRPT2b, a member of the 26S proteasome regulatory subunit P45-like protein family, was revealed as an intra-modular hub in sub-module 8. This protein is crucial for the 26S proteasome regulation, responsible for the degradation of intracellular proteins in the ubiquitin-proteasome system, and exhibits variable expression levels during rice grain development [75]. However, further experimental evidence is needed to confirm its involvement in seed development.

The sub-module 9 of the network is associated with translation, which is crucial for rice grain protein synthesis. RPL3B, a predicted protein, was revealed as an intra-modular hub in this sub-module. It belongs to the ribosomal protein L3 family and its mutations impact ribosome biogenesis in rice, which results in abnormal architecture [76]. This predicted hub is an ideal candidate for further studies focusing on improving rice yield. The ethylene-activated signaling pathway is associated with sub-module 10 in this study. Ethylene is a gaseous hormone in plants that plays a pivotal role in multiple stages of rice grain development, such as germination, seedling growth, and grain maturation [77]. The protein MPK1 was detected as an intra-modular hub in this sub-module. It is a member of the mitogen-activated protein kinase (MAPK) family of proteins and plays a vital role in intracellular signaling pathways. This protein family responds to various stimuli and regulates essential cellular processes, including cell proliferation, differentiation, apoptosis, and stress response [78]. Through the analysis of CRISPR-edited mutants, a study has uncovered the significant role of MPK1 in rice development [79]. Although its involvement with ethylene-signaling pathways is poorly understood, it closely interacts with ethylene-related signaling proteins such as OsEIL2 and EIA1 [80], garnering further investigations into this module hub.

The cell differentiation is linked to sub-module 11, and it is essential for the growth and development of rice grains as it contributes to the architecture of the grain. The duration of cellularization in rice typically spans a period of 3 to 5 days [81]. The proper timing of cellularization in rice is dependent on the correct expression of various genes involved in the cell cycle, including CYCB2, which belongs to the cyclin protein family. In this study, CYCB2 was revealed as an intra-modular hub within the sub-module 11. Cyclins are well known for their involvement in regulating cell division and differentiation via cyclin-CDK complex control. CYCB2 has also been implicated in regulating endosperm development and seed size [82].

The sub-module 12 in this analysis is associated with the Abscisic acid-activated signaling pathway and ABI5 serves as an intra-modular hub which was predicted during the analysis. It plays a crucial role in multiple functions such as grain maturation, vigor, and dormancy, predominantly through regulating ABA-mediated transcription [83]. The sub-module 14 of the rice seed development sub-network is annotated to the brassinosteroid (BR)-mediated signaling pathway. Several studies have elaborated on the mediation of BRs in rice grain development. One study particularly emphasized that enhancing BR biosynthesis can boost crop productivity [84]. Additionally, it was observed that rice plants deficient in or insensitive to BRs produced smaller, shorter seeds [85]. Sterol delta-7-reductase (OS02T0465400-01) was identified as an intra-modular hub of this sub-module. This enzyme is involved in the biosynthesis of brassinosteroids [86]. However, its direct involvement with seed development is yet to be understood and serves as a candidate for future studies.

### 4.2 Analysis of inter-modular hubs

Six inter-modular hubs were discovered during the analysis and they are described in Additional file 1: Table S6. Among the discovered, OS01T0685800-01 and OS10T0320400-01, both predicted proteins for rice seed development, belong to the ATP synthase family, hinting at their involvement in ATP biogenesis. OS10T0320400-01 acts as a connector hub, linking translation, endosperm development, hydrogen ion transmembrane transport, and transitional ion homeostasis, while OS01T0685800-01 connects pathways such as hydrogen ion transmembrane transport, leucine biosynthesis, and translation. Despite the energy-intensive nature of these pathways, no studies have yet confirmed their mediation through these inter-modular hubs. Therefore, these are ideal candidates for future investigations.

Furthermore, SDH1 was revealed as both an inter-modular and an intra-modular hub during the analysis. It was found in the sub-module 4, which is annotated to leucine biosynthesis and it interconnects various sub-modules, including hydrogen ion transmembrane transport, protein refolding, glycolytic process, and translation. SDH1 is known to be associated with the tricarboxylic acid (TCA) cycle. Hydrogen ion (H+) transmembrane transport in rice plays a crucial role in regulating enzymes involved in the TCA cycle in the mitochondria [87]. The regulation of enzymes in the TCA cycle is complex, involving multiple levels of control, including transcriptional and translation regulation [71]. Protein biosynthesis and refolding in cells require ATP-dependent mechanisms, which can be energy-intensive [88]. Additionally, the redox status of cells can impact protein folding [73]. These findings emphasize the complex interplay between TCA cycle intermediates, protein translation, and H+ ion transport in developing rice grain cells. Importantly, SDH1 appears to be a crucial protein regulating the crosstalk between those sub-modules and an ideal genetic engineering target in future experiments.

### 4.3 Using an ensemble method and selection of the prediction thresholds

Ensemble methods combine the strengths of multiple models to improve the accuracy of predictions [42]. The rule of sum, a widely used technique to produce ensemble models, involves adding the predicted probabilities from each model to obtain a combined confidence score [24]. This study used the rule of sum to integrate four widely used network-based candidate protein prediction algorithms to predict novel seed development proteins. Using this ensemble model reduces the error and bias caused by individual algorithms and ensures the most accurate predictions. To our knowledge, this is the first instance where an ensemble of network-based prediction algorithms was used to predict novel plant proteins.

The selection of the prediction score cutoff for novel candidate protein predictions is a crucial factor in PPI analysis [89]. It impacts the sensitivity and specificity of predicted interactions. Raising the cutoff enhances specificity but decreases sensitivity while lowering the cutoff enhances sensitivity but decreases specificity. Hence, the choice of cutoff must strike a balance between these two parameters. This study determined the optimal cutoffs for the most reliable predictions using a modified precision top-N curve. Unlike, the conventional approach of simply calculating the precision at the N number of top-ranked predictions [90], the modified curve considers the percentage of overlap between the DEPs and the top-ranked N predictions. Here, the DEPs are differentially expressed proteins associated with seed development collected from literature and act as an independent source of validation for the predicted proteins. Using the traditional top-N curve for detecting the cut-off may lead to a substantial inclusion of false positives in the results, which is considerably reduced when using the modified curve.

In PPI analysis, intra-modular hubs are important for identifying protein complexes as they often represent the core proteins within a module. The specific cutoff for picking hubs can have a significant impact on the results [91]. The methods used to identify intra-modular hubs, i.e., hubs that have a high number of interactions within a module, typically involve calculating the degree of each protein either within the whole network or within the module and selecting the proteins with the highest degree, often the top 10% of proteins [92]. However, this technique can introduce a bias towards detecting a majority of hubs from more highly connected modules. Hence the within-module degree z-score (Z), which gauges the extent to which a node’s degree centrality differs from the anticipated degree of nodes with the same module membership, was used for this study [53]. Previous works have used intra-modular z-score cutoffs of 2.5 [93] and 1.5 [54] based on the properties of the network and research query. This study used 1.5 as the cutoff as 2.5 was unable to identify the protein with the highest within-module degree in the network.

For the prediction of inter-modular hubs, which connect different modules, the PC values were calculated. The choice of a PC cutoff for their detection in a PPI network may depend on the specific research question and the characteristics of the network being analyzed[94]. Inter-modular hubs were selected using a PC cutoff of 0.5, which has been employed in previous studies [95]. All of the identified hubs were connected to at least three sub-modules.

## 5 Conclusions

In this study, an ensemble of network-based algorithms was employed to predict new candidate proteins associated with rice seed development and study their systems biology landscape. The use of the ensemble approach was novel in predicting candidate proteins and the relevance of the predicted candidates was demonstrated by the enrichment of crucial GO terms such as “biosynthesis of organic hydroxy compounds” and “organic hydroxy compound metabolic process.” Furthermore, the sub-module analysis revealed specific pathways linked to seed growth regulation, endosperm development, and general processes such as translation, protein refolding, cell differentiation, and plant hormone signaling.

To identify promising targets for developing high-yielding rice varieties, the study highlighted key proteins crucial for seed development, known as hubs. Among the 17 intra-modular hubs, DHAR1, OsJ_16709, CYCB2-2, HMA4, and OsJ_26405 were the top five according to their Z-scores. Additionally, six inter-modular hubs that regulate cross-talk between different sub-modules were identified, including two predicted proteins and four seed proteins. Of these, the SDH1 protein achieved the highest score and was also revealed to have a dual role as an intra-modular hub, underscoring its potential as a target for genetic improvements. Newly predicted proteins such as RPL3B, OS02T0465400-01, OS10T0320400-01, HMA4, and ABI5 were also identified as important hub proteins potentially crucial for the seed development process.

Collectively, these novel protein candidates offer promising avenues for gaining deeper insights into the regulatory mechanisms governing rice grain development. This work provides the first comprehensive view of the protein interaction landscape associated with rice seed development. It is crucial to experimentally validate the identified hub proteins and predicted proteins in future studies as they could be important targets for genetic research aiming to improve rice grain quality and yield in rice. This study demonstrates how systems biology analysis techniques can be used for studying crucial developmental biology processes and serves as a blueprint for future studies.

## 6 Supplementary information

**Additional file 1:** The supplementary tables provided in this file contain information on seed proteins, results of enrichment analysis, and findings from the pathway, inter-modular hub, and intra-modular hub analyses referenced throughout the manuscript.

## 7 Declarations

### Ethics approval and consent to participate

Not Applicable.

### Consent for publication

All authors have read and approved the final manuscript for publication.

### Data and materials availability

All data generated or analyzed in support of the findings of this study are included in the manuscript and its supplementary information file. The Python codes used for this study are available at https://github.com/rashpr88/RiceSeedDevelopment.

### Author contribution

All authors were involved in planning and designing the experiments. PCF conceived the study and MRP wrote the Python scripts for the analysis, except for the RWR script, which was written by JW. MRP performed the experiments under the supervision of PCF. PCF and JW contributed to the preparation of the final manuscript. All authors analyzed the results and read and approved the final version of the manuscript.

## References

1. Huang R, Jiang L, Zheng J, Wang T, Wang H, Huang Y, et al. Genetic bases of rice grain shape: So many genes, so little known. Trends Plant Sci. 2013. p. 218–26.

2. Waterworth WM, Bray CM, West CE. Seeds and the art of genome maintenance. Front Plant Sci. Frontiers Media S.A.; 2019.

3. King T, Cole M, Farber JM, Eisenbrand G, Zabaras D, Fox EM, et al. Food safety for food security: Relationship between global megatrends and developments in food safety. Trends Food Sci Technol. 2017;68:160–75.

4. Sadigov R. Rapid Growth of the World Population and Its Socioeconomic Results. Scientific World Journal. 2022;2022.

5. Deng ZY, Gong CY, Wang T. Use of proteomics to understand seed development in rice. Proteomics. 2013. p. 1784–800.

6. An L, Tao Y, Chen H, He M, Xiao F, Li G, et al. Embryo-Endosperm Interaction and Its Agronomic Relevance to Rice Quality. Front Plant Sci. 2020.

7. Kozaki A, Aoyanagi T. Molecular Aspects of Seed Development Controlled by Gibberellins and Abscisic Acids. Int J Mol Sci. 2022.

8. Tappiban P, Ying Y, Xu F, Bao J. Proteomics and post-translational modifications of starch biosynthesis-related proteins in developing seeds of rice. Int J Mol Sci. 2021;22.

9. Wimalagunasekara S, Tirimanne S, Fernando PC. Protein-protein interaction (PPI) network analysis reveals important hub proteins and sub-network modules for root development in rice (Oryza sativa).

10. Vella D, Marini S, Vitali F, Di Silvestre D, Mauri G, Bellazzi R. MTGO: PPI Network Analysis Via Topological and Functional Module Identification. Sci Rep. 2018;8.

11. Fernando PC, Mabee PM, Zeng E. Integration of anatomy ontology data with protein-protein interaction networks improves the candidate gene prediction accuracy for anatomical entities. BMC Bioinformatics. 2020;21.

12. Sharan R, Ulitsky I, Shamir R. Network-based prediction of protein function. Mol Syst Biol. Nature Publishing Group; 2007. p. 1–13.

13. Fernando PC, Mabee PM, Zeng E. Protein–protein interaction network module changes associated with the vertebrate fin-to-limb transition. Sci Rep. 2023;13.

14. Roy A, Kucukural A, Zhang Y. I-TASSER: A unified platform for automated protein structure and function prediction. Nat Protoc. 2010;5.

15. Fortunato S. Community detection in graphs. 2009; Available from: http://arxiv.org/abs/0906.0612

16. Blondel VD, Guillaume J-L, Lambiotte R, Lefebvre E. Fast unfolding of communities in large networks. 2008; Available from: http://arxiv.org/abs/0803.0476

17. Kiran M, Nagarajaram HA. Interaction and localization diversities of global and local hubs in human protein-protein interaction networks. Mol Biosyst. 2016;12:2875–82.

18. He X, Zhang J. Why do hubs tend to be essential in protein networks? PLoS Genet. 2006;2:0826–34.

19. Hasan MI, Rahman MH, Islam MB, Islam MZ, Hossain MA, Moni MA. Systems Biology and Bioinformatics approach to Identify blood based signatures molecules and drug targets of patient with COVID-19. Inform Med Unlocked. 2022;28.

20. Wu B, Xi S. Bioinformatics analysis of differentially expressed genes and pathways in the development of cervical cancer. BMC Cancer. 2021;21.

21. Zhuang DY, Jiang LI, He QQ, Zhou P, Yue T. Identification of hub subnetwork based on topological features of genes in breast cancer. Int J Mol Med. 2015;35.

22. Wimalagunasekara SS, Weeraman JWJK, Tirimanne S, Fernando PC. Protein-protein interaction (PPI) network analysis reveals important hub proteins and sub-network modules for root development in rice (Oryza sativa). Journal of Genetic Engineering and Biotechnology. 2023;21.

23. Backiyarani S, Sasikala R, Sharmiladevi S, Uma S. Decoding the molecular mechanism of parthenocarpy in Musa spp. through protein–protein interaction network. Sci Rep. 2021;11.

24. Dua S, Chowriappa P. Data Mining for Bioinformatics.

25. STRING: functional protein association networks [Internet]. [cited 2024 Aug 19]. Available from: https://string-db.org/

26. Deng JL, Xu YH, Wang G. Identification of potential crucial genes and key pathways in breast cancer using bioinformatic analysis. Front Genet. 2019;10.

27. Chen L, Gao W, Chen S, Wang L, Zou J, Liu Y, et al. High-resolution QTL mapping for grain appearance traits and co-localization of chalkiness-associated differentially expressed candidate genes in rice. Rice. 2016;9.

28. Li C, Liu CQ, Zhang HS, Chen CP, Yang XR, Chen LF, et al. Lps1, encoding iron-sulfur subunit sdh2-1 of succinate dehydrogenase, affects leaf senescence and grain yield in rice. Int J Mol Sci. 2021;22:1–20.

29. Mahto A, Mathew IE, Agarwal P. Decoding the Transcriptome of Rice Seed During Development. Advances in Seed Biology. InTech; 2017.

30. Kim YJ, Kim S-I, Kesavan M, Kwak JS, Song JT, Seo HS. Ascorbate Peroxidase OsAPx1 is Involved in Seed Development in Rice. Plant Breed Biotechnol. 2015;3:11–20.

31. Yang Y, Dai L, Xia H, Zhu K, Liu H, Chen K. Protein profile of rice (Oryza sativa) seeds [Internet]. 2013. Available from: www.sbg.org.br

32. You C, Chen L, He H, Wu L, Wang S, Ding Y, et al. iTRAQ-based proteome profile analysis of superior and inferior Spikelets at early grain filling stage in japonica Rice. BMC Plant Biol. 2017;17.

33. QuickGO::Annotation List [Internet]. [cited 2024 Aug 19]. Available from: https://www.ebi.ac.uk/QuickGO/annotations

34. Xue LJ, Zhang JJ, Xue HW. Genome-wide analysis of the complex transcriptional networks of rice developing seeds. PLoS One. 2012;7.

35. Lee J, Koh HJ. A label-free quantitative shotgun proteomics analysis of rice grain development. Proteome Sci. 2011;9.

36. Kotu V, Deshpande B. Predictive analytics and data mining: concepts and practice with RapidMiner.

37. Cao M, Pietras CM, Feng X, Doroschak KJ, Schaffner T, Park J, et al. New directions for diffusion-based network prediction of protein function: Incorporating pathways with confidence. Bioinformatics. 2014;30.

38. Hishigaki H, Nakai K, Ono T, Tanigami A, Takagi T. Assessment of prediction accuracy of protein function from protein-protein interaction data. Yeast. 2001;18.

39. Nabieva E, Jim K, Agarwal A, Chazelle B, Singh M. Whole-proteome prediction of protein function via graph-theoretic analysis of interaction maps. Bioinformatics. 2005;21.

40. Cowen L, Ideker T, Raphael BJ, Sharan R. Network propagation: A universal amplifier of genetic associations. Nat Rev Genet. Nature Publishing Group; 2017. p. 551–62.

41. Fu J, Gao J, Liang Z, Yang D. PDI-regulated disulfide bond formation in protein folding and biomolecular assembly. Molecules. MDPI AG; 2021.

42. Xiao Y, Wu J, Lin Z, Zhao X. A deep learning-based multi-model ensemble method for cancer prediction. Comput Methods Programs Biomed. 2018;153:1–9.

43. Schwikowski B, Uetz P, Fields S. A network of protein-protein interactions in yeast. Nat Biotechnol. 2000;18.

44. Nabieva E, Jim K, Agarwal A, Chazelle B, Singh M. Whole-proteome prediction of protein function via graph-theoretic analysis of interaction maps. Bioinformatics. 2005;21.

45. Jiang Z, Liu H, Fu B, Wu Z, Zhang T. Recommendation in heterogeneous information networks based on generalized random walk model and Bayesian Personalized Ranking. WSDM 2018 - Proceedings of the 11th ACM International Conference on Web Search and Data Mining. 2018.

46. Zhang Z, Zhang J. A big world inside small-world networks. PLoS One. 2009;4.

47. Wu TY, Müller M, Gruissem W, Bhullar NK. Genome Wide Analysis of the Transcriptional Profiles in Different Regions of the Developing Rice Grains. Rice. 2020;13.

48. Li X, Lv J, Yi Z. Outlier Detection Using Structural Scores in a High-Dimensional Space. IEEE Trans Cybern. 2020;50.

49. DAVID Functional Annotation Bioinformatics Microarray Analysis [Internet]. [cited 2024 Aug 19]. Available from: https://david.ncifcrf.gov/

50. Fan G, Wei J. Identification of potential novel biomarkers and therapeutic targets involved in human atrial fibrillation based on bioinformatics analysis. Kardiol Pol. 2020;78:694–702.

51. Alcalá-Corona SA, Sandoval-Motta S, Espinal-Enríquez J, Hernández-Lemus E. Modularity in Biological Networks. Front Genet. 2021.

52. Tang D, Zhao X, Zhang L, Wang Z, Wang C. Identification of hub genes to regulate breast cancer metastasis to brain by bioinformatics analyses. J Cell Biochem. 2019;120.

53. McGarry K, Daniel U. Computational techniques for identifying networks of interrelated diseases. 2014 14th UK Workshop on Computational Intelligence, UKCI 2014 - Proceedings. Institute of Electrical and Electronics Engineers Inc.; 2014.

54. Zilidou VI, Frantzidis CA, Romanopoulou ED, Paraskevopoulos E, Douka S, Bamidis PD. Functional Re-organization of Cortical Networks of Senior Citizens After a 24-Week Traditional Dance Program. Front Aging Neurosci. 2018;10.

55. Vértes PE, Rittman T, Whitaker KJ, Romero-Garcia R, Váša F, Kitzbichler MG, et al. Gene transcription profiles associated with inter-modular hubs and connection distance in human functional magnetic resonance imaging networks. Philosophical Transactions of the Royal Society B: Biological Sciences. 2016;371.

56. Liu Y, Hong X, Bengson JJ, Kelley TA, Ding M, Mangun GR. Deciding Where to Attend: Large-scale Network Mechanisms Underlying Attention and Intention Revealed by Graph-theoretic Analysis. 2017.

57. Varadi M, Anyango S, Deshpande M, Nair S, Natassia C, Yordanova G, et al. AlphaFold Protein Structure Database: Massively expanding the structural coverage of protein-sequence space with high-accuracy models. Nucleic Acids Res. 2022;50.

58. Jumper J, Evans R, Pritzel A, Green T, Figurnov M, Ronneberger O, et al. Highly accurate protein structure prediction with AlphaFold. Nature. 2021;596.

59. Zhou Y-F, Qing T, Shu X-L, Liu J-X. Unfolded protein response and storage product accumulation in rice grains. Seed Biology. 2022;1.

60. Usman B, Nawaz G, Zhao N, Liu Y, Li R. Generation of high yielding and fragrant rice (Oryza sativa l.) lines by CRISPR/Cas9 targeted mutagenesis of three homoeologs of cytochrome p450 gene family and osbadh2 and transcriptome and proteome profiling of revealed changes triggered by mutations. Plants. 2020;9:1–28.

61. Heang D, Sassa H. An atypical bHLH protein encoded by POSITIVE REGULATOR OF GRAIN LENGTH 2 is involved in controlling grain length and weight of rice through interaction with a typical bHLH protein APG. Breed Sci. 2012;62:133–41.

62. Nallamilli BRR, Zhang J, Mujahid H, Malone BM, Bridges SM, Peng Z. Polycomb Group Gene OsFIE2 Regulates Rice (Oryza sativa) Seed Development and Grain Filling via a Mechanism Distinct from Arabidopsis. PLoS Genet. 2013;9.

63. Wang L, Yuan J, Ma Y, Jiao W, Ye W, Yang DL, et al. Rice Interploidy Crosses Disrupt Epigenetic Regulation, Gene Expression, and Seed Development. Mol Plant. 2018;11:300– 14.

64. Yamaji N, Xia J, Mitani-Ueno N, Yokosho K, Ma JF. Preferential delivery of zinc to developing tissues in rice is mediated by P-type heavy metal ATPase OsHMA2. Plant Physiol. 2013;162.

65. Huang XY, Deng F, Yamaji N, Pinson SRM, Fujii-Kashino M, Danku J, et al. A heavy metal P-type ATPase OsHMA4 prevents copper accumulation in rice grain. Nat Commun. 2016;7.

66. Zhang L, Ren Y, Lu B, Yang C, Feng Z, Liu Z, et al. FLOURY ENDOSPERM7 encodes a regulator of starch synthesis and amyloplast development essential for peripheral endosperm development in rice. J Exp Bot. 2016;67:633–47.

67. Gayen D, Ghosh S, Paul S, Sarkar SN, Datta SK, Datta K. Metabolic regulation of carotenoid-enriched golden rice line. Front Plant Sci. 2016;7.

68. He W, Wang L, Lin Q, Yu F. Rice seed storage proteins: Biosynthetic pathways and the effects of environmental factors. J Integr Plant Biol. 2021.

69. Timabud T, Yin X, Pongdontri P, Komatsu S. Gel-free/label-free proteomic analysis of developing rice grains under heat stress. J Proteomics. 2016;133:1–19.

70. Shi Y, Zhang Y, Sun Y, Xie Z, Luo Y, Long Q, et al. Natural variations of OsAUX5, a target gene of OsWRKY78, control the neutral essential amino acid content in rice grains. Mol Plant. 2023;16:322–36.

71. Zhang Y, Fernie AR. On the role of the tricarboxylic acid cycle in plant productivity. J Integr Plant Biol. Blackwell Publishing Ltd; 2018. p. 1199–216.

72. Li Y, Fan C, Xing Y, Yun P, Luo L, Yan B, et al. Chalk5 encodes a vacuolar H + - translocating pyrophosphatase influencing grain chalkiness in rice. Nat Genet. 2014;46:398– 404.

73. Kim YS, Kim IS, Bae MJ, Choe YH, Kim YH, Park HM, et al. Homologous expression of cytosolic dehydroascorbate reductase increases grain yield and biomass under paddy field conditions in transgenic rice (Oryza sativa L. japonica). Planta. 2013;237:1613–25.

74. Erales J, Coffino P. Ubiquitin-independent proteasomal degradation. Biochim Biophys Acta Mol Cell Res. 2014. p. 216–21.

75. Timabud T, Yin X, Pongdontri P, Komatsu S. Gel-free/label-free proteomic analysis of developing rice grains under heat stress. J Proteomics. 2016;133:1–19.

76. Zheng M, Wang Y, Liu X, Sun J, Wang Y, Xu Y, et al. The RICE MINUTE-LIKE1 (RML1) gene, encoding a ribosomal large subunit protein L3B, regulates leaf morphology and plant architecture in rice. J Exp Bot. 2016;67:3457–69.

77. Yin CC, Zhao H, Ma B, Chen SY, Zhang JS. Diverse roles of ethylene in regulating agronomic traits in rice. Front Plant Sci. 2017;8.

78. Kyosseva S V. MITOGEN-ACTIVATED PROTEIN KINASE SIGNALING. 2004.

79. Minkenberg B, Xie K, Yang Y. Discovery of rice essential genes by characterizing a CRISPR-edited mutation of closely related rice MAP kinase genes. Plant Journal. 2017;89:636–48.

80. Zhao H, Yin CC, Ma B, Chen SY, Zhang JS. Ethylene signaling in rice and Arabidopsis: New regulators and mechanisms. J Integr Plant Biol. Blackwell Publishing Ltd; 2021. p. 102– 25.

81. Wu X, Liu J, Li D, Liu CM. Rice caryopsis development II: Dynamic changes in the endosperm. J Integr Plant Biol. 2016;58:786–98.

82. Yang BJ, Wendrich JR, De Rybel B, Weijers D, Xue HW. Rice microtubule-associated protein IQ67-DOMAIN14 regulates grain shape by modulating microtubule cytoskeleton dynamics. Plant Biotechnol J. 2020;18:1141–52.

83. Ali F, Qanmber G, Li F, Wang Z. Updated role of ABA in seed maturation, dormancy, and germination. J Adv Res. Elsevier B.V.; 2022. p. 199–214.

84. Divi UK, Krishna P. Brassinosteroid: a biotechnological target for enhancing crop yield and stress tolerance. N Biotechnol. Elsevier; 2009. p. 131–6.

85. Hong Z, Ueguchi-Tanaka M, Fujioka S, Takatsuto S, Yoshida S, Hasegawa Y, et al. The rice brassinosteroid-deficient dwarf2 mutant, defective in the rice homolog of arabidopsis DIMINUTO/DWARF1, is rescued by the endogenously accumulated alternative bioactive brassinosteroid, dolichosterone. Plant Cell. 2005;17:2243–54.

86. Ito Y, Thirumurugan T, Serizawa A, Hiratsu K, Ohme-Takagi M, Kurata N. Aberrant vegetative and reproductive development by overexpression and lethality by silencing of OsHAP3E in rice. Plant Science. 2011;181:105–10.

87. Liao JL, Zhou HW, Peng Q, Zhong PA, Zhang HY, He C, et al. Transcriptome changes in rice (Oryza sativa L.) in response to high night temperature stress at the early milky stage. BMC Genomics. 2015;16.

88. Cao H, Duncan O, Millar AH. The molecular basis of cereal grain proteostasis. Essays Biochem. 2022.

89. Gorji-bahri G, Moghimi HR, Hashemi A. RAB5A is associated with genes involved in exosome secretion: Integration of bioinformatics analysis and experimental validation. J Cell Biochem. 2021;122.

90. Liong VE, Lu J, Wang G, Moulin P, Zhou J. Deep Hashing for Compact Binary Codes Learning.

91. Xiong Y, You W, Wang R, Peng L, Fu Z. Prediction and validation of hub genes associated with colorectal cancer by integrating PPI network and gene expression data. Biomed Res Int. 2017;2017.

92. Wang W, Shen J, Qi C, Pu J, Chen H, Zuo Z. The key candidate genes in tubulointerstitial injury of chronic kidney diseases patients as determined by bioinformatic analysis. Cell Biochem Funct. 2020;38:761–72.

93. Guimerà R, Mossa S, Turtschi A, Amaral LAN, Wachter KW. The worldwide air transportation network: Anomalous centrality, community structure, and cities’ global roles [Internet]. 2005. Available from: https://www.pnas.org

94. Joyce KE, Laurienti PJ, Burdette JH, Hayasaka S. A new measure of centrality for brain networks. PLoS One. 2010;5.

95. Liu Y, Hong X, Bengson JJ, Kelley TA, Ding M, Mangun GR. Deciding Where to Attend: Large-scale Network Mechanisms Underlying Attention and Intention Revealed by Graph-theoretic Analysis. 2017.

