## Additional file 1 for "Prediction of new candidate proteins and analysis of sub-modules and protein hubs associated with seed development in rice (*Oryza sativa*) using an ensemble network-based systems biology approach"

**Table S1** List of Seed Proteins and Corresponding Biological Processes. The retention status of each seed protein in the 0.7 combined score filtered network is also indicated

| Seed Protein | Retained after 0.7 combined score cutoff | Biological Processes |
|---|---|---|
| OLE18 | ✓ | Lipid storage, response to freezing, seed oil body biogenesis |
| OS03T0803700-01 | × | Embryo development ending in seed dormancy, leaf vascular tissue pattern formation, root development |
| OS05T0459700-01 | × | Abscisic acid-activated signaling pathway, mucilage biosynthetic process involved in seed coat development, multidimensional cell growth |
| GIF1 | ✓ | Cell population proliferation, leaf development, positive regulation of DNA-templated transcription, positive regulation of transcription by RNA polymerase II, seed development |
| PDIL1-1 | ✓ | Endosperm development, protein folding, protein folding in endoplasmic reticulum, response to endoplasmic reticulum stress |
| SS1 | ✓ | Amylopectin biosynthetic process, endosperm development, starch biosynthetic process |

| | | |
|---|---|---|
| OsJ_25971 | ✓ | Endosperm development, heterochromatin assembly, histone H3-K27 methylation, maintenance of seed dormancy, negative regulation of transcription by RNA polymerase II, post-embryonic plant organ development |
| OsJ_25973 | ✓ | Embryo development ending in seed dormancy, endosperm development, flower development, heterochromatin assembly, histone H3-K27 methylation, negative regulation of transcription by RNA polymerase II, seed development |
| SERK2 | ✓ | Brassinosteroid mediated signaling pathway, positive regulation of innate immune response, regulation of defense response to fungus, regulation of growth, somatic embryogenesis |
| MET1B | ✓ | DNA methylation on cytosine within a CG sequence, embryo development ending in seed dormancy, maintenance of DNA methylation, regulation of gene expression by genomic imprinting |
| SSIIIA | ✓ | Amylopectin biosynthetic process, endosperm development, starch biosynthetic process |
| OS06T0473800-00 | × | Lipid storage, response to freezing, seed oil body biogenesis |
| OS01T0605400-00 | ✓ | Embryo development ending in seed dormancy, pollen tube guidance |
| YSL15 | ✓ | Response to iron ion, seed development |

| | | |
|---|---|---|
| OS05T0510200-01 | ✓ | Chloroplast organization, embryo development ending in seed dormancy, protein insertion into mitochondrial outer membrane |
| OS01T0841100-01 | ✓ | Abscisic acid-activated signaling pathway, mucilage biosynthetic process involved in seed coat development, multidimensional cell growth |
| DN1 | ✓ | Floral organ structural organization, G protein-coupled receptor signaling pathway, regulation of seed development |
| AP2-2 | ✓ | Regulation of DNA-templated transcription, regulation of flower development, regulation of seed development, vegetative to reproductive phase transition of meristem |
| G1L6 | ✓ | Mrna transcription, negative regulation of DNA-templated transcription, post-embryonic plant morphogenesis, regulation of seed development, response to light stimulus |
| ILI6 | ✓ | Regulation of seed growth |
| FCA | ✓ | Positive regulation of mrna splicing, via spliceosome, regulation of cell size, regulation of flower development, regulation of seed growth, regulation of timing of transition from vegetative to reproductive phase |
| APG | ✓ | Regulation of seed growth |

| | | |
|---|---|---|
| BG1 | ✓ | Auxin transport, gravitropism, positive regulation of auxin mediated signaling pathway, regulation of seed growth, response to auxin |
| FLO6 | ✓ | Regulation of endosperm development, starch biosynthetic process |
| OsJ_06933 | ✓ | Translational elongation |
| SDH1 | ✓ | Mitochondrial electron transport, succinate to ubiquinone |
| OS03T0266200-01 | ✓ | Glycolytic process |
| OS01T0118000-01 | ✓ | Fructose 1,6-bisphosphate metabolic process, glycolytic process |
| OS08T0478800-01 | ✓ | Gluconeogenesis, glucose 6-phosphate metabolic process, glycolytic process |
| OS06T0114000-02 | ✓ | Protein folding |
| UGD1 | ✓ | Glycosaminoglycan biosynthetic process |
| OS02T0794700-01 | ✓ | Proteolysis |
| RINO1 | ✓ | Inositol biosynthetic process |
| AGPL2 | ✓ | Starch biosynthetic process |
| G6PGH2 | ✓ | D-gluconate catabolic process, pentose-phosphate shunt, oxidative branch, response to abscisic acid, response to cold, response to salt stress, response to water deprivation |
| OS11T0303050-00 | ✓ | Arginine biosynthetic process, argininosuccinate metabolic process, urea cycle |

| | | |
|---|---|---|
| OsJ_05367 | ✓ | Cytoplasmic translational initiation |
| OsIDHa | ✓ | Isocitrate metabolic process, tricarboxylic acid cycle |
| OS02T0158900-01 | ✓ | Cellular response to glucose starvation, protein phosphorylation, regulation of catalytic activity |
| OsJ_07364 | ✓ | Mitochondrial translational elongation, translational elongation |
| ILL1 | × | Auxin metabolic process |
| CYL2 | × | Tryptophan catabolic process to kynurenine |
| OS05T0230900-01 | ✓ | Methylglyoxal catabolic process to D-lactate via S-lactoyl-glutathione |
| GF14D | ✓ | Protein localization, signal transduction |
| GF14A | ✓ | Protein localization, signal transduction |
| OS01T0322300-00 | ✓ | Phosphate-containing compound metabolic process |
| Chib3H-h | ✓ | Defense response to fungus |
| DHAR1 | ✓ | Ascorbate glutathione cycle, cellular oxidant detoxification, protein glutathionylation |
| OsJ_29684 | ✓ | Glutathione metabolic process |
| OsJ_32231 | ✓ | Glutathione metabolic process |
| OsJ_25719 | ✓ | Polar nucleus fusion |

| | | |
|---|---|---|
| PAIR1 | ✓ | Embryo sac development, megasporogenesis, meiotic DNA double-strand break formation, microsporogenesis |
| PAIR2 | ✓ | Homologous chromosome pairing at meiosis,megasporogenesis,microsporogenesis,proteolysis involved in protein catabolic process |
| MEL1 | ✓ | Female meiotic nuclear division,histone H3-K9 demethylation, male meiotic nuclear division, megagametogenesis, microgametogenesis, negative regulation of histone H3-K9 acetylation, positive regulation of histone H3-K9 dimethylation, positive regulation of meiotic DNA double-strand break formation, post-transcriptional gene silencing by RNA |
| PAIR3 | ✓ | Homologous chromosome pairing at meiosis |
| MPK1 | ✓ | Intracellular signal transduction |
| CBP1 | ✓ | Secondary metabolic process |
| OsJ_09943 | ✓ | Microtubule cytoskeleton organization, mitotic cell cycle |
| OsJ_15699 | ✓ | Malate metabolic process, NADH metabolic process, oxaloacetate metabolic process, tricarboxylic acid cycle |
| SODCC1 | ✓ | Removal of superoxide radicals |
| SODCP | ✓ | Removal of superoxide radicals |

| | | |
|---|---|---|
| SODCC2 | ✓ | Removal of superoxide radicals |
| APX1 | ✓ | Cellular response to oxidative stress, hydrogen peroxide catabolic process, response to reactive oxygen species |
| RNB5 | ✓ | Protein complex oligomerization, protein folding, response to heat, response to hydrogen peroxide, response to salt stress |
| GW2 | ✓ | Protein ubiquitination, regulation of seed growth |
| OsJ_13316 | ✓ | Signal transduction |
| OS08T0537800-01 | ✓ | Protein K48-linked deubiquitination |
| OsJ_34870 | ✓ | Regulation of DNA-templated transcription |
| OsJ_15624 | ✓ | Photomorphogenesis, regulation of DNA-templated transcription |
| CYCB2-2 | ✓ | Mitotic cell cycle phase transition, regulation of cyclin-dependent protein serine/threonine kinase activity |
| MYB3R-2 | ✓ | Positive regulation of DNA-templated transcription, positive regulation of response to salt stress, positive regulation of response to water deprivation, regulation of DNA-templated transcription, response to cold |

| | | |
|---|---|---|
| OsJ_04024 | ✓ | Cellular response to unfolded protein, chaperone cofactor-dependent protein refolding, protein refolding |
| OS04T0486500-01 | ✓ | Mitotic spindle assembly checkpoint signaling |
| G6PGH1 | ✓ | D-gluconate catabolic process, pentose-phosphate shunt, oxidative branch, response to abscisic acid, response to cold, response to salt stress, response to water deprivation |
| V-ATPase B | ✓ | Vacuolar acidification |
| OsJ_07966 | ✓ | Aspartyl-trna aminoacylation |
| PDIL1-4 | ✓ | Protein folding, response to endoplasmic reticulum stress |
| OS01T0685800-01 | ✓ | Proton motive force-driven mitochondrial ATP synthesis |
| OsJ_22300 | ✓ | Gluconeogenesis,glycolytic process |
| GLYI-11 | ✓ | Methylglyoxal catabolic process to D-lactate via S-lactoyl-glutathione, response to osmotic stress, response to oxidative stress, response to salt stress |
| OsJ_12328 | ✓ | Carbohydrate metabolic process |
| OS12T0540900-01 | ✓ | Tryptophanyl-trna aminoacylation |
| OsJ_06100 | ✓ | Protein dephosphorylation |
| OsJ_35174 | ✓ | Leucine biosynthetic process |

| | | |
|---|---|---|
| OS03T0655700-01 | ✓ | Leucine biosynthetic process |
| OsJ_35666 | ✓ | Arginine biosynthetic process, argininosuccinate metabolic process, urea cycle |
| OS03T0363500-02 | ✓ | Transmembrane transport |
| OsJ_31804 | ✓ | Protein folding |
| OS07T0467900-01 | ✓ | Mitochondrial DNA replication, replication fork processing, response to UV, translesion synthesis |
| OsJ_07639 | ✓ | Proteasomal protein catabolic process |
| OsJ_02861 | ✓ | Isocitrate metabolic process, NADP metabolic process |
| OsJ_10329 | ✓ | Cellular response to unfolded protein, chaperone cofactor-dependent protein refolding, protein refolding |
| PBF1 | ✓ | Proteasomal protein catabolic process |
| OsJ_16709 | ✓ | Phosphate-containing compound metabolic process |
| OsJ_26737 | ✓ | Intracellular protein transport, SNARE complex disassembly |
| RPA1B | ✓ | DNA unwinding involved in DNA replication, double-strand break repair via homologous recombination, meiotic cell cycle, nucleotide-excision repair, telomere maintenance via telomerase |

| | | |
|---|---|---|
| B1045D11.6 | ✓ | Asparaginyl-trna aminoacylation |
| OS08T0308100-01 | ✓ | Cytoplasmic translational initiation |
| RACK1A | ✓ | Defense response to fungus, positive regulation of protein phosphorylation, positive regulation of respiratory burst, rescue of stalled ribosome |
| TPI | ✓ | Gluconeogenesis, glyceraldehyde-3-phosphate biosynthetic process, glycerol catabolic process, glycolytic process |
| UGP | ✓ | UTP: glucose-1-phosphate uridylyltransferase activity, glycogen metabolic process, UDP-glucose metabolic process |
| FBA3 | ✓ | Fructose 1,6-bisphosphate metabolic process, gluconeogenesis, glycolytic process |

**Table S2** Differentially Expressed Proteins (DEPs) associated with rice seed development retrieved from the literature

| Preferred Name | Annotation |
|---|---|
| OS10T0132300-01 | Jacalin-like lectin domain containing protein, expressed; Os10g0132300 protein; cDNA clone: 002-180-H05, full insert sequence |

| | |
|---|---|
| OsJ_23479 | Os07g0202450 protein; Os10g0486600 protein; Putative dim1p; Putative thioredoxin-like U5 small ribonucleoprotein particle protein; Thioredoxin-like protein 4A, putative, expressed; cDNA clone: 001-121-D12, full insert sequence; cDNA clone: J033122K22, full insert sequence |
| OsJ_13064 | Sex determination protein tassel seed 2, putative, expressed; Os03g0810800 protein; Putative alcohol dehydrogenase; Putative short chain alcohol dehydrogenase; Sex determination protein tassel seed 2, putative, expressed; cDNA clone: J023042C15, full insert sequence |
| OS03T0278900-01 | ATP synthase B chain, chloroplast, putative, expressed; H+-transporting ATP synthase chain 9-like protein; Os03g0278900 protein; cDNA clone: 001-205-A08, full insert sequence; cDNA clone: J013049M17, full insert sequence; Belongs to the ATPase B chain family |
| OsJ_13442 | Os03g0858400 protein; Putative WD repeat protein; WD domains, G-beta repeats containing protein, putative, expressed; cDNA clone: J033115J18, full insert sequence |
| OS03T0172000-01 | Nucleoporin family protein, putative, expressed; Os03g0172000 protein |
| OsJ_09866 | Methionyl-tRNA synthetase, putative, expressed; Os03g0209600 protein; cDNA clone: J013160O09, full insert sequence; Belongs to the class-I aminoacyl-tRNA synthetase family |
| OsJ_35872 | Histone H3, putative, expressed; Os12g0415800 protein |

| | |
|---|---|
| OS03T0810900-01 | cDNA clone: J023108N19, full insert sequence; Os03g0810900 protein; Patatin, putative, expressed; Putative patatin-like phospholipase; Putative phospholipase; cDNA clone: J023108N19, full insert sequence |
| Act | Actin; Actin-7, putative, expressed; Os11g0163100 protein; cDNA clone: 001-035-C05, full insert sequence; Belongs to the actin family |
| OS10T0506800-01 | cDNA clone: J013122F06, full insert sequence; Os10g0506800 protein; TPR Domain containing protein, expressed; Tetratricopeptide repeat, putative; cDNA clone: J013122F06, full insert sequence |
| OsJ_11750 | T-complex protein 1, beta subunit, putative, expressed; Molecular chaperone; assists the folding of proteins upon ATP hydrolysis |
| OsJ_13407 | Cleavage and polyadenylation specificity factor, 73 kDa subunit, putative, expressed; Os03g0852900 protein; Putative cleavage and polyadenylation specifity factor; Putative cleavage and polyadenylation specifity factor protein |
| OsJ_10521 | Os03g0300700 protein; Retrotransposon protein, putative, unclassified, expressed; cDNA clone: J023112P05, full insert sequence |
| OS12T0105700-01 | Os12g0105700 protein; Outer membrane protein, OMP85 family protein, expressed; cDNA clone: J013119K16, full insert sequence |
| OsRPT2b | 26S protease regulatory subunit 4, putative, expressed; 26S proteasome regulatory particle triple-A ATPase subunit2b; Os03g0298400 protein; cDNA clone: J023030D05, full insert sequence; Belongs to the AAA ATPase family |

| | |
|---|---|
| OS03T0741500-01 | cDNA clone: 002-172-E02, full insert sequence; Cytochrome P450 71D11, putative, expressed; Putative cytochrome P450 protein; cDNA clone: 002-172-E02, full insert sequence |
| OS03T0725000-01 | Mitochondrial 60S ribosomal protein L6, putative, expressed; Os03g0725000 protein; Putative ribosomal protein L6; cDNA clone: 001-025-E08, full insert sequence; cDNA clone: 002-157-C05, full insert sequence; Belongs to the universal ribosomal protein uL6 family |
| OS03T0722500-01 | Glucan endo-1,3-beta-glucosidase 7, putative, expressed; Glycosyl hydrolases family 17, putative; Os03g0722500 protein; cDNA clone: J023004N24, full insert sequence; Belongs to the glycosyl hydrolase 17 family |
| OsJ_23275 | 40S ribosomal protein S18, putative, expressed; Os03g0794700 protein; Os07g0173700 protein; Putative ribosomal protein S18; Putative ribosomal protein S18A; cDNA clone: 001-007-B06, full insert sequence; cDNA clone: 001-029-F08, full insert sequence; Belongs to the universal ribosomal protein uS13 family |
| OsJ_09272 | Glyceraldehyde-3-phosphate dehydrogenase; EST C74302(E30840) corresponds to a region of the predicted gene; Glyceraldehyde-3-phosphate dehydrogenase B, chloroplast, putative, expressed; Os03g0129300 protein; Putative glyceraldehyde-3-phosphate dehydrogenase |
| OsJ_09517 | Glutathione reductase, chloroplast, putative, expressed; Os03g0163300 protein; Putative glutathione reductase; Putative glutathione reductase (NADPH); cDNA clone: J033128J11, full insert sequence; Belongs to the class-I pyridine nucleotide-disulfide oxidoreductase family |

| | |
|---|---|
| OS03T0213100-01 | Os03g0213100 protein; Protein transport protein Sec61 alpha subunit isoform 2, putative, expressed; Belongs to the SecY/SEC61-alpha family |
| OsJ_12299 | Os03g0709100 protein; Plastocyanin-like domain containing protein, expressed; Plastocyanin-like domain, putative; cDNA clone: 001-024-C04, full insert sequence |
| OsJ_12021 | RanBP1 domain containing protein, expressed; cDNA, clone: J043022O19, full insert sequence |
| OsJ_10495 | 40S ribosomal protein S7, putative, expressed; Os03g0297100 protein; cDNA clone: 001-023-H09, full insert sequence; cDNA clone: J033136J20, full insert sequence; Belongs to the eukaryotic ribosomal protein eS7 family |
| OsJ_10726 | Eukaryotic translation initiation factor 2 beta subunit, putative, expressed; Os03g0333300 protein; cDNA clone: J033023N08, full insert sequence |
| OS10T0566700-00 | Chloroplast chaperonin 10, putative, expressed; Os10g0566700 protein; Putative chloroplast chaperonin; Belongs to the GroES chaperonin family |
| OsJ_11922 | Elongation factor Tu family protein, putative, expressed; Os03g0650700 protein; Putative Translation Elongation factor protein |
| OsJ_32962 | Os11g0147800 protein; Stem-specific protein; Stem-specific protein TSJT1, putative, expressed; cDNA clone: 001-039-G04, full insert sequence; cDNA clone: 006-211-B05, full insert sequence; cDNA clone: J023109H06, full insert sequence |

| | |
|---|---|
| OsJ_12657 | 26S proteasome non-ATPase regulatory subunit 10, putative, expressed; Putative ankyrin; cDNA clone: J013073E13, full insert sequence |
| OsJ_09152 | 70 kDa heat shock protein; Heat shock 70 kDa protein, mitochondrial, putative, expressed; Os03g0113700 protein; cDNA clone: J033148E17, full insert sequence |
| OsJ_36515 | Phosphoinositide phospholipase c; Phospholipase C, putative, expressed |
| OsJ_09121 | 40S ribosomal protein S17, putative, expressed; Os03g0109500 protein; Putative 40S ribosomal protein S17 |
| OsJ_33580 | Os11g0266000 protein; Transposon protein, putative, unclassified; Transposon protein, putative, unclassified, expressed; cDNA clone: J033028G10, full insert sequence |
| OS03T0126000-01 | Anthranilate phosphoribosyltransferase, chloroplast, putative, expressed; Os03g0126000 protein; cDNA clone: J033069P19, full insert sequence |
| OsJ_30578 | cDNA clone: J033109K13, full insert sequence; NHP2-like protein 1, putative, expressed; Os10g0124000 protein; Putative ribosomal protein L7Ae-like; cDNA clone: J033109K13, full insert sequence |
| OS10T0574800-01 | GTPase activating protein, putative, expressed; Os10g0574800 protein; Putative uncharacterized protein OSJNBa0003O19.4 |
| OS03T0840200-01 | Os03g0840200 protein; Putative F8K7.10 protein; WD40-like Beta Propeller Repeat family protein, expressed; cDNA clone: J013116D11, full insert sequence; cDNA clone: J023087F22, full insert sequence |

| | |
|---|---|
| OsJ_35976 | O-methyltransferase ZRP4, putative, expressed; Belongs to the class I-like SAM-binding methyltransferase superfamily. Cation-independent O- methyltransferase family |
| OS03T0720300-01 | Glutamate decarboxylase, putative, expressed; Os03g0720300 protein; Putative glutamate decarboxylase isozyme; Belongs to the group II decarboxylase family |
| OsJ_32460 | Hexose carrier protein HEX6, putative, expressed; Os10g0561300 protein; Putative hexose carrier protein; Putative monosaccharide transporter; cDNA clone: J013148F21, full insert sequence; Belongs to the major facilitator superfamily. Sugar transporter (TC 2.A.1.1) family |
| OS03T0418800-01 | Os03g0418800 protein; Nuclear transport factor 2 domain containing protein, expressed; Os03g0418800 protein; Putative GAP SH3 binding protein; cDNA clone: J033058N07, full insert sequence |
| bip108 | 60S ribosomal protein L22-2, putative, expressed; BRI1-KD interacting protein 108; Os03g0343500 protein; cDNA clone: 001-019-G04, full insert sequence; cDNA clone: 001-030-E04, full insert sequence; cDNA clone: J033022G11, full insert sequence |
| OS03T0835700-00 | Putative heat shock protein; Small heat shock protein, chloroplast, putative; Belongs to the small heat shock protein (HSP20) family |

| | |
|---|---|
| OsJ_12590 | 50S ribosomal protein L6, putative, expressed; Os03g0751400 protein; Putative chloroplast 50S ribosomal protein L6; Putative ribosomal protein L6; cDNA clone: J023004G24, full insert sequence; Belongs to the universal ribosomal protein uL6 family |
| OsJ_36723 | Os12g0594600 protein; Skp1 family, dimerisation domain containing protein, expressed; cDNA clone: J033031N08, full insert sequence |
| OsJ_10070 | Transmembrane 9 superfamily member; Endomembrane protein 70 containing protein, expressed; Os03g0237000 protein; Putative endosomal protein; cDNA clone: J013000C08, full insert sequence; Belongs to the nonaspanin (TM9SF) (TC 9.A.2) family |
| OS11T0153800-01 | 40S ribosomal protein S25, putative, expressed; Os11g0153800 protein; S25 ribosomal protein; cDNA clone: 001-018-C09, full insert sequence |
| OS03T0748300-01 | Aldose 1-epimerase family protein, expressed; Os03g0748300 protein; Putative antifreeze glycoprotein; cDNA clone: J013156O10, full insert sequence |
| OS10T0320400-01 | ATP synthase gamma chain, mitochondrial, putative, expressed; Os10g0320400 protein; cDNA clone: J023135E20, full insert sequence |
| OsJ_32235 | Glutathione S-transferase GSTU6, putative, expressed; Putative glutathione S-transferase; Putative glutathione S-transferase OsGSTU9 |

| | |
|---|---|
| OsJ_00798 | High-affinity nickel-transport family protein, putative, expressed; Os03g0156700 protein; Putative uncharacterized protein OSJNBa0011L14.10; Uncharacterized protein |
| OsJ_13099 | Os03g0815900 protein; Putative transcription elongation factor; Transcription elongation factor S-II family protein, expressed |
| RPL3B | 60S ribosomal protein L3, putative, expressed; Os11g0168200 protein; Ribosomal protein L3, putative; Ribosomal protein L3B; cDNA clone: J033040J04, full insert sequence; cDNA clone: J033041E13, full insert sequence; Belongs to the universal ribosomal protein uL3 family |
| OsJ_32925 | 2-isopropylmalate synthase B, putative, expressed |
| OsJ_36378 | Os12g0538900 protein; Vesicle tethering family protein, putative, expressed; cDNA clone: J013091G19, full insert sequence |
| OS04T0653600-01 | Os04g0653600 protein; cDNA clone: J033072A06, full insert sequence |
| OS11T0129700-01 | Os11g0129700 protein |
| OS03T0313100-01 | Os03g0313100 protein |
| OS01T0778100-01 | Uncharacterized protein; Os01g0778100 protein |
| OS01T0909500-02 | cDNA clone: J013116D08, full insert sequence |
| OS04T0690100-01 | Os04g0690100 protein |
| OsJ_19788 | Os05g0597100 protein; cDNA clone: J023142J09, full insert sequence |

| | |
|---|---|
| OS07T0101800-00 | C2H2-type zinc finger protein-like protein |
| ZFP39 | C2H2-type zinc finger transcription factor; Os07g0588700 protein; Putative zinc-finger protein 1 |
| OsJ_27725 | cDNA clone: J023004M17, full insert sequence; Os08g0485700 protein; Putative uncharacterized protein P0481F05.12; cDNA clone: J023004M17, full insert sequence |
| OS09T0449400-01 | annotation not available |
| Ehd2 | Cys2/His2-type zinc finger transcription factor; Early heading date 2 |
| OS12T0279100-01 | JmjC domain containing protein, expressed; Os12g0279100 protein |
| OS01T0277500-01 | Os01g0277500 protein; Ascorbate oxidase promoter-binding protein AOBP-like; Os01g0277500 protein |
| DOF2 | Putative uncharacterized protein; Transcription factor that may transactivate seed storage protein genes in developing seeds |
| DOF4 | Dof zinc finger protein 4; Transcription factor that may transactivate seed storage protein genes in developing seeds |
| OsJ_09572 | Dof domain, zinc finger family protein, expressed; Os03g0169600 protein; Putative H-protein promoter binding factor-2a; Putative zinc finger DNA-binding protein; cDNA clone: 001-035-D05, full insert sequence; cDNA clone: 006-303-F03, full insert sequence |
| OS03T0821200-00 | Os03g0821200 protein |
| OS05T0112200-01 | Os05g0112200 protein; Putative dof-type zinc finger domain-containing protein |
| OS07T0236700-01 | Os07g0236700 protein; Putative dof zinc finger protein; cDNA clone: J013155H18, full insert sequence |

| | |
|---|---|
| OsJ_25624 | AOBP-like protein; Os07g0685000 protein; Putative ascorbate oxidase promoter-binding protein AOBP; cDNA clone: 001-032-E07, full insert sequence |
| OS10T0406300-01 | Os10g0406300 protein |
| HOX16 | Homeobox-leucine zipper protein HOX16; Probable transcription factor |
| HOX4 | Homeobox-leucine zipper protein HOX4; Probable transcription activator that binds to the DNA sequence 5'-CAAT[AT]ATTG-3'. May be involved in the regulation of gibberellin signaling |
| HOX13 | Homeobox-leucine zipper protein HOX13; Probable transcription factor |
| HOX23 | Homeobox-leucine zipper protein HOX23; Probable transcription factor |
| HOX1 | Homeobox-leucine zipper protein HOX1; Probable transcription repressor involved leaf development. Binds to the DNA sequence 5'-CAAT[GC]ATTG-3'. May act as a regulatory switch to specify provascular cell fate |
| HOX15 | Homeobox-leucine zipper protein hox15; Probable transcription factor |
| HOX22 | Homeobox-leucine zipper protein HOX22; Probable transcription factor; Belongs to the HD-ZIP homeobox family. Class I subfamily |
| HOX2 | Homeobox-leucine zipper protein HOX2; Probable transcription factor that binds to the DNA sequence 5'-CAAT[GC]ATTG-3'; Belongs to the HD-ZIP homeobox family. Class II subfamily |

| | |
|---|---|
| HOX6 | Homeobox-leucine zipper protein HOX6; Probable transcription factor that binds to the DNA sequence 5'-CAAT[AT]ATTG-3' |
| HOX12 | Homeobox-leucine zipper protein HOX12; Probable transcription factor |
| ROC1 | Homeobox-leucine zipper protein roc1; Probable transcription factor that may be involved in protoderm differentiation and radial pattern formation during early embryogenesis |
| ROC2 | Homeobox-leucine zipper protein ROC2; Probable transcription factor |
| ROC7 | Homeobox-leucine zipper protein ROC7; Probable transcription factor |
| OS09T0526200-01 | annotation not available |
| ROC3 | Homeobox-leucine zipper protein ROC3; Probable transcription factor |
| TF1 | Homeobox-leucine zipper protein TF1; Probable transcription factor; Belongs to the HD-ZIP homeobox family. Class IV subfamily |
| TIP2 | Os01g0293100 protein; Putative bHLH transcription factor; Transcription factor that binds to the E-box-containing promoter regions of the transcription factors TDR and EAT1, activating their expression. May have a role in specifying the cell pattern of the inner anther walls and functioning in meiosis progression. Required for male reproduction. Acts downstream of UDT1 and GAMYB, but upstream of TDR1 and EAT1 in pollen development |
| OsJ_02312 | Basic helix-loop-helix (BHLH) family protein-like; Uncharacterized protein; cDNA clone: 002-101-F10, full insert sequence |

| | |
|---|---|
| OS01T0915600-01 | cDNA clone: J033022O19, full insert sequence; BHLH transcription factor; Os01g0915600 protein; Putative bHLH transcription factor; cDNA clone: J033022O19, full insert sequence |
| ILI5 | Transcription factor ili5; Atypical and probable non DNA-binding bHLH transcription factor that acts as a positive regulator of grain size. Binds the transcription repressor APG and forms a heterodimer of antagonistic basic helix-loop-helix transcription factors that regulates grain length and weight by controlling cell elongation in lemma and palea |
| OS02T0759000-01 | Os02g0759000 protein |
| OS02T0795800-00 | annotation not available |
| OS03T0231950-01 | Putative uncharacterized protein; Os03g0231950 protein |
| OS03T0279500-00 | annotation not available |
| OS03T0391700-01 | Expressed protein; Expressed protein (With alternative splicing); Os03g0391700 protein; cDNA clone: J013036B10, full insert sequence |
| OS03T0639300-01 | Expressed protein; Os03g0639300 protein |
| OS03T0725800-01 | Helix-loop-helix DNA-binding domain containing protein, expressed; Os03g0725800 protein; Putative symbiotic ammonium transport protein; Putative transcription factor; cDNA clone: J013129F07, full insert sequence |
| OS04T0557500-00 | annotation not available |

| | |
|---|---|
| EAT1 | OSJNBa0083N12.3 protein; Os04g0599300 protein; Transcription factor involved in the regulation of tapetum programmed cell death (PCD) and degradation during male reproductive development. Interacts with TDR and promote tapetal PCD by regulating the expression of RTS, and the two lipid-transfer proteins C4 and C6, which function in microspore development. Acts downstream from and interacts with TDR in the regulation of tapetal PCD. Regulates directly the aspartic protease AP25 and AP37 during tapetal PCD. May not target the cysteine protease CP1 |
| OS05T0455400-01 | Putative uncharacterized protein; Os05g0455400 protein |
| OS05T0501200-00 | annotation not available |
| OsJ_20430 | BHLH transcription factor PTF1; Os06g0193400 protein; cDNA clone: J013042L14, full insert sequence |
| OS06T0275600-01 | Os06g0275600 protein; Putative TA1 protein |
| OS07T0143200-00 | Transcription factor BHLH9-like protein |
| BHLH094 | Transcription factor bhlh094; Transcription factor that forms a ternary complex with RSS3 and TIFY11A/JAZ9 to negatively regulate jasmonate-responsive genes |
| UDT1 | Putative uncharacterized protein; Transcription factor that plays a crucial role in tapetum development. Required for male fertility and pollen differentiation within the developing anther. Plays a major role in maintaining tapetum development, starting in early meiosis. Required for pollen mother cell meiosis. May regulate the anther-specific cysteine protease CP1 |

| | |
|---|---|
| | and lipid-transfer proteins C4 and C6. Required for anther development. Functions in parallel with GAMYB to regulate early anther development. Functions upstream of the transcription factor TDR. |
| OS08T0490000-01 | cDNA clone: J023030J22, full insert sequence; BHLH protein family-like; BHLH transcription factor; Os08g0490000 protein; cDNA clone: J023030J22, full insert sequence |
| OsJ_27873 | Os08g0506700 protein; Putative transcription factor RAU1; cDNA clone: J023004M08, full insert sequence |
| OS09T0417400-00 | annotation not available |
| OS09T0474100-01 | cDNA clone: J023133H05, full insert sequence; BHLH-transcription factor; Os09g0474100 protein; TA1 protein-like; cDNA clone: J023133H05, full insert sequence |
| OS09T0487900-01 | Os09g0487900 protein |
| OsJ_29905 | Os09g0501600 protein; cDNA clone: 001-021-C10, full insert sequence |
| OsJ_32337 | Helix-loop-helix DNA-binding domain containing protein, expressed; Os10g0544200 protein; cDNA clone: 001-119-D04, full insert sequence |
| MYC2 | Os10g0575000 protein; Transcription factor MYC7E, putative, expressed; Transcriptional activator involved in jasmonate (JA) signaling pathway during spikelet development. Binds to the G2 region G-box (5'-CACGTG-3') of the MADS1 promoter and thus directly regulates the expression of MADS1. Its function in MADS1 activation is abolished by TIFY3/JAZ1 which directly target MYC2 during spikelet development |

| | |
|---|---|
| OS11T0158500-01 | Os11g0158500 protein |
| OS12T0632600-00 | Putative uncharacterized protein; Os12g0632600 protein |
| ARF1 | Auxin response factor 1; Auxin response factors (ARFs) are transcriptional factors that bind specifically to the DNA sequence 5'-TGTCTC-3' found in the auxin-responsive promoter elements (AuxREs) |
| ARF2 | Auxin response factor 2; Auxin response factors (ARFs) are transcriptional factors that bind specifically to the DNA sequence 5'-TGTCTC-3' found in the auxin-responsive promoter elements (AuxREs) |
| ARF3 | Auxin response factor 3; Auxin response factors (ARFs) are transcriptional factors that bind specifically to the DNA sequence 5'-TGTCTC-3' found in the auxin-responsive promoter elements (AuxREs) |
| ARF12 | Auxin response factor 12; Auxin response factors (ARFs) are transcriptional factors that bind specifically to the DNA sequence 5'-TGTCTC-3' found in the auxin-responsive promoter elements (AuxREs) |
| ARF15 | Auxin response factor 15; Auxin response factors (ARFs) are transcriptional factors that bind specifically to the DNA sequence 5'-TGTCTC-3' found in the auxin-responsive promoter elements (AuxREs) |
| ARF16 | Auxin response factor 16; Auxin response factors (ARFs) are transcriptional factors that bind specifically to the DNA sequence 5'-TGTCTC-3' found in the auxin-responsive promoter elements (AuxREs) |
| OS08T0520500-01 | annotation not available |

| | |
|---|---|
| ARF24 | Auxin response factor 24; Auxin response factors (ARFs) are transcriptional factors that bind specifically to the DNA sequence 5'-TGTCTC-3' found in the auxin-responsive promoter elements (AuxREs) |
| OS01T0174600-01 | Zinc finger CCCH domain-containing protein 1 |
| OsJ_002707 | Zinc finger CCCH domain-containing protein 9 |
| P0638D12.6 | Zinc finger CCCH domain-containing protein 10 |
| OS05T0128200-01 | Zinc finger CCCH domain-containing protein 33 |
| OS06T0677700-00 | Zinc finger CCCH domain-containing protein 45 |
| OsJ_31392 | Zinc finger CCCH domain-containing protein 62 |
| OS01T0542700-01 | cDNA clone: 002-167-H10, full insert sequence; Os01g0542700 protein; Putative uncharacterized protein OSJNBa0062A24.41; Putative uncharacterized protein OSJNBa0065J17.8; cDNA clone: 002-167-H10, full insert sequence |
| OsJ_02888 | G-box binding protein; Os01g0658900 protein; Uncharacterized protein |
| ABI5 | Os01g0859300 protein; Transcription factor that possesses transactivation activity in yeast. Involved in abscisic acid (ABA) signaling pathway. Binds to the G-box motif 5'- CACGTG-3' of TRAB1 gene promoter. Involved in the regulation of pollen maturation. May act as negative regulator of salt stress response. Together with PYL5, PP2C30 and SAPK2, is part of an ABA signaling unit that modulates seed germination and early seedling growth |

| | |
|---|---|
| LG2 | Transcription factor lg2; Transcriptional regulator involved in defense response (By similarity). Acts as transcriptional activator in vitro |
| BZIP12 | Bzip transcription factor 12; Transcription activator that binds to the ABA-responsive elements (ABREs) in vitro. Involved in abiotic stress responses and abscisic acid (ABA) signaling. Involved in the signaling pathway that induces growth inhibition in response to D- allose |
| OsJ_05262 | Os02g0132500 protein; cDNA clone: J023064O22, full insert sequence |
| BZIP23 | Bzip transcription factor 23; Transcriptional activator that mediates abscisic acid (ABA) signaling Can regulate the expression of a wide spectrum of stress-related genes in response to abiotic stresses through an ABA- dependent regulation pathway. Confers ABA-dependent drought and salinity tolerance. Binds specifically to the ABA-responsive elements (ABRE) in the promoter of target genes to mediate stress-responsive ABA signaling |
| OsJ_09015 | Os02g0833600 protein; Putative uncharacterized protein OJ1282_E10.14 |
| OsJ_13975 | OSJNBa0027O01.8 protein |
| OS05T0569300-01 | Os05g0569300 protein; Putative G-box binding factor; Putative bZIP protein |
| TGAL10 | Putative transcription factor stga1; cdna clone: 002-121-a09, full insert sequence; Transcriptional regulator involved in defense response |
| OS08T0357300-01 | Putative uncharacterized protein; Os08g0357300 protein |

| | |
|---|---|
| OS09T0306400-01 | Os09g0306400 protein; Putative bZIP transcription factor; cDNA clone: J013000L18, full insert sequence; cDNA clone: J013001A19, full insert sequence; cDNA clone: J023058F04, full insert sequence |
| OS09T0456200-01 | cDNA clone: J013049N23, full insert sequence; Os09g0456200 protein; Putative bZIP transcription factor ABI5; cDNA clone: J013049N23, full insert sequence |
| OsJ_10404 | cDNA clone: 002-141-G12, full insert sequence; Os09g0474000 protein; Putative uncharacterized protein OSJNBa0026C08.19; cDNA clone: 002-141-G12, full insert sequence |
| OS12T0156200-00 | Os12g0156200 protein; BZIP-like protein |
| OS12T0233800-02 | BZIP transcription factor family protein, expressed; cDNA clone: J013039C04, full insert sequence; cDNA clone: J023123F07, full insert sequence |
| OS01T0229000-00 | Os01g0229000 protein; MYB23 protein |
| OS01T0695900-01 | Myb-related protein 1-like; Os01g0695900 protein; cDNA clone: 002-141-A12, full insert sequence |
| OsJ_03162 | Os01g0702700 protein; Uncharacterized protein; cDNA clone: J023084H22, full insert sequence |
| OS01T0975300-02 | Putative Myb factor protein; cDNA clone: 001-040-C01, full insert sequence; cDNA clone: J013058H19, full insert sequence |
| OsJ_04971 | Os01g0977300 protein; Putative tuber-specific and sucrose-responsive element binding factor; Uncharacterized protein; cDNA clone: 002-169-B11, full insert sequence |

| | |
|---|---|
| OS02T0641300-00 | annotation not available |
| MYB2 | Transcription factor myb2; Transcription factor involved in abiotic stress responses. Plays a regulatory role in tolerance to salt, cold, and drought stresses. Regulates positively the expression of genes involved in proline synthesis and transport, and genes involved in reactive oxygen species (ROS) scavenging such as peroxidase, superoxide dismutase and catalase during salt stress. Transactivates stress-related genes, including LEA3, RAB16A and DREB2A during salt stress |
| OS03T0410000-01 | annotation not available |
| OS04T0348300-01 | Putative uncharacterized protein; Os04g0348300 protein |
| OS05T0350900-01 | Putative uncharacterized protein; Os05g0350900 protein |
| B1122D01.8 | cDNA clone: J013135D01, full insert sequence; Os05g0449900 protein; Putative uncharacterized protein B1122D01.8; cDNA clone: J013135D01, full insert sequence |
| OS05T0459000-01 | Os05g0459000 protein |
| OS05T0543600-01 | Os05g0543600 protein; Myb-related protein |
| OS06T0162700-01 | Os06g0162700 protein; MYB transcription factor-like |
| OsJ_20870 | Myb related transcription factor; Os06g0258000 protein; Putative myb factor |
| OS07T0629000-01 | Os07g0629000 protein; Putative myb protein |
| OsJ_26048 | Myb transcription factor (ATMYB4)-like protein; Os08g0151300 protein; cDNA clone: J013114D05, full insert sequence |

| | |
|---|---|
| OsJ_29287 | Os09g0401000 protein; Putative Myb51 protein |
| OsJ_30158 | cDNA clone: 006-202-H01, full insert sequence; MYB transcription factor; Os09g0538400 protein; Putative Myb-related protein Zm38; cDNA clone: 006-202-H01, full insert sequence; cDNA clone: 006-203-E04, full insert sequence; cDNA clone: 006-301-B02, full insert sequence |
| OS12T0238000-00 | Os12g0238000 protein; Myb-like DNA-binding domain containing protein, expressed; Os12g0238000 protein |
| OS12T0564100-01 | annotation not available |
| RS2 | Protein rough sheath 2 homolog; Transcription factor required for normal cell differentiation. May interact with other proteins to repress the knox homeobox genes (By similarity) |
| OsJ_00642 | Uncharacterized protein; cDNA clone: 002-150-F11, full insert sequence |
| OsJ_10658 | Os03g0321700 protein; WRKY DNA binding domain containing protein, expressed; cDNA clone: J033057D07, full insert sequence |
| OS03T0444900-00 | Os03g0444900 protein |
| OsJ_16065 | OSJNBb0015N08.8 protein; Os04g0605100 protein; cDNA clone: 006-212-C10, full insert sequence; cDNA clone: J023149N23, full insert sequence |
| OS05T0129800-01 | Os05g0129800 protein |
| OS05T0183100-01 | Os05g0183100 protein; WRKY transcription factor 67 |

| | |
|---|---|
| OS08T0276200-01 | Os08g0276200 protein |
| WRKY30 | cDNA clone: J013025I24, full insert sequence; Os08g0499300 protein; Putative WRKY DNA-binding protein; Transcription factor WRKY30; WRKY transcription factor 30; WRKY30; cDNA clone: J013025I24, full insert sequence |
| OS10T0579400-01 | annotation not available |
| OsJ_32730 | WRKY DNA binding domain containing protein, expressed |
| OS12T0102300-01 | annotation not available |
| OS12T0116400-00 | WRKY DNA binding domain containing protein, expressed |
| LFL1 | B3 domain-containing protein LFL1; Transcription repressor involved in flowering time regulation. Represses the flowering activator EHD1 by binding specifically to the DNA sequence 5'-CATGCATG-3 of its promoter |
| OS01T0723700-01 | B3 domain-containing protein Os03g0120900 |
| VP1 | B3 domain-containing protein VP1; Probable transcription factor that may participate in abscisic acid-regulated gene expression during seed development. May be required for seed maturation and dormancy induction |
| OsJ_06608 | B3 domain-containing protein Os02g0455800 |
| OS03T0164300-01 | annotation not available |
| OS03T0212300-01 | B3 domain-containing protein Os03g0212300 |
| OsJ_11751 | B3 domain-containing protein Os03g0619600 |

| | |
|---|---|
| OS03T0619700-01 | Os03g0619700 protein; Os03g0619800 protein |
| OS04T0386900-01 | B3 domain-containing protein Os04g0386900 |
| OS07T0273700-00 | Os07g0273700 protein |
| P0703C03.6 | B3 domain-containing protein Os08g0324300 |
| OS10T0323000-01 | B3 domain-containing protein Os10g0323000; Os10g0323000 protein |
| OS11T0156000-01 | B3 domain-containing protein Os11g0156000 |
| OS12T0157000-01 | Os12g0157000 protein |
| OS02T0231000-00 | annotation not available |
| OS02T0655200-01 | annotation not available |
| OsJ_07793 | Os02g0656600 protein; Putative DRE binding factor 2 |
| OS03T0191900-01 | AP2 domain containing protein, expressed; Os03g0191900 protein |
| OS03T0815800-01 | cDNA clone: J013145D23, full insert sequence; AP2 domain containing protein, expressed; Os03g0815800 protein; Putative AP2 domain containing protein; Putative AP2 domain transcription factor; cDNA clone: J013145D23, full insert sequence |
| OS04T0550200-01 | cDNA clone: 002-143-E08, full insert sequence; OSJNBb0034G17.6 protein; Os04g0550200 protein; Transcription factor; cDNA clone: 002-143-E08, full insert sequence |
| OS04T0655700-00 | annotation not available |

| | |
|---|---|
| P0692D12.5 | cDNA clone: J023031H12, full insert sequence; Os05g0361700 protein; Putative uncharacterized protein P0530H10.12; Putative uncharacterized protein P0692D12.5; cDNA clone: J023031H12, full insert sequence |
| OS05T0437100-00 | Os05g0437100 protein; Os05g0437133 protein |
| P0648E08.7 | EREBP transcription factor; EREBP-like protein; Os06g0194000 protein; Putative ethylene responsive element binding factor; cDNA clone: 001-032-E01, full insert sequence; cDNA clone: J023031B06, full insert sequence |
| OS06T0553700-00 | Os06g0553700 protein; Putative Ap21 |
| OsJ_25135 | EREB-like protein; Os07g0617000 protein; cDNA clone: 006-205-F01, full insert sequence; cDNA clone: 006-206-B03, full insert sequence; cDNA clone: J013090P08, full insert sequence |
| OJ1081_B12.108 | cDNA clone: 002-140-D07, full insert sequence; Os08g0521600 protein; Putative uncharacterized protein OJ1003_A09.37; Putative uncharacterized protein OJ1081_B12.108; cDNA clone: 002-140-D07, full insert sequence |
| ERF44 | Dehydration-responsive element-binding protein 2C; Probable transcriptional activator that binds to the DNA sequence 5'-[AG]CCGAC-3' of the cis-acting dehydration-responsive element (DRE) |
| OsJ_28694 | cDNA clone: 001-206-F09, full insert sequence; AP2/ERF domain protein; Ethylene responsive protein-like; Os09g0287000 protein; cDNA clone: 001-206-F09, full insert sequence; cDNA clone: J013163J01, full insert sequence |
| OS10T0560700-00 | annotation not available |
| OS01T0187900-01 | Os01g0187900 protein; Putative MybSt1; cDNA clone: J023105I20, full insert sequence |

| | |
|---|---|
| OsJ_02512 | cDNA clone: J033128P13, full insert sequence; Putative MCB2 protein |
| OS01T0619900-01 | annotation not available |
| OS03T0425800-01 | Os03g0425800 protein |
| OS04T0583900-01 | OSJNBa0088A01.20 protein; Os04g0583900 protein |
| OS05T0114700-02 | Os05g0114700 protein |
| OS05T0589400-01 | cDNA clone: 006-205-B05, full insert sequence; MYB-related transcription factor; Os05g0589400 protein; Putative myb transcription factor; cDNA clone: 006-205-B05, full insert sequence; cDNA clone: J013154K05, full insert sequence |
| OS07T0695900-01 | Os07g0695900 protein; Putative uncharacterized protein P0627E10.29 |
| OS10T0443800-01 | Os10g0443800 protein |
| MYBS3 | Transcription factor mybs3; Transcription repressor that binds to 5'-TATCCA-3' elements in gene promoters. Contributes to the sugar-repressed transcription of promoters containing SRS or 5'-TATCCA-3' elements. Transcription repressor involved in a cold stress response pathway that confers cold tolerance. Suppresses the DREB1-dependent signaling pathway under prolonged cold stress. DREB1 responds quickly and transiently while MYBS3 responds slowly to cold stress. They may act sequentially and complementarily for adaptation to short- and long-term cold stress |

| | |
|---|---|
| MYBS2 | MCB1 protein, putative, expressed; Os10g0562100 protein; Putative Myb-related protein; Transcription activator that binds to 5'-TATCCA-3' elements in gene promoters. Derepresses weakly the sugar-repressed transcription of promoters containing SRS. Contributes to the sugar-repressed transcription of promoters containing 5'-TATCCA-3' elements |
| MYBAS2 | Myb-related protein MYBAS2; Transcription factor |
| OS01T0191300-01 | Os01g0191300 protein; NAC-type transcription factor |
| OS01T0393100-01 | NAC protein; Os01g0393100 protein; Putative OsNAC2 |
| OS01T0672100-02 | Os01g0672100 protein |
| NAC6 | NAC domain-containing protein 48; Transcription activator that binds to the promoter of the stress response gene LEA19. Involved in tolerance to abiotic stresses. Transcription activator involved in response to abiotic and biotic stresses. Involved in cold and salt stress responses, and defense response to the rice blast fungus. Transcription activator involved tolerance to cold and salt stresses. Transcription activator involved in tolerance to drought stress. Targets directly and activates genes involved in membrane modification, nicotianamine (NA) biosynthesis, glutathione relocatio [...] |
| OS01T0925400-01 | NAC4 protein; Os01g0925400 protein; Putative development regulation gene OsNAC4; cDNA clone: J013106M04, full insert sequence; cDNA clone: J013153H06, full insert sequence |
| P0660F12.31 | cDNA clone: J023080K20, full insert sequence; Os01g0946200 protein; P0660F12.31 protein; Putative uncharacterized protein P0614D08.37; cDNA clone: J023080K20, full insert sequence |

| | |
|---|---|
| OsJ_05880 | NAC6; Os02g0214500 protein; Putative OsNAC6 protein; cDNA clone: 002-126-B03, full insert sequence |
| NAC58 | NAC domain transcription factor, putative, expressed; Putative NAM (No apical meristem) protein; cDNA, clone: J100075D15, full insert sequence |
| OsJ_12786 | NAC-domain containing protein 9, putative, expressed; Os03g0777000 protein; Putative NAC domain protein; Putative NAM (No apical meristem) protein |
| OS05T0421600-01 | Os05g0421600 protein |
| NAC3 | NAC domain-containing protein 67; Probable transcription factor involved in stress response |
| NAC010 | NAC domain-containing protein 10; Transcription factor of the NAC family associated with male fertility. Involved in anther development, but not in senescence. Reduced expression of NAC5 via RNAi leads to male- sterility |
| OS08T0103900-01 | cDNA clone: 002-168-H04, full insert sequence; Os08g0103900 protein; Putative OsNAC7 protein; Secondary wall NAC transcription factor 3; cDNA clone: 002-168-H04, full insert sequence |
| P0470F10.1 | Os08g0115800 protein; Putative OsNAC7 protein |
| OsJ_26109 | NAC2 protein; NAC2 protein-like; Os08g0157900 protein |
| OS08T0433500-00 | Os08g0433500 protein |

| | |
|---|---|
| OsJ_28085 | cDNA clone: 001-038-H11, full insert sequence; NAC4 protein; Os08g0535800 protein; Putative development regulation gene OsNAC4; cDNA clone: 001-038-H11, full insert sequence; cDNA clone: 006-311-A02, full insert sequence; cDNA clone: J013160K11, full insert sequence |
| OS09T0552900-00 | annotation not available |
| OS10T0571600-01 | Os10g0571600 protein |
| NAC5 | NAC domain-containing protein 71; Transcription activator that binds to the promoter of the stress response gene LEA19. Involved in tolerance to abiotic stresses |
| OsJ_34014 | NAM protein; No apical meristem protein; cDNA, clone: J075106F03, full insert sequence |
| OS11T0512600-00 | Os11g0512600 protein; NAM protein |
| GLK2 | Probable transcription factor GLK2; Probable transcriptional activator that promotes chloroplast development. Acts as an activator of nuclear photosynthetic genes involved in chlorophyll biosynthesis, light harvesting, and electron transport (By similarity) |
| OS02T0174000-00 | annotation not available |
| OS02T0700300-01 | Os02g0700300 protein; cDNA clone: 001-028-A10, full insert sequence |
| OS03T0764600-01 | Myb-like DNA-binding domain, SHAQKYF class family protein, expressed; Os03g0764600 protein; Putative Myb-like DNA-binding protein; cDNA clone: 001-200-C11, full insert sequence |

| | |
|---|---|
| OS04T0665600-01 | Os04g0665600 protein |
| GLK1 | Probable transcription factor GLK1; Probable transcriptional activator that promotes chloroplast development. Acts as an activator of nuclear photosynthetic genes involved in chlorophyll biosynthesis, light harvesting, and electron transport |
| OS06T0664800-01 | Myb family transcription factor-like |
| OsJ_22904 | cDNA clone: J023048M12, full insert sequence; Os07g0119300 protein; Putative cytoskeletal protein-like protein; cDNA clone: J023048M12, full insert sequence |
| OsJ_26993 | cDNA clone: J033149L11, full insert sequence; Putative uncharacterized protein P0404D10.1-2; Putative uncharacterized protein P0410E11.132-2 |
| P0404D10.4 | Os08g0346500 protein; Putative transfactor |
| OS08T0434700-01 | Os08g0434700 protein |
| OS11T0106100-00 | annotation not available |
| OsJ_34927 | Myb-like DNA-binding domain, SHAQKYF class family protein, expressed; Os12g0105600 protein |
| OS03T0769800-01 | Homeobox domain containing protein, expressed; Os03g0769800 protein; Putative homeodomain leucine-zipper protein Hox9; cDNA clone: J023060H19, full insert sequence |
| OS05T0320300-01 | annotation not available |
| OS11T0163500-01 | Expressed protein; Os11g0163500 protein; cDNA clone: J033027D16, full insert sequence |

| | |
|---|---|
| OS02T0104500-01 | Os02g0104500 protein; Putative GT-2 factor |
| OS02T0174300-00 | Putative uncharacterized protein OSJNBa0073A21.24 |
| OS02T0516800-00 | annotation not available |
| OsJ_07175 | Os02g0565000 protein; Putative 6b-interacting protein 1 |
| OS02T0648300-01 | Putative DNA-binding protein Gt-2 |
| OS04T0397500-00 | Putative uncharacterized protein; Os04g0397500 protein |
| OS04T0541100-01 | Os04g0541100 protein |
| ZHD7 | Putative zf-hd homeobox protein; putative uncharacterized protein 49d11.22; Putative transcription factor |
| ZHD10 | Os08g0438400 protein; Putative ZF-HD homeobox protein; Putative transcription factor |
| ZHD2 | Os08g0479400 protein; Putative ZF-HD homeobox protein; cDNA clone: 002-116-C04, full insert sequence; Putative transcription factor |
| ZHD1 | Os09g0466400 protein; Putative ZF-HD homeobox protein; cDNA clone: 002-140-H06, full insert sequence; Putative transcription factor |
| MIF1 | Mini zinc finger protein 1; Inhibits zinc finger homeodomain (ZHD) transcription factors, by interacting with them to prevent both their nuclear localization and their DNA-binding properties |

| | |
|---|---|
| MIF2 | Mini zinc finger protein 2; Inhibits zinc finger homeodomain (ZHD) transcription factors, by interacting with them to prevent both their nuclear localization and their DNA-binding properties |
| OsJ_02811 | Uncharacterized protein; cDNA, clone: J080076J05, full insert sequence; Belongs to the GRAS family |
| P0406G08.8 | Os01g0842200 protein; SCARECROW-like protein; Belongs to the GRAS family |
| GAI | DELLA protein SLR1; Probable transcriptional regulator that acts as a repressor of the gibberellin (GA) signaling pathway. Probably acts by participating in large multiprotein complexes that repress transcription of GA-inducible genes. Upon GA application, it is degraded by the proteasome, allowing the GA signaling pathway. In contrast, its overexpression prevents the GA signaling pathway and induces a dwarf phenotype; Belongs to the GRAS family. DELLA subfamily |
| OS06T0610300-01 | Os06g0610300 protein; Putative uncharacterized protein P0490F09.18 |
| OS10T0369600-01 | Os10g0369600 protein; Chitin-inducible gibberellin-responsive protein 2, putative, expressed |
| MADS3 | Mads-box transcription factor 3; Probable transcription factor involved in the development of floral organs. Acts as C-class protein in association with MADS58. Involved in the control of lodicule number (whorl 2), stamen specification (whorl 3) and floral meristem determinacy (whorl 4), but not in the regulation of carpel morphogenesis. Plays a more predominant role in controlling lodicule development and in specifying stamen identity than MADS58 |
| OsJ_04312 | MADS-box transcription factor 2; Probable transcription factor involved in the development of floral organs. B-class protein required for normal development of lodicules (whorl 2) |

| | |
|---|---|
| MADS21 | MADS-box transcription factor 21; Probable transcription factor |
| OS02T0104100-00 | MADS-box protein PTM5-like |
| MADS29 | MADS-box transcription factor 29; Probable transcription factor |
| MADS27 | MADS-box transcription factor 27; Probable transcription factor |
| MADS6 | MADS-box transcription factor 6; Probable transcription factor. Regulates floral organ identity and floral meristem determinacy. May be involved in the control of flowering time |
| MADS1 | MADS-box transcription factor 1; Probable transcription factor involved in the development of floral organs. Required for the formation of inner floral organs (lodicules, stamens and carpels, or whorls 2, 3 and 4) and the lemma and palea (whorl 1), which are grass floral organs analogous to sepals. May be involved in the control of flowering time. Seems to act as transcriptional activator. May act upstream of the auxin-responsive protein GH3.8 |
| MADS17 | MADS-box transcription factor 17; Probable transcription factor. Plays minor but redundant roles with MADS6 in floral development |
| MADS31 | MADS-box transcription factor 31; Probable transcription factor |
| MADS58 | MADS-box transcription factor 58; Probable transcription factor involved in the development of floral organs. Acts as a C-class protein in association with MADS3. Involved in the control of lodicule number (whorl 2), stamen specification (whorl |

| | |
|---|---|
| | 3), floral meristem determinacy and regulation of the carpel morphogenesis (whorl 4). Plays a more predominant role in floral meristem determinacy than MADS3 |
| MADS4 | MADS-box transcription factor 4; Probable transcription factor involved in the development of floral organs. B-class protein required for normal development of lodicules and stamens (whorls 2 and 3). May function as a heterodimer with MADS16 |
| MADS5 | MADS-box transcription factor 5; Probable transcription factor. May be involved in the control of flowering time |
| MADS16 | MADS-box transcription factor 16; Probable transcription factor involved in the development of floral organs. Required for normal development of lodicules and stamens (whorls 2 and 3). May function as a heterodimer with MADS4 |
| MADS15 | MADS-box transcription factor 15; Probable transcription factor |
| MADS26 | MADS-box transcription factor 26; Probable transcription factor |
| M79 | MADS-box transcription factor 7; Probable transcription factor. May be involved in the control of flowering time |
| MADS8 | MADS-box transcription factor 8; Probable transcription factor. May be involved in the control of flowering time |
| MADS56 | MADS-box transcription factor 56; Probable transcription factor |
| MADS13 | MADS-box transcription factor 13; Probable transcription factor |
| WOX11 | WUSCHEL-related homeobox 11; Transcription factor which may be involved in developmental processes (By similarity). Promotes the development of crown roots (both initiation and elongation), main components of the fibrous root system, by regulating the expression of genes required for crown root development and hormone-responsive genes involved in cytokinin |

| | |
|---|---|
| | (e.g. RR1, RR2, RR3 and RR4) and auxin (e.g. IAA5, IAA11, IAA23 and IAA31) signaling; Belongs to the WUS homeobox family |
| OS01T0511000-01 | Os01g0511000 protein |
| OS03T0659700-01 | Os03g0659700 protein |
| OS07T0589000-01 | Os07g0589000 protein; Putative lateral organ boundaries (LOB) domain protein 37; cDNA clone: J023030F23, full insert sequence |
| OS02T0610500-01 | Os02g0610500 protein; Putative COL1 protein; cDNA clone: J023004D12, full insert sequence |
| HD1 | Zinc finger protein HD1; Probable transcription factor involved in the regulation of flower development. Required for the promotion of flowering under short day (SD) conditions and the suppression of flowering under long day (LD) conditions. Regulates positively the floral activator HEADING DATE 3a (HD3A) under SD and negatively under LD conditions; Belongs to the CONSTANS family |
| OS06T0298200-01 | cDNA clone: J013001A08, full insert sequence |
| OsJ_36996 | Associated with HOX family protein, expressed; Os12g0636200 protein; cDNA clone: J023054P21, full insert sequence |
| OsJ_12119 | Associated with HOX family protein, expressed; Os03g0680800 protein; Putative homeodomain protein; cDNA clone: J013124I05, full insert sequence |
| OS11T0158600-00 | Associated with HOX family protein, expressed; Os11g0158600 protein |

| | |
|---|---|
| qSH1 | Putative transcription factor qSH-1; QSH-1 |
| OS03T0124000-01 | Associated with HOX family protein, expressed; Os03g0124000 protein |
| OSH1 | Homeobox protein knotted-1-like 6; Transcription factor that regulates genes involved in development. May be involved in shoot formation during embryogenesis. Overexpression in transgenic plants causes altered leaf morphology. Regulates anther dehiscence via direct repression of the auxin biosynthetic gene YUCCA4. Binds to the DNA sequence 5'-TGAC-3' in the promoter of the YUCCA4 gene and represses its activity during anther development. Reduction of auxin levels at late stage of anther development, after meiosis of microspore mother cells, is necessary for normal anther dehiscence [...] |
| OSH15 | Homeobox protein knotted-1-like 12; Probable transcription factor that may be involved in shoot formation during embryogenesis; Belongs to the TALE/KNOX homeobox family |
| OSH6 | Homeobox protein knotted-1-like 1; Probable transcription factor that may be involved in shoot formation during early embryogenesis |
| OSH71 | Homeobox protein knotted-1-like 10; Probable transcription factor that may be involved in shoot formation during embryogenesis |
| OSH3 | Homeobox protein knotted-1-like 7; Probable transcription factor that may be involved in shoot formation during embryogenesis; Belongs to the TALE/KNOX homeobox family |
| HOS59 | Homeobox protein knotted-1-like 11 |

| | |
|---|---|
| HOS58 | Homeobox protein knotted-1-like 2 |
| HOS66 | Homeobox protein knotted-1-like 3; Belongs to the TALE/KNOX homeobox family |
| OS01T0273200-00 | annotation not available |
| OS02T0284500-01 | Os02g0284500 protein; Putative far-red impaired response protein |
| OS02T0542200-01 | Putative far-red impaired response protein |
| OS02T0550900-02 | Os02g0550900 protein; Putative far-red impaired response protein; cDNA clone: J033108J05, full insert sequence |
| OS03T0164400-01 | Transposon protein, putative, unclassified, expressed; cDNA clone: 001-114-D10, full insert sequence |
| OS03T0181600-02 | Os03g0181600 protein; Transposon protein, putative, unclassified, expressed; cDNA clone: J013120L22, full insert sequence |
| OsJ_10174 | FAR1 family protein, expressed; Os03g0255400 protein; cDNA clone: J013072M14, full insert sequence |
| OS05T0317300-01 | annotation not available |
| LOL4 | Protein LOL4; Putative zinc finger that may be involved in programmed cell death and defense response |
| PCF6 | Transcription factor PCF6; Transcription activator. Binds the promoter core sequence 5'-GGNCC-3' |
| PCF2 | Transcription factor PCF2; Transcription activator. Binds the promoter core sequence 5'-GGNCC-3', especially at sites IIa (5'-GGGCCCAC-3') and IIb (5'-GGTCCCAC-3') (essential for meristematic tissue- specificity expression) of the PCNA gene promoter |

| | |
|---|---|
| PCF8 | Transcription factor PCF8; Transcription activator. Binds the promoter core sequence 5'-GGNCC-3' |
| YAB2 | Protein YABBY 2; Belongs to the YABBY family |
| EIL1A | ETHYLENE-INSENSITIVE3-like 1 protein, putative, expressed; Transcription factor acting as a positive regulator in the ethylene response pathway Required for the inhibition of root growth by ethylene in etiolated seedlings Functions upstream of the auxin biosynthetic gene YUCCA8 and directly activates its expression. Functions downstream of the ethylene signaling factor EIN2 in disease resistance against the rice blast fungus (Magnaporthe oryzae). Binds directly to the promoters of the NADPH oxidases RBOHA and RBOHB, and the jasmonate biosynthetic gene OPR4 to activate their expressio [...] |
| OS04T0456900-00 | annotation not available |
| OsEIL2 | cDNA clone: 001-033-G01, full insert sequence; EIL transcription factor; Ethylene-insensitive-3-like protein; Os07g0685700 protein; Putative transcription factor OsEIL2; cDNA clone: 001-033-G01, full insert sequence |
| OS08T0508700-01 | Os08g0508700 protein; Putative ethylene-insensitive protein (EIL); cDNA clone: J023115E06, full insert sequence |
| OS12T0605500-01 | Os12g0605500 protein; Tesmin/TSO1-like CXC domain containing protein, expressed; cDNA clone: J033075J10, full insert sequence |
| OS02T0606200-01 | cDNA clone: 001-007-G06, full insert sequence; ORPHAN transcription factor; Os02g0606200 protein; Zinc finger protein; cDNA clone: 001-007-G06, full insert sequence; cDNA clone: 006-205-E01, full insert sequence |

| | |
|---|---|
| OsJ_15063 | Os04g0461300 protein |
| OS09T0116800-00 | MADS-box protein-like |
| OS01T0343300-01 | Os01g0343300 protein; Putative uncharacterized protein B1045F02.28 |
| SPL3 | Squamosa promoter-binding-like protein 3; Trans-acting factor that binds specifically to the consensus nucleotide sequence 5'-TNCGTACAA-3' (By similarity). May be involved in panicle development |
| OS12T0158800-01 | Os12g0158800 protein; Transcription factor E2F/dimerisation partner family protein, expressed |
| OS03T0191000-01 | Os03g0191000 protein |
| HSFA6B | Heat stress transcription factor A-6a; Transcriptional regulator that specifically binds DNA of heat shock promoter elements (HSE) |
| OsJ_34182 | CCAAT transcription factor; Histone-like transcription factor and archaeal histone family protein, expressed; Os11g0544700 protein; cDNA clone: J023143I16, full insert sequence |
| GA3OX2 | Os01g0177400 protein; Belongs to the iron/ascorbate-dependent oxidoreductase family |
| OS01T0188100-01 | Putative oxidoreductase; cDNA, clone: J100072I17, full insert sequence; Belongs to the iron/ascorbate-dependent oxidoreductase family |

| | |
|---|---|
| OsGA2ox3 | Gibberellin 2-oxidase; Os01g0757200 protein; cDNA clone: 001-028-C12, full insert sequence; cDNA clone: 001-031-D01, full insert sequence; cDNA clone: J033060N07, full insert sequence; Belongs to the iron/ascorbate-dependent oxidoreductase family |
| OS02T0103700-01 | Os02g0103700 protein; cDNA clone: 001-104-A12, full insert sequence; cDNA clone: J033086N17, full insert sequence |
| OS02T0567800-01 | Os02g0567800 protein; Putative PrMC3 |
| OS03T0252100-01 | Os03g0252100 protein; Esterase, putative, expressed; Os03g0252100 protein; cDNA, clone: J065073H04, full insert sequence |
| OsJ_11688 | cDNA clone: J023044E14, full insert sequence; Gibberellin regulated protein; Os03g0607200 protein; Putative gibberellin regulated protein; cDNA clone: J023044E14, full insert sequence |
| OS03T0618300-01 | Os03g0618300 protein; Oxidoreductase, 2OG-Fe oxygenase family protein, expressed; Putative oxidoreductase; cDNA clone: J033026A10, full insert sequence; Belongs to the iron/ascorbate-dependent oxidoreductase family |
| DAO | 2-oxoglutarate-dependent dioxygenase DAO; 2-oxoglutarate-dependent dioxygenase essential for auxin catabolism and maintenance of auxin homeostasis in reproductive organs. Catalyzes the irreversible oxidation of indole-3-acetic acid (IAA) to the biologically inactive 2-oxoindole-3-acetic acid (OxIAA) |
| OsJ_15497 | OSJNBa0019D11.23 protein; Os04g0522500 protein; Belongs to the iron/ascorbate-dependent oxidoreductase family |

| | |
|---|---|
| GA2OX1 | Gibberellin 2-beta-dioxygenase 1; Catalyzes the 2-beta-hydroxylation of several biologically active gibberellins, leading to the homeostatic regulation of their endogenous level. Catabolism of gibberellins (GAs) plays a central role in plant development. Controls the level of bioactive GAs in the shoot apical meristem, which regulates the vegetative to reproductive phase transition. In vitro, converts GA1, GA4, GA9, GA20, and GA44 to the corresponding 2-beta-hydroxylated products GA8, GA34, GA51, GA29, and GA98, respectively |
| GA3OX1 | Os05g0178100 protein; Putative gibberellin 3 beta-hydroxylase; cDNA clone: J023034H12, full insert sequence; Belongs to the iron/ascorbate-dependent oxidoreductase family |
| OsJ_18341 | cDNA clone: 002-115-G05, full insert sequence; Os05g0376800 protein; Putative gibberellin-induced protein; Putative uncharacterized protein OSJNBa0039A21.1; cDNA clone: 002-115-G05, full insert sequence |
| OS06T0214800-01 | Os06g0214800 protein; Putative PrMC3; cDNA clone: 006-204-G08, full insert sequence; cDNA clone: 006-303-C01, full insert sequence; cDNA clone: J023047F19, full insert sequence |
| OS06T0214850-00 | Os06g0214850 protein; Putative PrMC3 |
| OS06T0306600-01 | Os06g0306600 protein; Putative esterase; cDNA clone: 001-206-H11, full insert sequence |
| OsJ_22737 | cDNA clone: 001-029-A05, full insert sequence; Os06g0729400 protein; Putative uncharacterized protein OSJNBa0069C14.13; cDNA clone: 001-029-A05, full insert sequence |

| | |
|---|---|
| OS07T0162400-01 | cDNA clone: 006-306-D11, full insert sequence; Os07g0162400 protein; Putative cell death associated protein; cDNA clone: 006-306-D11, full insert sequence |
| OS07T0162500-02 | cDNA clone: 001-206-A09, full insert sequence; Os07g0162500 protein; Putative cell death associated protein; cDNA clone: 001-206-A09, full insert sequence; cDNA clone: 001-207-G10, full insert sequence; cDNA clone: 006-310-E02, full insert sequence |
| OS07T0162900-01 | Os07g0162900 protein; Putative cell death associated protein; cDNA clone: 002-100-G01, full insert sequence |
| OS07T0592000-01 | Os07g0592000 protein; cDNA clone: 002-128-B02, full insert sequence |
| OS07T0643400-01 | Os07g0643400 protein; Putative esterase; cDNA clone: 006-204-B04, full insert sequence; cDNA clone: 006-310-B02, full insert sequence |
| OS08T0560000-01 | Os08g0560000 protein; Putative iron deficiency protein Ids3; cDNA clone: 002-124-A12, full insert sequence; cDNA clone: 002-125-E03, full insert sequence; Belongs to the iron/ascorbate-dependent oxidoreductase family |
| OS09T0245500-01 | Os09g0245500 protein; Putative 2-oxoacid-dependent oxidase; Belongs to the iron/ascorbate-dependent oxidoreductase family |
| OsJ_29621 | Os09g0455900 protein; Putative PrMC3; cDNA clone: 006-204-A12, full insert sequence |
| OsJ_29649 | Os09g0461500 protein; Putative PrMC3; cDNA clone: 001-115-A02, full insert sequence; cDNA clone: J033090L22, full insert sequence |

| | |
|---|---|
| OsJ_32203 | Os10g0522900 protein; Oxidoreductase, 2OG-Fe oxygenase family protein, expressed; Putative anthocyanidin hydroxylase; Putative gibberellin 20-oxidase; cDNA clone: J013151N02, full insert sequence; Belongs to the iron/ascorbate-dependent oxidoreductase family |
| OS11T0240600-01 | cDNA clone: 001-021-G12, full insert sequence; Expressed protein; Os11g0240600 protein; PrMC3, putative, expressed; cDNA clone: 001-021-G12, full insert sequence |
| OsJ_03798 | Glycosyltransferase; Os01g0805400 protein; Putative UDP-glucose: salicylic acid glucosyltransferase; Uncharacterized protein; cDNA clone: 001-205-D05, full insert sequence; Belongs to the UDP-glycosyltransferase family |
| OsJ_06788 | Glycosyltransferase; Os02g0490500 protein; Putative UDP-glycosyltransferase; cDNA clone: J013153D03, full insert sequence; Belongs to the UDP-glycosyltransferase family |
| RR2 | Two-component response regulator orr2; Functions as response regulator involved in His-to-Asp phosphorelay signal transduction system. Phosphorylation of the Asp residue in the receiver domain activates the ability of the protein to promote the transcription of target genes. Type-A response regulators seem to act as negative regulators of the cytokinin signaling |
| OS02T0755900-01 | Glycosyltransferase; Os02g0755900 protein; Putative UDP-glucose glucosyltransferase1; cDNA clone: 002-107-F12, full insert sequence; cDNA clone: J033052B15, full insert sequence; Belongs to the UDP-glycosyltransferase family |

| | |
|---|---|
| OS04T0320700-02 | Glycosyltransferase; OSJNBa0041M06.3 protein; OSJNBb0026L04.11 protein; Os04g0320700 protein; cDNA clone: 001-203-C07, full insert sequence; cDNA clone: 001-205-E07, full insert sequence; Belongs to the UDP-glycosyltransferase family |
| OS04T0523600-01 | Glycosyltransferase; OSJNBb0065J09.9 protein; Os04g0523600 protein; Belongs to the UDP-glycosyltransferase family |
| RR6 | Two-component response regulator orr6; Functions as response regulator involved in His-to-Asp phosphorelay signal transduction system. Phosphorylation of the Asp residue in the receiver domain activates the ability of the protein to promote the transcription of target genes. Type-A response regulators seem to act as negative regulators of the cytokinin signaling |
| OS06T0729800-01 | annotation not available |
| OS08T0358800-01 | Os08g0358800 protein; TypeA response regulator 13 |
| OS11T0143300-02 | Os11g0143300 protein; Two-component response regulator ARR8, putative, expressed |
| CCD8A | Carotenoid cleavage dioxygenase 8 homolog A, chloroplastic; May be involved in strigolactones biosynthesis; Belongs to the carotenoid oxygenase family |
| OsJ_07653 | ABA-responsive protein-like; Os02g0636700 protein; cDNA clone: J023031H22, full insert sequence |
| CYP707A5 | Abscisic acid 8'-hydroxylase 1; Involved in the oxidative degradation of abscisic acid; Belongs to the cytochrome P450 family |

| | |
|---|---|
| NCED1 | 9-cis-epoxycarotenoid dioxygenase NCED1, chloroplastic; Has a 11,12(11',12') 9-cis epoxycarotenoid cleavage activity. Catalyzes the first step of abscisic-acid biosynthesis from carotenoids |
| OsJ_08376 | Os02g0747500 protein; Putative ABA-responsive protein; cDNA clone: 002-143-F05, full insert sequence |
| OsJ_09302 | HVA22-like protein i, putative, expressed; Os03g0132300 protein; cDNA clone: J013008G03, full insert sequence |
| SAPK1 | Serine/threonine-protein kinase SAPK1; May play a role in signal transduction of hyperosmotic response |
| SAPK10 | Serine/threonine-protein kinase SAPK10; May play a role in signal transduction of hyperosmotic response; Belongs to the protein kinase superfamily. Ser/Thr protein kinase family |
| NCED3 | 9-cis-epoxycarotenoid dioxygenase 3; Os03g0645900 protein; Viviparous-14, putative, expressed; Has a 11,12(11',12') 9-cis epoxycarotenoid cleavage activity. Catalyzes the first step of abscisic-acid biosynthesis from carotenoids |
| OS03T0736700-03 | cDNA clone: 006-201-F03, full insert sequence; GRAM domain containing protein, expressed; Os03g0736700 protein; Putative ABA-responsive protein; cDNA clone: 006-201-F03, full insert sequence; cDNA clone: 006-204-F03, full insert sequence; cDNA clone: 006-310-F11, full insert sequence |
| OS03T0790700-01 | Probable aldehyde oxidase 3 |
| OS03T0790900-01 | Probable aldehyde oxidase 2 |
| OsJ_14752 | HVA22-like protein; Os04g0415200 protein; cDNA clone: 002-142-E03, full insert sequence |
| SAPK7 | Serine/threonine-protein kinase SAPK7; May play a role in signal transduction of hyperosmotic response |

| | |
|---|---|
| ZEP | Zeaxanthin epoxidase, chloroplastic; Zeaxanthin epoxidase that plays an important role in the xanthophyll cycle and abscisic acid (ABA) biosynthesis. Converts zeaxanthin into antheraxanthin and subsequently violaxanthin. Required for resistance to osmotic and drought stresses, seed development and dormancy |
| CCD7 | Carotenoid cleavage dioxygenase 7, chloroplastic; Involved in strigolactones biosynthesis by cleaving asymmetrically a variety of linear and cyclic carotenoids at the 9-10 double bond. Produces one C(13) beta-ionone and the C(27) 10'-apo-beta-carotenal. Strigolactones are hormones that inhibit tillering and shoot branching through the MAX-dependent pathway, contribute to the regulation of shoot architectural response to phosphate-limiting conditions and function as rhizosphere signal that stimulates hyphal branching of arbuscular mycorrhizal fungi and trigger seed germination of root p [...] |
| OS07T0164900-01 | cDNA clone: J033131J21, full insert sequence |
| OS08T0467500-02 | Os08g0467500 protein; Putative abscisic acid-induced protein |
| OS10T0177400-02 | HVA22-like protein; Os10g0177400 protein; TB2/DP1, HVA22 family protein, expressed |
| SAPK3 | Serine/threonine-protein kinase SAPK3; May play a role in signal transduction of hyperosmotic response |
| OsJ_35292 | HVA22-like protein |
| NCED5 | 9-cis-epoxycarotenoid dioxygenase NCED5, chloroplastic; Has a 11,12(11',12') 9-cis epoxycarotenoid cleavage activity. Catalyzes the first step of abscisic-acid biosynthesis from carotenoids |
| GT3 | Glutelin type-A 3; Seed storage protein |

| | |
|---|---|
| EMP1 | Embryonic abundant protein 1; Em protein may act as a cytoplasm protectant during desiccation; Belongs to the small hydrophilic plant seed protein family |
| OS08T0127900-01 | Os08g0127900 protein; Putative early embryogenesis protein; cDNA clone: 001-117-E07, full insert sequence; cDNA clone: 002-130-H05, full insert sequence |
| MODD | Ninja-family protein MODD; Acts as negative regulator of abscisic acid (ABA) signaling and drought tolerance. Mediates deactivation and degradation of BZIP46, a positive regulator of ABA signaling and drought stress tolerance. Represses BZIP46 activity via interaction with the TPR3-HDAC1 corepressor complex and down- regulation of the histone acetylation level at BZIP46 target genes. Promotes BZIP46 degradation via interaction with the U-box type ubiquitin E3 ligase PUB70 |
| DGAT1-2 | Diacylglycerol o-acyltransferase 1-2; Involved in triacylglycerol (TAG) synthesis. Catalyzes the acylation of the sn-3 hydroxy group of sn-1,2-diacylglycerol using acyl-CoA |
| OS10T0505900-01 | Expressed protein; Os10g0505900 protein |
| OS01T0159600-01 | cDNA clone: 001-125-F04, full insert sequence; Os01g0159600 protein; Putative uncharacterized protein P0041E11.22; cDNA clone: 001-125-F04, full insert sequence; cDNA clone: J033143J06, full insert sequence |
| OS12T0183300-01 | 3'(2'),5'-bisphosphate nucleotidase; Converts adenosine 3'-phosphate 5'-phosphosulfate (PAPS) to adenosine 5'-phosphosulfate (APS) and 3'(2')-phosphoadenosine 5'- phosphate (PAP) to AMP. Regulates the flux of sulfur in the sulfur-activation pathway by converting PAPS to APS (By similarity); Belongs to the inositol monophosphatase superfamily |

| | |
|---|---|
| OS02T0456100-01 | annotation not available |
| OsJ_32040 | cDNA clone: J023096B06, full insert sequence; Epoxide hydrolase, putative, expressed; Os10g0498000 protein; Putative epoxide hydrolase; cDNA clone: J023096B06, full insert sequence |
| OS02T0596700-00 | annotation not available |
| ASA1 | Anthranilate synthase alpha subunit 1, chloroplastic; Part of a heterotetrameric complex that catalyzes the two- step biosynthesis of anthranilate, an intermediate in the biosynthesis of L-tryptophan. In the first step, the glutamine-binding beta subunit of anthranilate synthase (AS) provides the glutamine amidotransferase activity which generates ammonia as a substrate that, along with chorismate, is used in the second step, catalyzed by the large alpha subunit of AS to produce anthranilate |
| OsJ_18684 | cDNA clone: J023104C20, full insert sequence; Actin-7; Belongs to the actin family |
| OS05T0481900-01 | cDNA clone: 002-115-B08, full insert sequence; Os05g0481900 protein; Putative uncharacterized protein OSJNBa0095J22.10; cDNA clone: 002-115-B08, full insert sequence |
| OsJ_35204 | Os12g0145100 protein; Seed developmental stage protein; Stem-specific protein TSJT1, putative, expressed; cDNA clone: 001-034-F10, full insert sequence; cDNA clone: 006-202-B06, full insert sequence; cDNA clone: J013095J21, full insert sequence |

| | |
|---|---|
| OsJ_21850 | Os06g0592500 protein; Putative ethylene-responsive transcriptional coactivator; cDNA clone: 002-157-H08, full insert sequence; cDNA clone: J033114G13, full insert sequence |
| OsJ_20326 | Os06g0178100 protein; Putative iron/ascorbate-dependent oxidoreductase; Belongs to the iron/ascorbate-dependent oxidoreductase family |
| OS06T0182100-00 | Apoptosis-related protein PNAS-4 like; Os06g0182100 protein |
| OS07T0471900-00 | Os07g0471900 protein |
| OsJ_27030 | Os08g0356500 protein; Putative uncharacterized protein B1104G07.27; Putative uncharacterized protein P0426E02.5 |
| OsJ_27077 | Os08g0366100 protein; Putative ethylene-responsive transcriptional coactivator; cDNA clone: 001-041-F01, full insert sequence; cDNA clone: J013060M10, full insert sequence |
| OS10T0558900-01 | Putative uncharacterized protein; Os10g0558900 protein; Belongs to the iron/ascorbate-dependent oxidoreductase family |
| OsJ_36443 | cDNA clone: J013066N15, full insert sequence; Fiber protein Fb19, putative, expressed; Os12g0552500 protein; cDNA clone: J013066N15, full insert sequence |
| OS12T0623900-01 | 5-methyltetrahydropteroyltriglutamate--homocysteine methyltransferase 1; Catalyzes the transfer of a methyl group from 5-methyltetrahydrofolate to homocysteine resulting in methionine formation |
| P0436E04.5 | Probable 3-beta-hydroxysteroid-Delta(8),Delta(7)-isomerase; Catalyzes the conversion of Delta(8)-sterols to their corresponding Delta(7)-isomers |

| | |
|---|---|
| OsJ_01312 | cDNA clone: 006-207-E01, full insert sequence; Os01g0279800 protein; Putative LRR protein; Uncharacterized protein; cDNA clone: 006-207-E01, full insert sequence |
| OS01T0354200-01 | cDNA clone: 002-110-E10, full insert sequence |
| BRI1 | Extra sporogenous cells-like; os01g0718300 protein; cdna clone: j033069j12, full insert sequence; Receptor kinase involved brassinosteroid (BR) signal transduction. Regulates, in response to BR binding, a signaling cascade involved in plant development, promotion of cell elongation and flowering (Probable). Activates BR signaling by targeting and phosphorylating BSK3, a positive regulator of BR signaling. Forms at the plasma membrane a receptor complex with BAK1 which is activated in response to brassinolide. Phosphorylates BAK1. Phosphorylates REM4.1, which reduces REM4.1 binding affi [...] |
| OS01T0935800-01 | Os01g0935800 protein; Nodulation receptor kinase-like; Os01g0935800 protein |
| OS02T0236200-01 | Os02g0236200 protein; Putative Shaggy-related protein kinase dzeta (ASK-dzeta); Belongs to the protein kinase superfamily |
| OS02T0465400-01 | Os02g0465400 protein; Putative sterol delta-7 reductase |
| OS02T0728500-01 | cDNA clone: J033072J21, full insert sequence |
| Smt2-1 | 24-methylenesterol C-methyltransferase 2; Catalyzes the methyl transfer from S-adenosyl-methionine to the methylene group of 24-methylene lophenol to form 24- ethylidene lophenol; Belongs to the class I-like SAM-binding methyltransferase superfamily. Erg6/SMT family |

| | |
|---|---|
| OS03T0703200-01 | Os03g0703200 protein; Expressed protein |
| DRM3 | Probable inactive dna (cytosine-5)-methyltransferase drm3; Involved in de novo DNA methylation. Involved in RNA-directed DNA methylation (RdDM) |
| OsJ_18551 | Putative systemin receptor SR160; Putative uncharacterized protein P0017D10.20; Belongs to the protein kinase superfamily |
| OsJ_19027 | Os05g0491400 protein; Putative uncharacterized protein OSJNBa0088I06.11 |
| GSK4 | Os06g0547900 protein; Shaggy-like kinase etha; cDNA clone: J023139I22, full insert sequence; Probable serine-threonine kinase that may regulate brassinosteroid signaling |
| Smt1-1 | Cycloartenol-C-24-methyltransferase 1; Catalyzes the methyl transfer from S-adenosyl-methionine to the C-24 of cycloartenol to form 24-methylene cycloartenol |
| BZR1 | Putative mature anther-specific protein lat61; cdna clone: 002-115-c06, full insert sequence; Positive brassinosteroid-signaling protein. Mediates downstream brassinosteroid-regulated growth response and feedback inhibition of brassinosteroid (BR) biosynthetic genes May act as transcriptional repressor by binding the brassinosteroid-response element (BREE) (5'-CGTG(T/C)G-3') in the promoter of DLT (AC Q9LWU9), another positive regulator of BR signaling. Acts as transcriptional repressor of LIC, a negative regulator of BR signaling, by binding to the BRRE element of its promoter. BZR1 a [...] |
| OsJ_30372 | cDNA, clone: J080318K21, full insert sequence |

| | |
|---|---|
| OS11T0525200-01 | cDNA clone: 001-014-C05, full insert sequence; Cytochrome P450 51, putative, expressed; Os11g0525200 protein; cDNA clone: 001-014-C05, full insert sequence |
| CYP85A1 | Cytochrome P450 85A1; May convert 6-deoxoteasterone to teasterone, 3-dehydro- 6-deoxoteasterone to 3-dehydroteasterone, and 6-deoxotyphasterol to typhasterol. Catalyzes the C6-oxidation step in brassinosteroids biosynthesis. Involved in the organization and elongation of the leaf and stem cells |
| CYP734A4 | Cytochrome P450 734A4; Cytochrome P450 involved in brassinosteroids (BRs) inactivation and regulation of BRs homeostasis. Is a multifunctional and multisubstrate enzyme that controls the endogenous bioactive BR content both by direct inactivation of castasterone and by decreasing the levels of BR precursors. Catalyzes the oxidation of carbon 22 hydroxylated BR intermediates to produce C26 oxidized metabolites |
| BSL2 | Serine/threonine-protein phosphatase bsl2 homolog; Belongs to the PPP phosphatase family. BSU subfamily |
| OS03T0100800-01 | Os03g0100800 protein; Plasma membrane ATPase 1, putative, expressed |
| OS11T0446500-00 | Phospholipid-transporting atpase; Belongs to the cation transport ATPase (P-type) (TC 3.A.3) family. Type IV subfamily |
| a1 | H-ATPase; Os03g0689300 protein; Plasma membrane ATPase 3, putative, expressed; Plasma membrane H+ ATPase |
| OS04T0353000-01 | annotation not available |
| a2 | Os07g0191200 protein; Plasma membrane H+ ATPase |
| a3 | Os12g0638700 protein; Plasma membrane ATPase 1, putative, expressed; Plasma membrane H+ ATPase |

| | |
|---|---|
| OS03T0120100-01 | LEM3 family/CDC50 family protein, expressed; Os03g0120100 protein |
| OS04T0656100-01 | Plasma membrane ATPase; The plasma membrane ATPase of plants and fungi is a hydrogen ion pump. The proton gradient it generates drives the active transport of nutrients by H(+)-symport. The resulting external acidification and/or internal alkinization may mediate growth responses (By similarity) |
| OS10T0184300-01 | V-type proton ATPase subunit a; Essential component of the vacuolar proton pump (V- ATPase), a multimeric enzyme that catalyzes the translocation of protons across the membranes. Required for assembly and activity of the V-ATPase |
| OS03T0183900-01 | Os03g0183900 protein |
| VATP-P1 | V-type proton ATPase 16 kDa proteolipid subunit; Proton-conducting pore forming subunit of the membrane integral V0 complex of vacuolar ATPase. V-ATPase is responsible for acidifying a variety of intracellular compartments in eukaryotic cells |
| OsJ_05583 | Os02g0175400 protein; Putative vacuolar proton-ATPase |
| OsJ_02567 | Os01g0610100 protein; Uncharacterized protein; Vacuolar H+-exporting ATPase chain c.PPA1-like; cDNA clone: 001-110-C12, full insert sequence; cDNA clone: 006-205-A07, full insert sequence; Belongs to the V-ATPase proteolipid subunit family |
| OS04T0643100-01 | cDNA clone: 002-120-A07, full insert sequence; OSJNBa0033G05.3 protein; OSJNBa0063C18.17 protein; Os04g0643100 protein; cDNA clone: 002-120-A07, full insert sequence; cDNA clone: J013110H12, full insert sequence |

| | |
|---|---|
| OsPTR1 | Os03g0719900 protein; Peptide transporter; Peptide transporter PTR2, putative, expressed; Putative peptide transporter 1; cDNA clone: J013116K11, full insert sequence |
| OsJ_32603 | cDNA clone: J013092G19, full insert sequence; Os10g0579800 protein; POT family protein, expressed; Putative peptide transporter; cDNA clone: J013092G19, full insert sequence |
| OS01T0871600-01 | OSJNBb0008G24.12 protein |
| OsJ_37010 | Os12g0638200 protein; POT family protein, expressed |
| OS05T0431700-01 | Putative uncharacterized protein OJ1301_G07.2; Putative uncharacterized protein OJ1378_A04.7 |
| OsJ_18532 | Putative proton-dependent oligopeptide transporter (POT); cDNA clone: 002-117-H03, full insert sequence; cDNA clone: J023060I20, full insert sequence |
| OsJ_04247 | cDNA clone: J033088C23, full insert sequence; Os01g0872000 protein; Putative peptide transporter; Uncharacterized protein; cDNA clone: J033088C23, full insert sequence |
| OS11T0282800-00 | POT family protein, expressed |
| OsJ_13171 | cDNA clone: 001-020-D02, full insert sequence; Os03g0823500 protein; POT family protein, expressed; Putative peptide transport protein; cDNA clone: 001-020-D02, full insert sequence; cDNA clone: J023109G05, full insert sequence |
| OS06T0581000-01 | Os06g0581000 protein; Putative nitrate transporter NTL1 |
| OS04T0491350-00 | OSJNBa0076N16.18 protein |

| | |
|---|---|
| YSL16 | Probable metal-nicotianamine transporter YSL16; May be involved in the transport of nicotianamine- chelated metals; Belongs to the YSL (TC 2.A.67.2) family |
| B1065G12.20 | Putative peptide transport protein |
| OS07T0603800-01 | Os07g0603800 protein; Putative peptide transporter |
| OS06T0125400-00 | Putative sexual differentiation process protein isp4 |
| OsJ_04516 | cDNA clone: J023114K11, full insert sequence; Os01g0913300 protein; Putative nitrate transporter NRT1-5; Uncharacterized protein; cDNA clone: J023114K11, full insert sequence |
| YSL11 | Probable metal-nicotianamine transporter YSL11; May be involved in the transport of nicotianamine- chelated metals |
| OsJ_32602 | cDNA clone: J013086I05, full insert sequence; Os10g0579600 protein; POT family protein, expressed; Putative peptide transporter; cDNA clone: J013086I05, full insert sequence; cDNA clone: J023018P15, full insert sequence; cDNA clone: J023060H09, full insert sequence |
| YSL6 | Probable metal-nicotianamine transporter YSL6; May be involved in the transport of nicotianamine- chelated metals |
| OS10T0370700-01 | Os10g0370700 protein; POT family protein, expressed; cDNA clone: J013049E13, full insert sequence |
| OS04T0464400-00 | annotation not available |
| OS06T0705600-00 | annotation not available |
| OsJ_04248 | Os01g0872100 protein; Uncharacterized protein |

| | |
|---|---|
| OsJ_10060 | Os03g0235700 protein |
| OsJ_08148 | Os02g0716800 protein; Peptide transporter-like |
| OS11T0283500-01 | Os11g0283500 protein; POT family protein, expressed; cDNA clone: J033023F23, full insert sequence |
| OS01T0556700-00 | annotation not available |
| OS01T0872600-01 | Os01g0872600 protein |
| OsJ_03537 | cDNA clone: 001-128-A10, full insert sequence; Os01g0761400 protein; Putative nitrite transporter; Putative peptide transporter; Uncharacterized protein; cDNA clone: 001-128-A10, full insert sequence |
| YSL14 | Probable metal-nicotianamine transporter ysl14; May be involved in the transport of nicotianamine-chelated metals |
| OsJ_04242 | Os01g0871500 protein; Putative oligopeptide transporter; Uncharacterized protein |
| YSL2 | Metal-nicotianamine transporter YSL2; Involved in the phloem transport of iron and manganese and their translocation into the grain. Transports iron- and manganese-nicotianamine chelates, but not iron-phytosiderophore |
| OS05T0567700-01 | Os05g0567700 protein; Putative uncharacterized protein OJ1781_H11.16 |
| OS11T0135300-00 | Os11g0135300 protein; Major facilitator superfamily antiporter, putative |
| OS04T0453200-01 | OSJNBa0027G07.5 protein; Os04g0453200 protein; Belongs to the major facilitator superfamily. Sugar transporter (TC 2.A.1.1) family |
| MST2 | Sugar transport protein mst2; Mediates active uptake of hexoses by sugar: proton symport. Can transport glucose |

| | |
|---|---|
| OS12T0132500-00 | annotation not available |
| OS09T0394500-01 | cDNA, clone: J065054A13, full insert sequence; Belongs to the major facilitator superfamily. Sugar transporter (TC 2.A.1.1) family |
| VDAC2 | Mitochondrial outer membrane protein porin 2; Forms a channel through the mitochondrial outer membrane that allows diffusion of small hydrophilic molecules. The channel adopts an open conformation at low or zero membrane potential and a closed conformation at potentials above 30-40 mV. The open state has a weak anion selectivity whereas the closed state is cation- selective (By similarity); Belongs to the eukaryotic mitochondrial porin (TC 1.B.8.1) family |
| OS09T0452300-01 | Putative uncharacterized protein; Os09g0452300 protein; Belongs to the major facilitator superfamily. Sugar transporter (TC 2.A.1.1) family |
| OS05T0579000-01 | Os05g0579000 protein; Putative sugar transporter; Belongs to the major facilitator superfamily. Sugar transporter (TC 2.A.1.1) family |
| MST6 | Os07g0559700 protein; Putative monosaccharide transporter 3; Mediates active uptake of hexoses by sugar: proton symport (Probable). Can transport glucose, fructose, mannose, galactose, xylose and ribose |
| OsJ_15010 | Os04g0453400 protein; Belongs to the major facilitator superfamily. Sugar transporter (TC 2.A.1.1) family |
| OsJ_09774 | cDNA clone: J023048F11, full insert sequence; Mannitol transporter, putative, expressed; Os03g0197100 protein; cDNA clone: J023048F11, full insert sequence; Belongs to the major facilitator superfamily. Sugar transporter (TC 2.A.1.1) family |

| | |
|---|---|
| B1364A02.4 | Putative proton myo-inositol transporter; Belongs to the major facilitator superfamily. Sugar transporter (TC 2.A.1.1) family |
| OsJ_09056 | Hexose carrier protein HEX6, putative, expressed; Os03g0101300 protein; cDNA clone: J023037O19, full insert sequence; Belongs to the major facilitator superfamily. Sugar transporter (TC 2.A.1.1) family |
| OS09T0297300-00 | Os09g0297300 protein; Putative monosaccharide transporter 6; Belongs to the major facilitator superfamily. Sugar transporter (TC 2.A.1.1) family |
| OsJ_32870 | Os11g0135900 protein; Transporter, putative, expressed |
| OS09T0268300-02 | Putative hexose carrier protein HEX6; Belongs to the major facilitator superfamily. Sugar transporter (TC 2.A.1.1) family |
| VDAC3 | Mitochondrial outer membrane protein porin 3; Forms a channel through the mitochondrial outer membrane that allows diffusion of small hydrophilic molecules. The channel adopts an open conformation at low or zero membrane potential and a closed conformation at potentials above 30-40 mV. The open state has a weak anion selectivity whereas the closed state is cation- selective (By similarity); Belongs to the eukaryotic mitochondrial porin (TC 1.B.8.1) family |
| OS02T0229400-01 | Os02g0229400 protein; Putative hexose transporter; Tonoplast monosaccharide transporter 2; Belongs to the major facilitator superfamily. Sugar transporter (TC 2.A.1.1) family |
| OS02T0564300-01 | cDNA, clone: J080022K01, full insert sequence; Os02g0564300 protein; Putative uncharacterized protein OJ1712_E04.24; Putative uncharacterized protein P0020C11.13; cDNA, clone: J080022K01, full insert sequence |
| OS04T0533900-01 | Os04g0533900 protein |

| | |
|---|---|
| COPT6 | Copper transporter 6; Involved in the transport of copper |
| MTP7 | Metal tolerance protein 7; Involved in sequestration of excess metal in the cytoplasm into vacuoles to maintain metal homeostasis |
| OS08T0467400-01 | cDNA clone: J023054A11, full insert sequence; Os08g0467400 protein; Putative IAA-alanine resistance protein; cDNA clone: J023054A11, full insert sequence |
| OsJ_16134 | Zinc transporter 3; Zinc transporter that may mediate zinc uptake from the rhizosphere. Seems specific to zinc ions and may not transport other divalent cations |
| NRAMP3 | Metal transporter Nramp3; Probable metal transporter |
| IRT2 | Fe (2+) transport protein 2; Iron transporter that may play a role in the uptake of iron from the rhizosphere across the plasma membrane in the root epidermal layer |
| HMA2 | Cadmium/zinc-transporting atpase hma2; Zinc/cadmium transporter that plays an essential role in promoting translocation of zinc and cadmium from roots to shoots. May control cadmium loading into xylem. In roots, transports zinc and cadmium from the apoplast to the symplast to facilitate translocation via the phloem. In nodes, functions to load zinc and cadmium to the phloem for the preferential distribution to the upper nodes and panicles |
| OS03T0178100-00 | Putative uncharacterized protein; Os03g0178100 protein |

| | |
|---|---|
| P0044F08.14 | cDNA clone: 001-008-D10, full insert sequence; Os01g0125600 protein; P0044F08.14 protein; Putative uncharacterized protein P0037C04.32; cDNA clone: 001-008-D10, full insert sequence |
| MTP6 | Metal tolerance protein 6; Involved in sequestration of excess metal in the cytoplasm into vacuoles to maintain metal homeostasis |
| OS03T0345700-01 | Os03g0345700 protein; Heavy metal-associated domain containing protein, expressed |
| P0532H03.5 | Os06g0690700 protein; Putative cadmium resistance protein |
| NRAMP1 | Metal transporter Nramp1; Probable metal transporter that may participate in the control of iron homeostasis |
| ZIP10 | Zinc transporter 10; Zinc transporter that may be involved in zinc uptake from the rhizosphere |
| STAR2 | UPF0014 membrane protein STAR2; Associates with STAR2 to form a functional transmembrane ABC transporter required for detoxification of aluminum (Al) in roots. Can specifically transport UDP-glucose; Belongs to the UPF0014 family |
| HMA4 | Os02g0196600 protein; Putative copper-transporting P-type ATPase; Copper (Cu) transporter that mediates Cu transport in root vacuoles. Involved in Cu detoxification by sequestrating Cu into root vacuoles and limiting translocation of Cu from the roots to the shoots, and accumulation in grains |
| HMA5 | Putative uncharacterized protein; Copper (Cu) transporter that plays an essential role in promoting translocation of Cu from roots to shoots. Involved in loading Cu to the xylem of the roots and other organs, including panicles |
| NRAMP4 | Metal transporter Nramp4; Probable metal transporter |

| | |
|---|---|
| OsJ_18601 | Os05g0424000 protein; Putative amino acid transporter |
| P0525F01.5 | cDNA clone: 002-115-E07, full insert sequence; Os06g0228600 protein; Putative amino acid transport protein; cDNA clone: 002-115-E07, full insert sequence |
| OS12T0181600-01 | cDNA clone: 001-202-G09, full insert sequence; Amino acid transporter, putative, expressed; Os12g0181600 protein; cDNA clone: 001-202-G09, full insert sequence; cDNA clone: 006-308-D07, full insert sequence |
| OS12T0580400-01 | annotation not available |
| OVP3 | Os02g0802500 protein; Proton translocating pyrophosphatase; Vacuolar proton pyrophosphatase |
| OS11T0195600-01 | Amino acid carrier, putative, expressed; Os11g0195600 protein; Transmembrane amino acid transporter protein; cDNA clone: 001-035-E06, full insert sequence; cDNA clone: J013129H02, full insert sequence |
| OS03T0576900-01 | Amino acid permease family protein, putative, expressed; cDNA clone: 001-047-G03, full insert sequence |
| OS03T0654400-01 | Amino acid permease family protein, expressed; Os03g0654400 protein; Putative amino acid permease; Putative cationic amino acid transporter; cDNA clone: J013063A11, full insert sequence |
| OS12T0156866-00 | Amino acid permease family protein, expressed |
| OS01T0621200-00 | annotation not available |
| OS12T0194900-01 | Amino acid permease I, putative, expressed; Os12g0194900 protein |

| | |
|---|---|
| PUT1 | Polyamine transporter PUT1; Cell membrane polyamine/proton symporter involved in the polyamine uptake in cells. Possesses high affinity for spermidine and lower affinity for spermine and putrescine. Transports paraquat, a polyamine analog, and thus confers sensitivity to this chemical which is used as a herbicide; Belongs to the amino acid-polyamine-organocation (APC) superfamily. Polyamine: cation symporter (PHS) (TC 2.A.3.12) family |
| OS06T0644200-01 | Inorganic diphosphatase, H+-translocating, vacuolar membrane-like |
| OS04T0460300-01 | OSJNBa0072F16.7 protein; Os04g0460300 protein; cDNA clone: 001-208-F06, full insert sequence |
| OsJ_04987 | cDNA clone: J033075N18, full insert sequence; Amino acid transporter-like protein |
| OS02T0727100-01 | Os02g0727100 protein; Putative amino acid transporter; cDNA clone: J023006P21, full insert sequence |
| OsJ_05707 | Os02g0191300 protein; Putative amino acid transporter A1; cDNA clone: J023027D15, full insert sequence |
| OS04T0470700-01 | OSJNBa0089K21.6 protein; Os04g0470700 protein; cDNA clone: J023081F09, full insert sequence |
| OS07T0100800-01 | Probable proline transporter 2; Proline transporter that mediates proline transport across the plasma membrane |
| OS01T0878700-02 | Os01g0878700 protein; Putative amino acid carrier; cDNA clone: 001-203-C05, full insert sequence; cDNA clone: J013002M14, full insert sequence; cDNA clone: J033051N12, full insert sequence |
| OS02T0184200-01 | annotation not available |
| OS02T0670900-02 | Os02g0670900 protein; Putative amino acid transport protein; cDNA clone: 001-038-F02, full insert sequence; cDNA clone: J023111D06, full insert sequence |

| | |
|---|---|
| OsJ_21679 | Os06g0556000 protein; Putative amino acid transporter; cDNA clone: J033107D14, full insert sequence |
| OsJ_36108 | LILLIM08, putative, expressed; Os12g0485600 protein |
| OS11T0155500-01 | Putative uncharacterized protein; Os11g0155500 protein |
| P0451D05.6 | Os01g0593700 protein; Sulfate transporter 2-like |
| OS01T0719300-01 | Os01g0719300 protein; Putative plasma membrane sulphate transporter |
| PHT4;3 | Probable anion transporter 3, chloroplastic; Probable anion transporter |
| PT4 | Probable inorganic phosphate transporter 1-4; High-affinity transporter for external inorganic phosphate |
| OsJ_07345 | Phosphate transporter; Sodium-phosphate symporter which plays a fundamental housekeeping role in phosphate transport |
| NRT2.3 | Membrane transporter-like; os01g0704100 protein; Involved in nitrate transport, but does not seem to be able to mediate transport by its own. Acts as a dual component transporter with NAR2.1. Imports nitrate with high affinity when expressed with NAR2.1 in a heterologous system (Xenopus oocytes). Plays a key role in long-distance nitrate transport from root to shoot particularly at low external nitrate supply |
| NAR2.1 | Component of high affinity nitrate transporter-like protein; os02g0595900 protein; Acts as a dual component transporter with NTR2.1, NRT2.2 and NRT2.3. Required for high-affinity nitrate transport. Involved in the regulation of NRT2.1, NRT2.2 and NRT2.3 expression, and in both, HATS (high-affinity transport system) and LATS (low-affinity transport system) |

| | |
|---|---|
| | activities in plant roots. Imports nitrate with high affinity when expressed with NTR2.1, NTR2.2 or NTR2.3 in a heterologous system (Xenopus oocytes) |
| OS01T0695700-00 | Os01g0695700 protein |
| mdr17 | MDR-like ABC transporter; MDR-like p-glycoprotein-like; Os01g0723800 protein; Uncharacterized protein |
| OS01T0904200-01 | ABC1-like; Os01g0904200 protein; cDNA clone: J013153C13, full insert sequence |
| OS01T0911300-01 | Os01g0911300 protein |
| OS02T0288400-00 | Os02g0288400 protein; MRP-like ABC transporter |
| OS02T0323000-01 | Os02g0323000 protein |
| OsJ_07239 | Os02g0575500 protein; Putative ABC transporter |
| mdr11 | Putative MDR-like ABC transporter |
| OS02T0826500-01 | Putative ABC transporter |
| OS03T0157400-01 | ABC transporter family protein, putative, expressed; Os03g0157400 protein |
| OS04T0209200-01 | Os04g0209200 protein |
| OS04T0209300-01 | OSJNBb0022P19.1 protein; Os04g0209300 protein; cDNA clone: J013026A05, full insert sequence |
| OsJ_19973 | Os06g0128300 protein; Putative mitochondrial half-ABC transporter |
| OS06T0503100-01 | Os06g0503100 protein; Putative ABCG4 |

| | |
|---|---|
| P0462E11.2 | Putative ABC transporter AbcG1 |
| ABCG44 | Abc transporter g family member 44; May be a general defense protein |
| OsJ_28532 | ABC1 family protein-like; Os09g0250700 protein |
| OS09T0250800-00 | Os09g0250800 protein |
| OsJ_30438 | Os09g0572400 protein; Putative iron inhibited ABC transporter 2 |
| ABCG51 | Abc transporter g family member 51; May be a general defense protein |
| OS10T0432200-01 | Retrotransposon protein, putative, unclassified, expressed |
| OsJ_32262 | Expressed protein; Os10g0533100 protein; Putative uncharacterized protein OSJNBa0053C23.22 |
| OS06T0158900-00 | Os06g0158900 protein; Putative multidrug-resistance associated protein |
| OS11T0177400-01 | Os11g0177400 protein |
| OS11T0546000-01 | Os11g0546000 protein; ATP-binding cassette sub-family E member 1, putative, expressed; Os11g0546000 protein |
| ABCG45 | Abc transporter g family member 45; May be a general defense protein |
| OS12T0239900-01 | Os12g0239900 protein |
| OS12T0409700-00 | annotation not available |
| P0031D11.2 | Putative uncharacterized protein P0031D11.2 |
| PIN5A | Probable auxin efflux carrier component 5a; May act as a component of the auxin efflux carrier |

| | |
|---|---|
| acsF | Magnesium-protoporphyrin IX monomethyl ester [oxidative] cyclase; Catalyzes the formation of the isocyclic ring in chlorophyll biosynthesis. Mediates the cyclase reaction, which results in the formation of divinylprotochlorophyllide (Pchlide) characteristic of all chlorophylls from magnesium-protoporphyrin IX 13-monomethyl ester (MgPMME) |
| RH3 | Os01g0508100 protein; cDNA, clone: J065211P06, full insert sequence; Binds to and represses NPR1/NH1-mediated transcriptional activation of LG2 in vitro |
| RnrS2 | Os06g0127900 protein; Putative ribonucleotide reductase R2; Ribonucleotide diphosphate reductase small subunit 2; cDNA clone: J033022H19, full insert sequence |
| OsJ_01952 | cDNA clone: 006-310-A06, full insert sequence; Farnesylated protein 2-like; Os01g0507700 protein; Uncharacterized protein; cDNA clone: 006-310-A06, full insert sequence |
| OsJ_04662 | Copper chaperone (CCH)-related protein-like; Os01g0933200 protein; Uncharacterized protein |
| P0020E09.5 | Heavy-metal-associated domain-containing protein-like; Os01g0976300 protein; cDNA clone: 002-181-G11, full insert sequence |
| OS02T0510600-01 | Os02g0510600 protein; Putative farnesylated protein |
| OsJ_09136 | cDNA, clone: J100050C22, full insert sequence |
| OsJ_11078 | Heavy-metal-associated domain-containing protein, putative, expressed; Os03g0383900 protein; cDNA clone: J023033I19, full insert sequence |

| | |
|---|---|
| OSJNBa0032E21.09 | Heavy metal-associated domain containing protein, expressed; Putative uncharacterized protein OJ1112_G08.2; Putative uncharacterized protein OSJNBa0032E21.09; cDNA clone: 006-310-A12, full insert sequence; cDNA clone: 006-311-G12, full insert sequence |
| OS04T0667600-01 | Os04g0667600 protein |
| B1032F05.3 | Os07g0298900 protein; Putative heavy-metal-associated domain-containing protein |
| OS07T0671400-01 | Os07g0671400 protein |
| OsJ_26405 | Os08g0205400 protein; Putative copper chaperone; cDNA, clone: J100082I23, full insert sequence |
| OsJ_27250 | cDNA clone: 002-141-A09, full insert sequence; Os08g0405700 protein; Putative uncharacterized protein P0685B10.33; cDNA clone: 002-141-A09, full insert sequence |
| OS09T0364800-01 | Os09g0364800 protein |
| OS09T0408550-00 | Os09g0408500 protein |
| OS12T0421000-01 | Heavy metal-associated domain containing protein, expressed; Os12g0421000 protein |
| OS03T0571700-01 | Os03g0571700 protein |
| OsJ_11504 | MATE efflux family protein, expressed; Os03g0572900 protein; Putative MATE efflux family protein; cDNA clone: J033087G07, full insert sequence; Belongs to the multi antimicrobial extrusion (MATE) (TC 2.A.66.1) family |
| OS05T0554000-01 | cDNA clone: 001-123-D07, full insert sequence |

| | |
|---|---|
| OS06T0495100-00 | annotation not available |
| P0430F03.2 | Os07g0502200 protein; Putative MATE efflux protein family protein; Belongs to the multi antimicrobial extrusion (MATE) (TC 2.A.66.1) family |
| OS07T0516600-01 | Os07g0516600 protein; cDNA clone: J023025C14, full insert sequence; Belongs to the multi antimicrobial extrusion (MATE) (TC 2.A.66.1) family |
| OS10T0206800-01 | MATE efflux family protein, putative, expressed; Os10g0206800 protein; Putative membrane protein; cDNA clone: J033052E02, full insert sequence; Belongs to the multi antimicrobial extrusion (MATE) (TC 2.A.66.1) family |
| OS10T0344900-01 | MATE efflux family protein, expressed; Os10g0344900 protein; Putative integral membrane protein; cDNA clone: J033012J16, full insert sequence; cDNA clone: J033119H22, full insert sequence; Belongs to the multi antimicrobial extrusion (MATE) (TC 2.A.66.1) family |
| OS10T0345100-01 | Protein detoxification; Os10g0345100 protein; Belongs to the multi antimicrobial extrusion (MATE) (TC 2.A.66.1) family |
| OS11T0129200-00 | Os11g0129100 protein |
| OS12T0125500-01 | annotation not available |
| OsJ_35077 | MATE efflux family protein, expressed; Os12g0126000 protein; Belongs to the multi antimicrobial extrusion (MATE) (TC 2.A.66.1) family |

| | |
|---|---|
| OsJ_36878 | Os12g0615700 protein; TRANSPARENT TESTA 12 protein, putative, expressed; cDNA clone: 001-016-G09, full insert sequence; Belongs to the multi antimicrobial extrusion (MATE) (TC 2.A.66.1) family |
| OS07T0606900-01 | Putative uncharacterized protein OSJNBa0072I06.16; Putative uncharacterized protein P0493C06.31 |
| OS05T0138300-03 | Putative uncharacterized protein OSJNBa0069I13.5 |

**Table S3** DAVID enrichment analysis results for each sub-module of the seed development network of rice.

| Module | GO-BP Term | P Value |
|---|---|---|
| 1 | GO: 0055076~transition metal ion homeostasis | 1.95E-08 |
| | GO: 0055065~metal ion homeostasis | 2.24E-07 |
| | GO: 0055080~cation homeostasis | 1.11E-06 |
| | GO: 0098771~inorganic ion homeostasis | 1.33E-06 |
| | GO: 0050801~ion homeostasis | 2.16E-06 |
| | GO: 0048878~chemical homeostasis | 9.52E-06 |
| | GO: 0042592~homeostatic process | 2.75E-05 |
| | GO: 0055085~transmembrane transport | 7.70E-05 |
| 2 | GO: 0006096~glycolytic process | 1.86E-10 |
| | GO: 0006094~gluconeogenesis | 2.71E-06 |
| | GO: 0019521~D-gluconate metabolic process | 3.31E-06 |

| | | |
|---|---|---|
| | GO: 0006098~pentose-phosphate shunt | 2.08E-04 |
| | GO: 0006006~glucose metabolic process | 4.11E-04 |
| | GO: 0046177~D-gluconate catabolic process | 0.003247 |
| | GO: 0009051~pentose-phosphate shunt, oxidative branch | 0.007562 |
| 3 | GO: 0042026~protein refolding | 4.95E-14 |
| | GO: 0051085~chaperone mediated protein folding requiring cofactor | 2.84E-06 |
| | GO: 0034620~cellular response to unfolded protein | 6.72E-05 |
| | GO: 0006457~protein folding | 0.004277 |
| 4 | GO: 0009098~leucine biosynthetic process | 5.12E-09 |
| | GO: 0009082~branched-chain amino acid biosynthetic process | 7.31E-08 |
| | GO: 0006526~arginine biosynthetic process | 4.67E-05 |
| | GO: 0006099~tricarboxylic acid cycle | 8.53E-04 |
| | GO: 0000053~argininosuccinate metabolic process | 0.001402 |
| | GO: 0000050~urea cycle | 0.002102 |
| 5 | GO: 0030154~cell differentiation | 0.001517 |
| | GO: 0045944~positive regulation of transcription from RNA polymerase II promoter | 0.002467 |
| | GO: 0070734~histone H3-K27 methylation | 0.003183 |
| | GO: 0009960~endosperm development | 0.009523 |
| | GO: 0000122~negative regulation of transcription from RNA polymerase II promoter | 0.012048 |
| | GO: 0006342~chromatin silencing | 0.014568 |
| 6 | GO: 1902600~hydrogen ion transmembrane transport | 5.58E-15 |

| | | |
|---|---|---|
| | GO: 0051453~regulation of intracellular pH | 3.89E-12 |
| | GO: 0015986~ATP synthesis coupled proton transport | 0.02558 |
| 7 | GO: 0009086~methionine biosynthetic process | 4.96E-04 |
| | GO: 0006431~methionyl-tRNA aminoacylation | 0.004582 |
| | GO: 0006749~glutathione metabolic process | 0.005105 |
| | GO: 0019243~methylglyoxal catabolic process to D-lactate via S-lacteal-glutathione | 0.005724 |
| | GO: 0006436~tryptophanyl-tRNA aminoacylation | 0.005724 |
| 8 | GO: 0010499~proteasomal ubiquitin-independent protein catabolic process | 0.007309 |
| | GO: 0043161~proteasome-mediated ubiquitin-dependent protein catabolic process | 0.025552 |
| 9 | GO: 0006412~translation | 9.26E-09 |
| | GO: 0002183~cytoplasmic translational initiation | 0.010661 |
| | GO: 0000398~mRNA splicing, via spliceosome | 0.02266 |
| | GO: 0006414~translational elongation | 0.030172 |
| | GO: 0000470~maturation of LSU-rRNA | 0.037578 |
| 10 | GO: 0009873~ethylene-activated signaling pathway | 1.75E-05 |
| | GO: 1901001~negative regulation of response to salt stress | 0.005597 |
| | GO: 0010104~regulation of ethylene-activated signaling pathway | 0.006991 |
| | GO: 0006378~mRNA polyadenylation | 0.017395 |
| 11 | GO: 0010097~specification of stamen identity | 4.46E-06 |
| | GO: 0030154~cell differentiation | 3.21E-05 |
| | GO: 0006357~regulation of transcription from RNA polymerase II promoter | 0.001049 |

| | | |
|---|---|---|
| | GO: 0045944~positive regulation of transcription from RNA polymerase II promoter | 0.003 |
| 12 | GO: 0009738~abscisic acid-activated signaling pathway | 1.28E-12 |
| | GO: 0009688~abscisic acid biosynthetic process | 2.31E-07 |
| | GO: 0009414~response to water deprivation | 3.84E-05 |
| | GO: 0006817~phosphate ion transport | 2.08E-04 |
| | GO: 0045893~positive regulation of transcription, DNA-templated | 3.80E-04 |
| 13 | GO: 0080113~regulation of seed growth | 0.001911 |
| | GO: 0080050~regulation of seed development | 0.004582 |
| | GO: 0048638~regulation of developmental growth | 0.00801 |
| | GO: 0050794~regulation of cellular process | 0.009702 |
| | GO: 0050789~regulation of biological process | 0.015047 |
| | GO: 0048589~developmental growth | 0.016734 |
| 14 | GO: 0016126~sterol biosynthetic process | 2.17E-14 |
| | GO: 0009742~brassinosteroid mediated signaling pathway | 1.92E-07 |
| | GO: 0016125~sterol metabolic process | 3.03E-05 |
| | GO: 0032259~methylation | 0.001446 |
| | GO: 0040008~regulation of growth | 0.001533 |

**Table S4** The most relevant enriched GO-BP term and statistics for each discovered module in the extracted seed development subnetwork of rice, including the total number of proteins within the module, lists of seed proteins, predicted proteins, and their respective counts.

| **Module** | **Enriched GO-BP Term** | **Total number of proteins** | **Number of seed proteins** | **Seed proteins** | **Number of predicted proteins** | **Predicted proteins** |
|---|---|---|---|---|---|---|
| 1 | Transition metal ion homeostasis (GO: 0055076) | 17 | 3 | SODCP, SODCC2, SODCC1 | 14 | HMA4, OsJ_26405, HMA5, OsJ09136, YSL6, OS03T0178100-00, OS08T0467400-01, NRAMP4, HMA2, NRAMP3, OS07T0671400-01, ZIP10, P0532H03.5, NRAMP1 |
| 2 | Glycolytic process (GO: 0006096) | 16 | 15 | OsJ_22300, OsJ12328, OS08T0478800-01, OS03T0266200-01, TPI, OS01T0118000-01, FBA3, AGPL2, OsJ15699, G6PGH1, G6PGH2, UGP, SSIIIA, | 1 | OsJ_09272 |

| | | | | | | |
|---|---|---|---|---|---|---|
| | | | | OS02T0794700-01, SS1 | | |
| 3 | Protein refolding (GO: 0042026) | 9 | 4 | OsJ_31804, OS06T0114000-02, OsJ_04024, OsJ_10329 | 5 | OsJ_09152, OS04T0348300-01, OS10T0566700-00, OsJ_30438, OsJ_04987 |
| 4 | Leucine biosynthetic process (GO: 0009098) | 9 | 7 | SDH1, OS03T0655700-01, OsIDHa, OS11T0303050-00, OsJ_35666, OsJ_35174, OsJ_02861 | 2 | OsJ_32925, OS03T0720300-01 |
| 5 | Endosperm development (GO: 0009960) | 12 | 3 | OsJ_25971, OsJ_25973, MET1B | 9 | OS12T0279100-01, Ehd2, MADS56, HD1, OS10T0506800-01, OsJ_35872, DRM3, MADS15, OS02T0610500-01 |

| | | | | | | |
|---|---|---|---|---|---|---|
| 6 | Hydrogen ion transmembrane transport (GO: 1902600) | 20 | 6 | OsJ_16709, V-ATPase B, OS01T0685800-01, RINO1, OS03T0363500-02, OLE18 | 14 | OS10T0320400-01, OS04T0643100-01, VATP-P1, OsJ05583, OsJ02567, OS10T0184300-01, OS03T0278900-01, OS03T0183900-01, a2, OS03T0100800-01, OS04T0656100-01, a3, OsJ_36878, a1 |
| 7 | Methionine biosynthetic process (GO: 0009086) | 16 | 9 | DHAR1, OsJ_29684, APX1, GLYI-11, OS12T0540900-01, OS05T0230900-01, OsJ_32231, OsJ_07966, B1045D11.6 | 7 | OsJ_09866, OsJ_09517, mdr17, OS12T0623900-01, OS11T0195600-01, OS03T0313100-01, mdr11 |
| 8 | Proteasomal ubiquitin-independent protein catabolic process (GO: 0010499) | 8 | 2 | OsJ_07639, PBF1 | 6 | OsRPT2b, OsJ_21850, OsJ_12657, OsJ_27077, OsJ_13099, HSFA6B |

| | | | | | | |
|---|---|---|---|---|---|---|
| 9 | Translation (GO: 0006412) | 30 | 6 | RACK1A, OsJ_06933, OS08T0308100-01, OsJ_07364, OsJ_05367, PDIL1-1 | 24 | RPL3B, OsJ_09121, OsJ_10495, OsJ_23275, OS02T0103700-01, OS11T0153800-01, OS11T0546000-01, OS03G0109500, OsJ_30578, OsJ_10726, OS03T0725000-01, OsJ_12590, OsJ_11750, OsJ_11922, OS03T0213100-01, OsJ_33580, OsJ_19788, OS02T0564300-01, OS02T0826500-01, OsJ_23479, OsJ_19973, YAB2, RS2, qSH1 |
| 10 | Ethylene-activated signaling pathway (GO: 0055076) | 14 | 3 | MPK1, GF14D, GF14A | 11 | BSL2, OS06T0677700-00, OsJ_13407, OS08T0508700-01, EIL1A, OsJ_12021, OS03T0172000-01, OsEIL2, OsJ_13442, MYC2, OS04T0456900-00 |
| 11 | Cell differentiation (GO: 0030154) | 15 | 4 | CYCB2-2, OS04T0486500-01, OsJ_09943, RPA1B | 11 | OsJ_04312, OS12T0158800-01, Act, RnrS2, OsJ_18684, MADS16, MADS4, OS01T0229000-00, |

| | | | | | | |
|---|---|---|---|---|---|---|
| | | | | | | OS12T0605500-01, OS12T0238000-00, OS01T0695900-01 |
| 12 | Abscisic acid-activated signaling pathway (GO: 0009738) | 22 | 0 | - | 22 | ABI5, OsJ_13064, BZIP12, SAPK10, ZEP, SAPK3, SAPK1, BZIP23, OS09T0456200-01, SAPK7, OS03T0790700-01, CYP707A5, OS03T0790900-01, NCED1, VP1, OsJ_07239, OsJ_07345, OS07T0164900-01, DAO, LFL1, OsJ_28532, PT4 |
| 13 | Regulation of seed growth (GO: 0080113) | 6 | 1 | APG | 5 | OS03T0639300-01, GAI, OS07T0143200-00, OsJ_02811, ILI5 |
| 14 | Brassinosteroid-mediated signaling pathway (GO: 0016126) | 19 | 2 | SERK2, ILI6 | 17 | OS02T0465400-01, Smt2-1, OS01T0354200-01, P0436E04.5, OsJ_30372, BRI1, Smt1-1, DGAT1-2, BZR1, OS11T0525200-01, CYP85A1, OsJ_18551, OS03T0810900-01, |

| | | | | | | |
|---|---|---|---|---|---|---|
| | | | | | | OS03T0703200-01, GSK4, OS02T0236200-01, CYP734A4 |

**Table S5** Details of intra-modular hubs and prominent seed and predicted proteins with literature support for their implications in the rice seed development of each sub-module.

| **Sub-module** | **Associated pathway** | **Protein name** | **Protein Category** | **Implications of rice seed development and literature support** |
|---|---|---|---|---|
| 1 | Transition metal ion homeostasis | HMA4 | Predicted protein Module hub | Responsible for the transportation of copper and associated with the quantitative trait loci that influence the accumulation of copper in rice grains. [1]. A potential solution for addressing copper micronutrient deficiency. |
| | | Osj_26405 | Predicted protein Module hub | Further experimental studies are warranted to explore its role in rice grain development. |
| | | HMA2 | Predicted protein | Participates in transition metal ion processes [2]. Mutant lines showing decreased expression of |

| | | | | |
|---|---|---|---|---|
| | | | | HMA2 experienced reductions in biomass and grain yield. |
| 2 | Glycolytic process | Phosphoglycerate kinase (PGK) | Seed<br>Module hub | Increases the concentration of pyruvate, resulting in heightened levels of carotenoids in rice seeds.<br>[3] |
| | | Enolase, Triosephosphate, SSIIIa, and UGP isomerase | Seed | Grain starch synthesis<br>[4–6] |
| 3 | Protein Refolding | 60 kDa chaperonin (OsJ_31804) | Seed<br>Module hub | Stabilize and refold proteins in the grain-filling stage.<br>[7] |
| 4 | Leucine biosynthetic process | OS03T0655700-01 (3-isopropyl malate dehydrogenase) | Seed<br>Module hub | Leucine biosynthesis<br>[8] |
| | | SDH1 | Seed<br>Module hub | Critical role as an enzyme participating in both the electron transport chain (ETC) and the tricarboxylic acid (TCA) cycle to generate ATP for cellular energy production [9].<br>No direct evidence of involvement in leucine biosynthesis. However, this protein may contribute to the ATP pool demand required for the process. |

| | | | | |
|---|---|---|---|---|
| 5 | Endosperm development | OsFIE2 (OsJ_25971) and METB1 | Seed | Crucial for the development of endosperm in rice grains.<br>[10,11]. |
| 6 | Hydrogen ion transmembrane transport | OsJ_16709 (Inorganic disphosphatase) | Seed<br>Module hub | Plays a critical role in the metabolic process of phosphate-containing compounds by hydrolyzing diphosphate in the presence of water.<br>Although its involvement in H+ ion transmembrane transport is uncertain, the hydrolysis reaction of diphosphate releases H+ ions that can participate in the electrochemical gradient driving ion transport across biological membranes. |
| | | OS03T0100800-01, V-ATPase B, OS03T0183900-01, and OS04T0656100-01 (Plasma membrane ATPase), a1, a2, and a3 | Seed | Hydrogen ion transmembrane transport in rice grain development.<br>[12] |

| | | | | |
|---|---|---|---|---|
| 7 | Methionine biosynthetic process | DHAR1 | Seed<br>Module hub | Significantly enhance grain yield and biomass. Also, it improves the AsA pool (Ascorbic Acid) and redox homeostasis [13]. |
| | | OS12T0623900-01 (Lactoylglutathione lyase), OS12T0540900-01 (Tryptophanyl-tRNA synthetase) | Seed | Exhibit a preferential expression pattern and are linked to the metabolism of amino acids associated with seed development [14]. |
| | | APx1 | Seed | Plays a crucial role in seed development, including fertilization, and acts as a regulator of seed development [15]. |
| 8 | Proteasomal ubiquitin-independent protein catabolic process | OsRPT2b | Predicted protein<br>Module hub | Interacts with SG6, a grain size-determining protein, suggesting that SG6 may be involved in the Ubiquitin-Proteasome System (UPS) pathway to regulate cell division [16]. |
| 9 | Translation | OsJ_09121(40S ribosomal protein S17), OsJ_23275 - 40S ribosomal protein S18, OS02T0103700-01, | Seed | Protein Translation<br>[17,18]. |

| | | | | |
|---|---|---|---|---|
| | | OS03T0725000-01 (Mitochondrial 60S ribosomal protein L6), (60S ribosomal protein L22-2), OsJ_12590 (50S ribosomal protein L6), OsJ_10495 (40S ribosomal protein S7), and OsJ_06933 (Elongation factor 2) | | |
| | | RPL3B | Predicted protein | Play a role in plant architecture in rice through its regulation of ribosome biogenesis. [19]. However, further research is needed to fully understand the mechanisms contributing to rice grain development. |
| 10 | Ethylene-activated signaling pathway | MPK1 | Seed<br>Module hub | Knock-out mutants of MPK1 display defective embryo development [20]. However, it is currently unclear whether MPK1 has a role in ethylene-signaling pathways. |
| | | OsEIL2 and EIA1 | Seed | Ethylene-signaling [21] |

| | | | | |
|---|---|---|---|---|
| 11 | Cell differentiation | CYCB2-2 | Seed<br>Module Hub | Involvement in regulating cell division and differentiation [22]. |
| 12 | Abscisic acid-activated signaling pathway | ABI5,<br>OsJ_13064 | Predicted proteins<br>Module hubs | ABA-mediated transcription [23]. |
| 13 | Regulation of seed growth | APG | Seed<br>Module hub | Regulates rice grain length and weight by controlling cell elongation in lemma/palea through heterodimerization [24]. |
| 14 | Brassinosteroid-mediated signaling pathway | BZR1, BRI1, CYP734A4, SERK2, and GSK4 | Seed | Increases seed grain width, length, and thickness [25–28] |
| | | Sterol delta-7-reductase (OS02T0465400-01) | Predicted protein | Involves in the biosynthesis of brassinosteroids [29]. Although its differential expression is associated with rice seed development [30], there is currently no available literature that |

| | | | | |
|---|---|---|---|---|
| | | | | characterizes the enzyme's specific contribution to this process. |

**Table S6** Overview of inter-module hubs in rice seed development, featuring protein, common name, protein family, seed/predicted protein status, connecting pathways, function, and notes.

| Protein | Common Name | Protein Family | Status | Pathways Connected | Function | Notes |
|---|---|---|---|---|---|---|
| SODCP | Superoxide dismutase copper/zinc | Copper/zinc superoxide dismutases | Seed | Transition metal ion homeostasis (sub-module 1), glycolytic process (sub-module 2), hydrogen ion transmembrane transport (sub-module 6), methionine biosynthesis (sub-module 7) | Combat ROS during grain filling, detoxifies superoxide radicals [31]. | shows a preferential accumulation in the rice embryo [32]. further investigation needed in methionine biosynthesis. |

| | | | | | | |
|---|---|---|---|---|---|---|
| SDH1 | Succinate dehydrogenase 1 | FAD-dependent oxidoreductase 2 | Seed | Hydrogen ion transmembrane transport (sub-module 6), protein refolding (sub-module 3), glycolytic process (sub-module 2), translation (sub-module 9) | Involve in tricarboxylic acid cycle. | Hydrogen ion (H+) transmembrane transport in rice is critical for regulating TCA cycle enzymes [33]. Post-transcriptional regulation, like translation, is key for TCA cycle enzyme control [34]. ATP-dependent mechanisms are essential for protein biosynthesis and refolding, while proper glycolysis and TCA cycle function are vital for rice grain development [35]. These findings highlight the |

| | | | | | | |
|---|---|---|---|---|---|---|
| | | | | | | importance of interplay between linked pathways, necessitating further experimental validation. |
| OS10T0320400-01 | Gamma subunit of F1 complex (ATP synthase) | ATPase gamma chain family. | Predicted protein | Translation (sub-module 9), endosperm development (sub-module 5), hydrogen ion transport (sub-module 6), transition metal ion homeostasis (sub-module 1) | Plays a crucial role in the light-dependent reactions of photosynthesis [36]. It utilizes the proton motive force across the thylakoid membrane to synthesize ATP from ADP and | Given the energy dependence of the connecting pathways, the protein OS10T0320400-01, involved in ATP biogenesis, likely plays a crucial role in mediating their functions. However, its role as an intermodular hub lacks experimental support. |

| | | | | | | |
|---|---|---|---|---|---|---|
| | | | | | inorganic phosphate. | |
| OS01T0685800-01 | Beta subunit of ATP synthase F1 complex | ATP synthase | Seed | Hydrogen ion transmembrane transport (sub-module 6), leucine biosynthetic process (sub-module 4), translation (sub-module 9) | Involve in ATP biosynthesis. | A study manipulating F1-ATPase β subunit (AtpB) expression in rice seeds found it crucial for grain filling and sensitive to high-temperature stress [37]. However, it is not clear how this protein mediates these pathways interlinking. |
| OS03T0278900-01 | ATP synthase B chain | ATP synthase, F0 complex | Predicted protein | Glycolytic process (sub-module 2), translation (sub-module 9), hydrogen ion transmembrane transport (sub-module 6) | Facilitates proton translocation during ATP biosynthesis [38]. | Further studies are needed to fully characterize the involvement of the protein in connecting these pathways during rice grain development. |

| | | | | | | |
|---|---|---|---|---|---|---|
| OsJ_07364 | Translation elongation factor (EFTu) | Translation elongation factor | Seed | Leucine biosynthetic process (sub-module 4), hydrogen ion transmembrane transport (sub-module 6), translation (sub-module 9) | Involve in the elongation step of translation [39]. Also, it has a role in rice grain filling [40]. | Currently no existing evidence for its role in mediating crosstalk among the connecting pathways. |

**References**


1. Huang XY, Deng F, Yamaji N, Pinson SRM, Fujii-Kashino M, Danku J, et al. A heavy metal P-type ATPase OsHMA4 prevents copper accumulation in rice grain. Nat Commun. 2016;7.
2. Yamaji N, Xia J, Mitani-Ueno N, Yokosho K, Ma JF. Preferential delivery of zinc to developing tissues in rice is mediated by P-type heavy metal ATPase OsHMA2. Plant Physiol. 2013;162.
3. Gayen D, Ghosh S, Paul S, Sarkar SN, Datta SK, Datta K. Metabolic regulation of carotenoid-enriched golden rice line. Front Plant Sci. 2016;7.
4. Zhang H, Chen J, Shan S, Cao F, Chen G, Zou Y, et al. Proteomic profiling reveals differentially expressed proteins associated with amylose accumulation during rice grain filling. BMC Genomics. 2020;21.
5. Wei X, Jiao G, Lin H, Sheng Z, Shao G, Xie L, et al. GRAIN INCOMPLETE FILLING 2 regulates grain filling and starch synthesis during rice caryopsis development. J Integr Plant Biol. 2017;59:134–53.
6. Tovy A, Tov RS, Gaentzsch R, Helm M, Ankri S. A new nuclear function of the Entamoeba histolytica glycolytic enzyme enolase: The metabolic regulation of cytosine-5 methyltransferase 2 (Dnmt2) activity. PLoS Pathog. 2010;6.
7. Timabud T, Yin X, Pongdontri P, Komatsu S. Gel-free/label-free proteomic analysis of developing rice grains under heat stress. J Proteomics. 2016;133:1–19.
8. Sikdar MSI, Kim JS. Isolation of a gene encoding 3-isopropylmalate dehydrogenase from rice. Russian Journal of Plant Physiology. 2011;58:190–6.
9. Li C, Liu CQ, Zhang HS, Chen CP, Yang XR, Chen LF, et al. Lps1, encoding iron-sulfur subunit sdh2-1 of succinate dehydrogenase, affects leaf senescence and grain yield in rice. Int J Mol Sci. 2021;22:1–20.

10. Nallamilli BRR, Zhang J, Mujahid H, Malone BM, Bridges SM, Peng Z. Polycomb Group Gene OsFIE2 Regulates Rice (Oryza sativa) Seed Development and Grain Filling via a Mechanism Distinct from Arabidopsis. PLoS Genet. 2013;9.
11. Wang L, Yuan J, Ma Y, Jiao W, Ye W, Yang DL, et al. Rice Interploidy Crosses Disrupt Epigenetic Regulation, Gene Expression, and Seed Development. Mol Plant. 2018;11:300–14.
12. Xue LJ, Zhang JJ, Xue HW. Genome-wide analysis of the complex transcriptional networks of rice developing seeds. PLoS One. 2012;7.
13. Kim YS, Kim IS, Bae MJ, Choe YH, Kim YH, Park HM, et al. Homologous expression of cytosolic dehydroascorbate reductase increases grain yield and biomass under paddy field conditions in transgenic rice (Oryza sativa L. japonica). Planta. 2013;237:1613–25.
14. Xu HH, Liu SJ, Song SH, Wang RX, Wang WQ, Song SQ. Proteomics analysis reveals distinct involvement of embryo and endosperm proteins during seed germination in dormant and non-dormant rice seeds. Plant Physiology and Biochemistry. 2016;103:219–42.
15. Kim YJ, Kim S-I, Kesavan M, Kwak JS, Song JT, Seo HS. Ascorbate Peroxidase OsAPx1 is Involved in Seed Development in Rice. Plant Breed Biotechnol. 2015;3:11–20.
16. Zhou SR, Xue HW. The rice PLATZ protein SHORT GRAIN6 determines grain size by regulating spikelet hull cell division. J Integr Plant Biol. 2020;62:847–64.
17. Doroshenk KA, Crofts AJ, Morris RT, Wyrick JJ, Okita TW. Proteomic analysis of cytoskeleton-associated RNA binding proteins in developing rice seed. J Proteome Res. 2009;8:4641–53.
18. Zhang YX, Xu HH, Liu SJ, Li N, Wang WQ, Møller IM, et al. Proteomic analysis reveals different involvement of embryo and endosperm proteins during aging of Yliangyou 2 hybrid rice seeds. Front Plant Sci. 2016;7.
19. Zheng M, Wang Y, Liu X, Sun J, Wang Y, Xu Y, et al. The RICE MINUTE-LIKE1 (RML1) gene, encoding a ribosomal large subunit protein L3B, regulates leaf morphology and plant architecture in rice. J Exp Bot. 2016;67:3457–69.
20. Minkenberg B, Xie K, Yang Y. Discovery of rice essential genes by characterizing a CRISPR-edited mutation of closely related rice MAP kinase genes. Plant Journal. 2017;89:636–48.
21. Zhao H, Yin CC, Ma B, Chen SY, Zhang JS. Ethylene signaling in rice and Arabidopsis: New regulators and mechanisms. J Integr Plant Biol. Blackwell Publishing Ltd; 2021. p. 102–25.
22. Yang BJ, Wendrich JR, De Rybel B, Weijers D, Xue HW. Rice microtubule-associated protein IQ67-DOMAIN14 regulates grain shape by modulating microtubule cytoskeleton dynamics. Plant Biotechnol J. 2020;18:1141–52.
23. Ali F, Qanmber G, Li F, Wang Z. Updated role of ABA in seed maturation, dormancy, and germination. J Adv Res. Elsevier B.V.; 2022. p. 199–214.
24. Heang D, Sassa H. An atypical bHLH protein encoded by POSITIVE REGULATOR OF GRAIN LENGTH 2 is involved in controlling grain length and weight of rice through interaction with a typical bHLH protein APG. Breed Sci. 2012;62:133–41.
25. LI ZW, Xiong J, QI XH, WANG JY, CHEN HF, ZHANG ZX, et al. Differential Expression and Function Analysis of Proteins in Flag Leaves of Rice During Grain Filling. Acta Agronomica Sinica. 2009;35:132–9.

26. Sun Y, Fan XY, Cao DM, Tang W, He K, Zhu JY, et al. Integration of Brassinosteroid Signal Transduction with the Transcription Network for Plant Growth Regulation in Arabidopsis. Dev Cell. 2010;19:765–77.
27. Qian W, Wu C, Fu Y, Hu G, He Z, Liu W. Novel rice mutants overexpressing the brassinosteroid catabolic gene CYP734A4. Plant Mol Biol. 2017;93:197–208.
28. Yin W, Chu C. Diversification of Plant Agronomic Traits by Genome Editing of Brassinosteroid Signaling Family Genes in Rice Towards Understanding of Molecular Basis of Abiotic Stresses in Plants View project Molecular mechanisms of nitrogen utilization efficiency in rice View project. Available from: https://academic.oup.com/plphys/advance-article/doi/10.1093/plphys/kiab394/6353036
29. Ito Y, Thirumurugan T, Serizawa A, Hiratsu K, Ohme-Takagi M, Kurata N. Aberrant vegetative and reproductive development by overexpression and lethality by silencing of OsHAP3E in rice. Plant Science. 2011;181:105–10.
30. Lee J, Koh HJ. A label-free quantitative shotgun proteomics analysis of rice grain development. Proteome Sci. 2011;9.
31. Hasanuzzaman M, Bhuyan MHMB, Parvin K, Bhuiyan TF, Anee TI, Nahar K, et al. Regulation of ros metabolism in plants under environmental stress: A review of recent experimental evidence. Int J Mol Sci. MDPI AG; 2020. p. 1–44.
32. Galland M, He D, Lounifi I, Arc E, Clément G, Balzergue S, et al. An integrated "multi-omics" comparison of embryo and endosperm tissue-specific features and their impact on rice seed quality. Front Plant Sci. 2017;8.
33. Liao JL, Zhou HW, Peng Q, Zhong PA, Zhang HY, He C, et al. Transcriptome changes in rice (Oryza sativa L.) in response to high night temperature stress at the early milky stage. BMC Genomics. 2015;16.
34. Zhang Y, Fernie AR. On the role of the tricarboxylic acid cycle in plant productivity. J Integr Plant Biol. Blackwell Publishing Ltd; 2018. p. 1199–216.
35. Meng X, Baine JM, Yan T, Wang S. Comprehensive Analysis of Lysine Lactylation in Rice (Oryza sativa) Grains. J Agric Food Chem. 2021;69:8287–97.
36. Chen F, Dong G, Wu L, Wang F, Yang X, Ma X, et al. A Nucleus-Encoded Chloroplast Protein YL1 Is Involved in Chloroplast Development and Efficient Biogenesis of Chloroplast ATP Synthase in Rice. Sci Rep. 2016;6.
37. Kusano H, Arisu Y, Nakajima J, Yaeshima M, She KC, Shimada H. Implications of the gene for F1-ATPase β subunit (AtpB) for the grain quality of rice matured in a high-temperature environment. Plant Biotechnology. 2016;33:169–75.
38. Muench SP, Trinick J, Harrison MA. Structural divergence of the rotary ATPases. Q Rev Biophys. 2011;44:311–56.
39. Timabud T, Yin X, Pongdontri P, Komatsu S. Gel-free/label-free proteomic analysis of developing rice grains under heat stress. J Proteomics. 2016;133:1–19.
40. Zhao H, Li Z, Amjad H, Zhong G, Khan MU, Zhang Z, et al. Proteomic analysis reveals a role of ADP-glucose pyrophosphorylase in the asynchronous filling of rice superior and inferior spikelets. Protein Expr Purif. 2021;183.
